# Examining the role of a chromosomal inversion in accumulating adaptive and barrier loci in a cold-adapted insect species

**DOI:** 10.1101/2025.06.16.659880

**Authors:** N. Poikela, V. Hoikkala, M. G. Ritchie, M. Kankare

## Abstract

Chromosomal inversions can play a crucial role in population adaptation and divergence by reducing gene flow and preserving adaptive allelic combinations between populations with different arrangements. However, demonstrating this empirically is challenging due to numerous interacting processes with similar genomic signatures affecting inversion evolution. In this study, we characterized a large (9.5Mb) polymorphic inversion in the cold-adapted and widely distributed species, *Drosophila montana*, using long- and short-read sequencing across several populations. The origin of this inversion predates the divergence of North American (NA) and Fennoscandian (North European) populations, suggesting it emerged in the ancestral *D. montana* population in the Rocky Mountains of NA. Despite the species’ expansion across the northern hemisphere, this inversion has remained exclusive to the Rocky Mountains populations, where it is fixed in the southernmost high-elevation population and appears at lower frequencies in the more northern and lower-elevation populations. By independently mapping SNPs linked to climate adaptation and barriers to gene flow (barrier loci; identified through reduced migration rates), we found enrichment of both within the inversion, with barrier and adaptive regions partially overlapping. However, the inversion was not enriched for SNPs related to cold tolerance. These findings suggest that inversions may maintain associations between multiple adaptive and barrier loci, effectively coupling them, and that locally adaptive regions may act as barriers to gene flow. Our study provides empirical evidence that inversions can contribute to population adaptation and divergence by reducing gene flow, maintaining adaptive allelic combinations, and facilitating the coupling of different barriers to gene flow.

**Significance:** Chromosomal inversions, genomic rearrangements with reversed gene order, are important drivers of population adaptation and divergence, but demonstrating this empirically is challenging due to complex interacting processes that produce similar genomic signatures. We characterized and analysed a large polymorphic inversion in *Drosophila montana*, a cold-adapted and widely distributed insect, using whole-genome sequencing data, phenotypic experiments, and recent methodological developments. This inversion is found only in mountain populations and is enriched with SNPs potentially associated with climate adaptation, as well as regions that act as barriers to gene flow, with these regions partially overlapping. These results suggest that adaptive loci can act as barriers to gene flow and that inversions help maintain adaptive allelic combinations and associations between multiple barrier loci, thereby contributing to population adaptation and divergence.

## Introduction

Barrier loci are defined as any loci that reduce gene exchange between populations, and they typically contribute to assortative mating and reduced fitness of hybrids (Butlin and Smadja 2018; Dopman et al. 2024). A central goal in evolutionary biology is to understand how barrier loci evolve. For example, do they begin as loci that promote local adaptation and become barrier loci, or are local adaptation and barrier loci independent? Can they become coupled by features of the genome, such as chromosomal inversions? Recent advances in genome sequencing have accelerated the development of computational tools to detect signatures of local adaptation, gene flow, and chromosomal rearrangements, enhancing our understanding of the selective, demographic, and molecular mechanisms driving population divergence despite ongoing gene flow.

Inversions, genomic regions with reversed gene order relative to the standard form, may play a major role in local adaptation and population divergence both with and without gene flow (reviewed in Hoffmann and Rieseberg 2008; Jackson 2011; Faria et al. 2019; Berdan et al. 2023). Inversions can gain a fitness advantage by significantly reducing recombination between individuals with different arrangements (Sturtevant 1921; Dobzhansky 1940), which allows for the accumulation and preservation of alleles with positive epistatic interactions (Hoffmann and Rieseberg 2008) and co-adapted gene complexes (Kirkpatrick and Barton 2006). The reduction in recombination is expected to be highest at the breakpoints, and it often extends beyond the inverted sequence (Kulathinal et al. 2009; Stevison et al. 2011), while gene flux that occurs via double crossovers and gene conversion can homogenize the regions towards the centre of the inversion (Navarro et al. 1997). Inversions may also be directly under selection if the breakpoints modify reading frames of genes, or if the new gene position, order or orientation, or changes in chromatin structure alter the expression of the genes (Villoutreix et al. 2021; Wright and Schaeffer 2022; Berdan et al. 2023). If there are no counteracting forces, these mechanisms can drive an inversion to high frequency, potentially leading to fixed differences between populations (Hoffmann and Rieseberg 2008; Berdan et al. 2023). On the other hand, inversion polymorphism within a species can be maintained through some form of balancing selection, such as heterozygote advantage (overdominance), disassortative mating, or frequency-dependent, spatially or temporally varying selection. However, distinguishing the different mechanisms that act on inversions can be extremely difficult because multiple, sometimes interacting processes affect their evolution and because the different processes may result in similar genomic signatures (Berdan et al. 2023).

Showing empirically that alternatively fixed inversions between populations restrict gene flow and accumulate barrier loci remains challenging. Traditionally, outlier loci have been identified through summary statistics, such as genetic divergence (d_xy_) and genetic differentiation (F_st_). However, these statistics can be misleading due to their susceptibility to different demographic, selective, and genetic factors, potentially resulting in false positives (Cruickshank and Hahn 2014; Ravinet et al. 2017; Laetsch et al. 2023). This limitation has led to the development of new methods that better account for these confounding factors and focus on detecting reduced migration as a signal of barrier regions (Fraïsse et al. 2021; Laetsch et al. 2023; Burban et al. 2024). While some studies have shown that mean migration rate (*m_e_*) is lower within inverted compared to non-inverted (colinear) regions between populations connected by gene flow (e.g. Lohse et al. 2015; Poikela et al. 2024), studies have rarely investigated individual barrier loci and their association with rearrangements (Mackintosh et al. 2023). In light of recent studies, both inversions that have emerged at the onset of lineage divergence (Lohse et al. 2015) and inversions that predate lineage divergence and have existed already in their common ancestral population (Fuller et al. 2018; Poikela et al. 2024) can potentially drive divergence with gene flow.

To better understand the role of inversions in population divergence, it is essential to search for signs of local adaptation alongside barriers to gene flow. The presence of inversion frequency clines along environmental gradients, where balanced polymorphisms are maintained by spatially varying selection, was one of the first compelling pieces of evidence for the role of inversions in local adaptation (Dobzhansky and Levene 1948). Such clines have since been found in other systems (reviewed in Wellenreuther and Bernatchez 2018). Also, Quantitative Trait Loci (QTL) mapping and genotype-environment association (GEA) analysis have proven effective in evaluating the role of inversions in adaptation. For example, Ayala et al. (2019) found that single nucleotide polymorphisms (SNPs) linked to desiccation resistance were located within the 2La and 2Rb inversions in *Anopheles gambiae.* It is also crucial to investigate the precise overlap between barrier loci and adaptive loci, as this may reveal whether adaptive loci act as barriers to gene flow as a result of ecologically-based divergent selection (Schluter 2001; Rundle and Nosil 2005). Finally, inversions may contribute to “coupling”, where different barrier effects become linked through reduced recombination within inversions, ultimately promoting the overall barrier to gene flow (Butlin and Smadja 2018; Dopman et al. 2024).

The malt fly *Drosophila montana* is an ideal species for studying the role of chromosomal inversions in facilitating local adaptation and reducing gene flow between populations. It occupies a broad range of climates, including extreme cold environments, exhibits reproductive barriers between populations, and possesses inversion polymorphisms. Spanning the northern hemisphere, its distribution ranges from Fennoscandia (Northern Europe) to Far East Asia near the Bering Strait, and North America (NA), with southern extensions into the Rocky Mountains (from elevations of 1400m to above 3000 m) and along the continents’ western coast (Fig. 1A, Table S1; Patterson 1952; Throckmorton 1982; Hoikkala and Poikela 2022). The split between Fennoscandian and North American *D. montana* is estimated to have occurred ∼1.75mya, soon after which the Crested Butte (NA) and other North American populations diverged (Garlovsky et al. 2020). Populations from these different regions show partial reproductive isolation (Jennings, Snook, et al. 2014). Additionally, *D. montana*, which has the highest recorded cold tolerance among *Drosophila* species (Kellermann et al. 2012; MacMillan et al. 2015) and can overwinter as an adult for up to six months under snow cover (Hoikkala and Poikela 2022), shows latitudinal variation in cold tolerance across its distribution (Poikela et al. 2021; Wiberg et al. 2021). While previous polytene chromosome studies suggest inversion polymorphism in *D. montana* (Stone et al. 1960; Throckmorton 1982), these inversions have not yet been characterized and studied at the genomic level.

**Figure 1.**
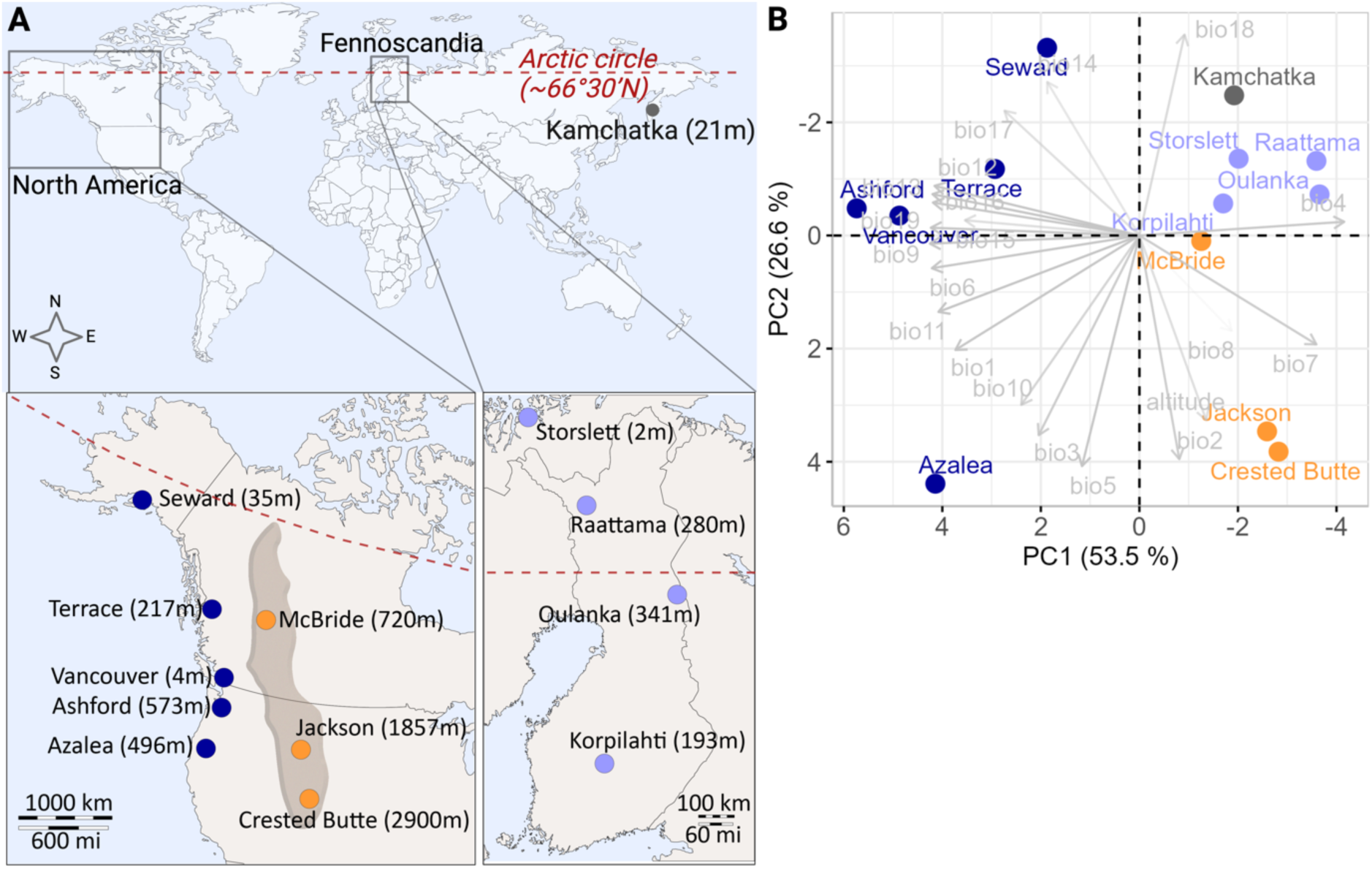
(A) *D. montana* sampling sites in the western coast and in the Rocky Mountains of North America, in Fennoscandia, and in Kamchatka (Asia). The Arctic circle is marked with a dashed line. (B) Climatic conditions across *D. montana* sampling sites summarized in a PCA using information on 19 bioclimatic variables (Table S2) of each fly collecting site.

In this study, we use long- and short-read whole-genome sequencing (WGS) data, along with climate and cold tolerance data, GEA, and recent developments in demographic modelling, to characterize polymorphic inversions and investigate their role in population adaptation and divergence. First, we examine variation in climate conditions and cold tolerance and characterize polymorphic inversions and their frequencies across the species’ distribution range. Next, we explore whether a large polymorphic inversion on chromosome 4 is enriched for loci associated with climate and cold adaptation. Finally, we investigate the origin of this inversion, assess its enrichment for barriers to gene flow, and determine whether barrier loci overlap with adaptive loci. Our results provide empirical evidence that inversions have promoted population adaptation and divergence by reducing gene flow, preserving adaptive allelic combinations, and facilitating the coupling of barriers to gene flow.

## Result and Discussion

### *D. montana* flies live in diverse geographic and climatic conditions

*D. montana* samples were collected from 13 sites in the western coast and Rocky Mountains of North America (NA), Fennoscandia, and Kamchatka (Fig. 1A; Table S1). To characterize the climatic environments each population has adapted to, we conducted a principal component analysis (PCA) based on 19 bioclimatic variables of the geographic coordinates of each sampling site (Fig. 1B; Table S2). Three PCs had Eigenvalues >1, but the first two had the highest contribution and explained more than 81% of the total variation (Fig. 1B; Table S3). PC1 separated the western coast populations from the others (Fig. 1B). The coastal populations have warmer annual and winter temperatures (bio1, bio6, bio9 and bio11), lower seasonal variation in temperatures (bio4 and bio7) and receive higher precipitation throughout the year (bio12-13, bio15-16, bio19) compared to the other populations (Fig. 1B, Table S2, Table S4). PC2 separated the southern and northern coastal populations, as well as the Rocky Mountains populations from the Fennoscandian and Kamchatkan populations (Fig. 1B). These populations differed in daily temperature changes (bio2), isothermality (bio3, the ratio of day-to-night temperature variation to summer-to-winter variation) and variation in temperature and precipitation during warm months (bio5 and bio18; Table S4). In conclusion, *D. montana* has adapted to highly variable climates in its circumboreal distribution.

### *D. montana* cold tolerance traits show latitudinal variation and are partially correlated

An ability to cope with cold temperatures is one of the most important adaptations to high latitudes and altitudes. To examine clinal variation in cold tolerance, we studied cold tolerance traits in *D. montana* across its distribution range using nine different populations (Fig. 2, Table S1). Cold tolerance was assessed using two ecologically relevant and widely used methods: critical thermal minimum (CT_min_) and chill-coma recovery time (CCRT). CT_min_ represents the lowest temperature the fly can withstand without falling into chill-coma, while CCRT measures the time required for flies to recover from a chill-coma. Lower values in both tests indicate better cold tolerance.

**Figure 2.**
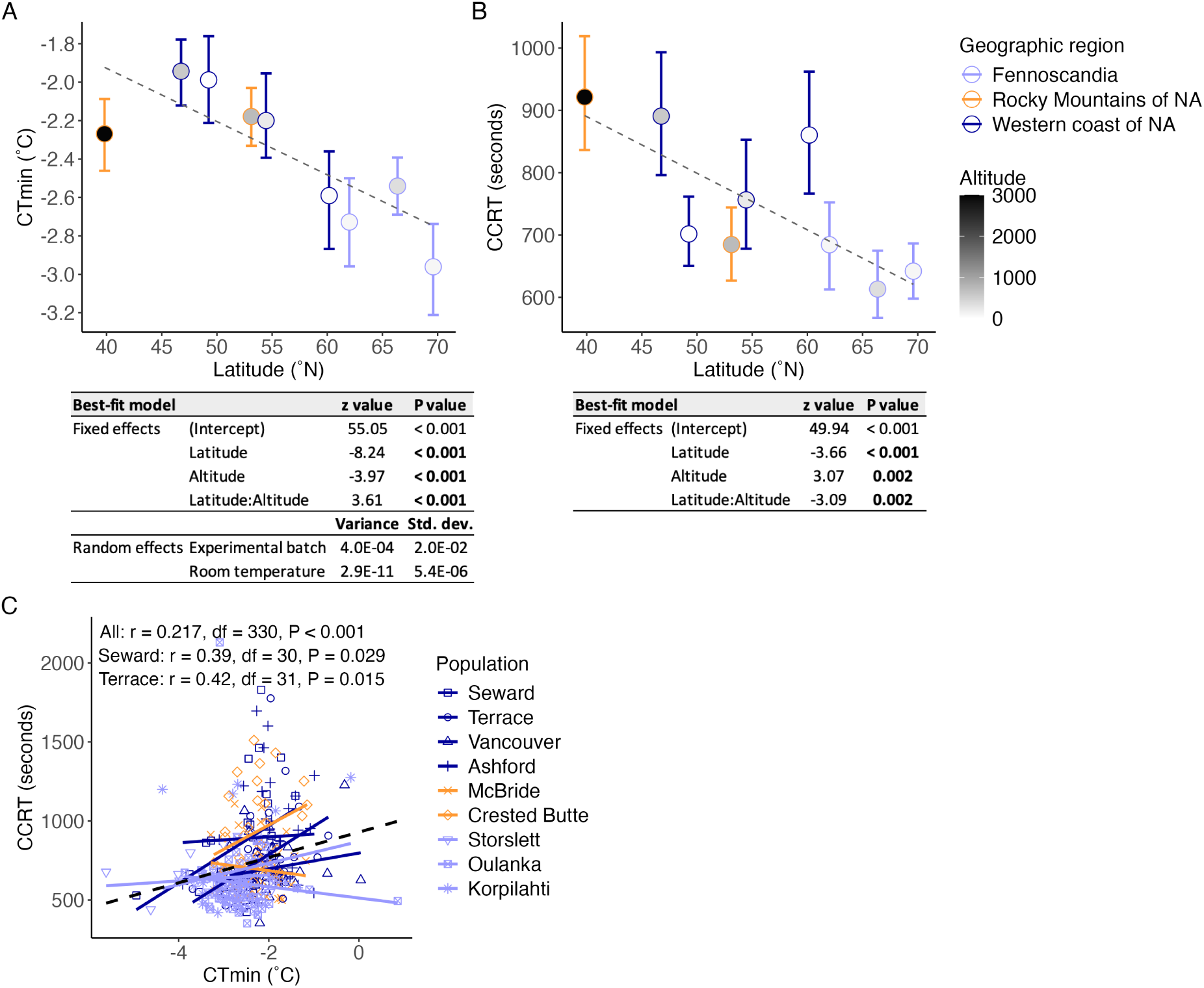
*D. montana* cold tolerance measured as (A) chill coma temperature (CT_min_) and (B) chill coma recovery time (CCRT) across latitudes and altitudes. The points and error bars show the means and standard errors, respectively. Dashed lines indicate the predicted values from the partial effect of latitude. The best-fit models are shown below the plots (model comparison in Table S5). (C) Pearson correlation and regression lines between CT_min_ and CCRT across all nine populations (black dashed line), and separately for each population (all statistics and separate plots for populations are given in Table S6 and Fig. S1, respectively).

The best-fit model for the traits was chosen based on Akaike’s Information Criterion (AIC; Table S5). The best-fit model for CT_min_ included latitude, altitude and their interaction as fixed effects and experimental batch and room temperature during the experiment as random effects (Fig. 2A; Table S5). The best-fit model for CCRT included latitude, altitude and their interaction as fixed effects and no random effects (Fig. 2B; Table S5).

Flies’ ability to resist cold temperatures (CT_min_) improves towards northern (higher/colder) latitudes, and the positive interaction term between latitude and altitude suggests that the effect of latitude on CT_min_ becomes more pronounced as altitude increases (Fig. 2A). In other words, flies’ ability to resist cold temperatures improves towards northern latitudes and higher altitudes. Flies’ ability to recover from chill coma (CCRT) also improves towards northern (higher/colder) latitudes, but the negative interaction term suggests that the relationship between latitude and CCRT becomes weaker as altitude increases (Fig. 2B). However, the significant interaction terms should be interpreted with caution since they likely derive from one outlier population (Crested Butte) with an altitude of 2900m. Finally, we found a weak positive correlation between CT_min_ and CCRT across all studied individuals (both measures were taken for the same individuals, CT_min_ first) (r = 0.22, P < 0.001; Fig. 2C, Table S6). However, when analysing the data separately for each population, the correlation was found to be population-dependent and significant only in the northern western coast populations (Seward: r = 0.39, P = 0.029, Terrace: r = 0.42, P = 0.015; Fig. 2C, Fig. S1; Table S6).

Generally, populations living at higher latitudes or altitudes experience colder environments than those at lower latitudes or altitudes, which is expected to lead to genetic differences and clinal variation in cold tolerance traits (Addo-Bediako et al. 2000). The clinal variation observed in *D. montana* supports previous findings in this species (Wiberg et al. 2021), and contributes to the expanding evidence for clinal variation in cold tolerance across ectotherms (e.g. Hoffmann et al. 2002; Overgaard et al. 2011; Zuther et al. 2012). While the discovery of clinal variation in cold tolerance traits is expected, the partial correlation between CT_min_ and CCRT has not been observed before in *D. montana* (Wiberg et al. 2021) or other species (e.g. Overgaard et al. 2011; Andersen et al. 2015). This partial correlation suggests that the resistance to chill coma (CT_min_) and the recovery from it (CCRT) may either share mechanisms or coevolve in specific populations or environments, but not universally.

### The Rocky Mountains populations harbour a large polymorphic inversion on chromosome 4

We characterized polymorphic inversions in *D. montana* using PacBio and Illumina re-sequence data and genome assemblies (Table S7-S10). We found one 9.5Mb polymorphic inversion located on chromosome 4 (Table S11, Fig. S2). This inversion was found only in the Rocky Mountains populations, being fixed in Crested Butte (100%) and present at lower frequencies in Jackson (17%) and McBride (22%) (Fig. 3, Table S11), though the inversion frequencies in Jackson and McBride are less reliable due to the low sample sizes (N = 3 and N = 9, respectively, Table S11). Overall, the inversion contains ∼850 genes; those closest to the proximal breakpoint include *tRNA-specific adenosine deaminase* and *UPF0428 protein CG16865*, while those near the distal breakpoint include *sorting nexin-17* and *tRNA-dihydrouridine(20a/20b) synthase [NAD(P)+]-like* (see the exact locations in Fig. S2).

**Figure 3.**
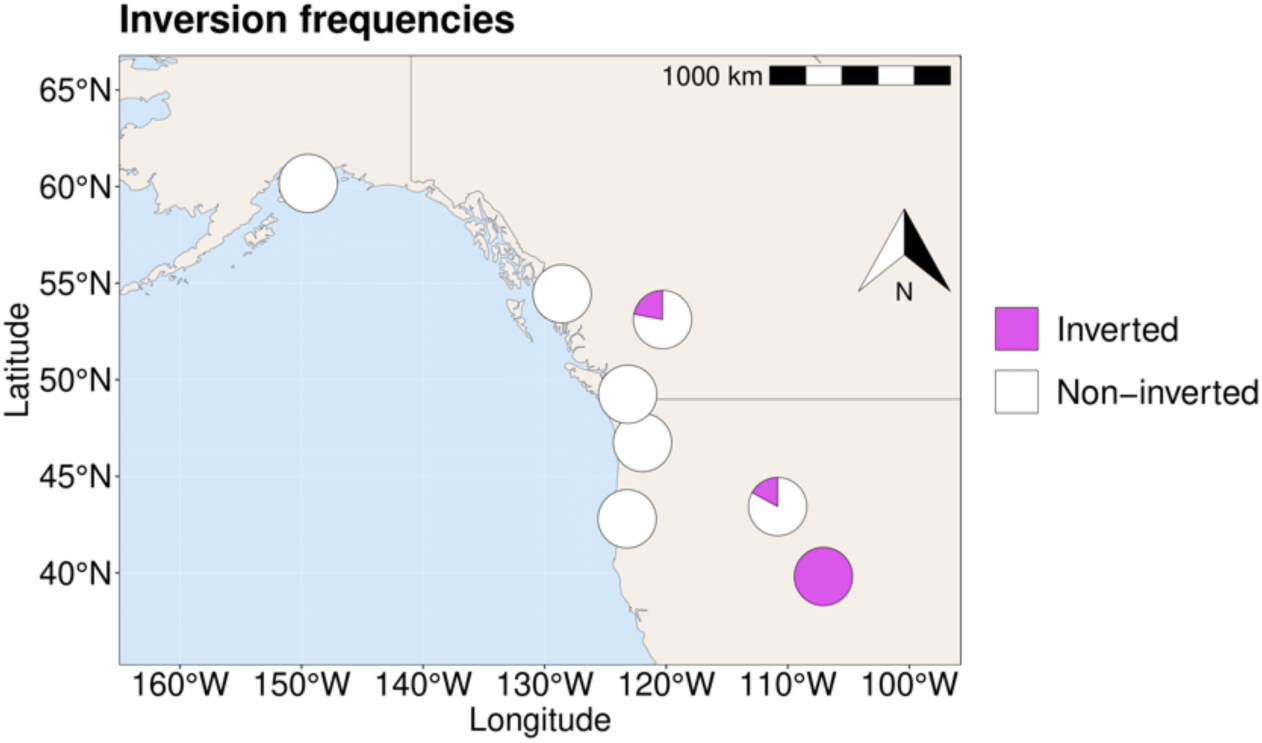
Frequencies of the large polymorphic inversion (9.5Mb) on chromosome 4. The inversion is fixed in the southernmost population of the Rocky Mountains (Crested Butte) and occurs at lower frequencies in the more northern Rocky Mountains populations (Jackson, McBride). It is absent in all other studied North American, Asian and Fennoscandian populations (see Table S11 for details).

We also found a small inversion of approximately 1500bp at the proximal end of chromosome 4 (Table S12, Fig. S2). This inversion was present only in the Fennoscandian and Kamchatkan samples and is located in the intronic region of a gene coding for a leucine-rich repeat transmembrane neuronal protein 3. Given its small size and the synonymous effects, which most likely do not impact the protein product, it was not investigated further. Older polytene chromosome studies (Stone et al. 1960; Throckmorton 1982) suggest that *D. montana* may harbor additional polymorphic inversions not identified here, possibly due to the limited number of strains sequenced.

Tahami et al. (2024) found Gypsy and MITE transposable elements (TEs) at the breakpoints of the large chromosome 4 inversion fixed in Crested Butte, which are absent in populations without the inversion. Similarly, the small chromosome 4 inversion in Fennoscandian populations contains unique Helitron and Gypsy TE insertions. These findings suggest that TE insertions near breakpoints may have facilitated the origin of the inversions through ectopic recombination, although further research is needed to confirm this. We explore the role of the large chromosome 4 inversion in local adaptation and population divergence in the following sections.

### Geography and the polymorphic chromosome 4 inversion contribute to the population genetic structure of *D. montana*

To summarize the population genetic structure of *D. montana* and the role of geography and the polymorphic inversion therein, we conducted separate PCAs for the whole genome and for the inversion using the Illumina re-sequence data of 101 wild-caught females (Fig. 1A, Table S1, S8). The first PCA included 1,986,534 filtered SNPs across the genome. The first three principal components (PCs) explained 35%, 22% and 6% of the genetic variation (Fig. 4A; Table S13). PC1 and PC2 revealed four major clusters corresponding to Crested Butte and other North American (NA) populations, Fennoscandian populations, and Kamchatka (Asia). PC3 showed a latitudinal cline within North American populations, which was primarily due to genetic variation on chromosome 4 (Fig. S3E). The second PCA included 166,193 filtered SNPs from the chromosome 4 inversion, revealing similar patterns to the genome-wide PCA (Fig. 4B, Table S13), with some key differences. Here, Jackson and McBride samples that carry the inversion in either homozygous or heterozygous states clustered more closely with Crested Butte, where the inversion is fixed (Fig. 4B, Table S11). Notably, a homozygous inversion carrier from McBride clustered closest to Crested Butte, while heterozygous inversion carriers were more distant. This pattern was not observed on colinear regions on chromosome 4 (Fig. S3A, Table S13). Hence, genetic variants in the inversion are maintained across populations.

**Figure 4.**
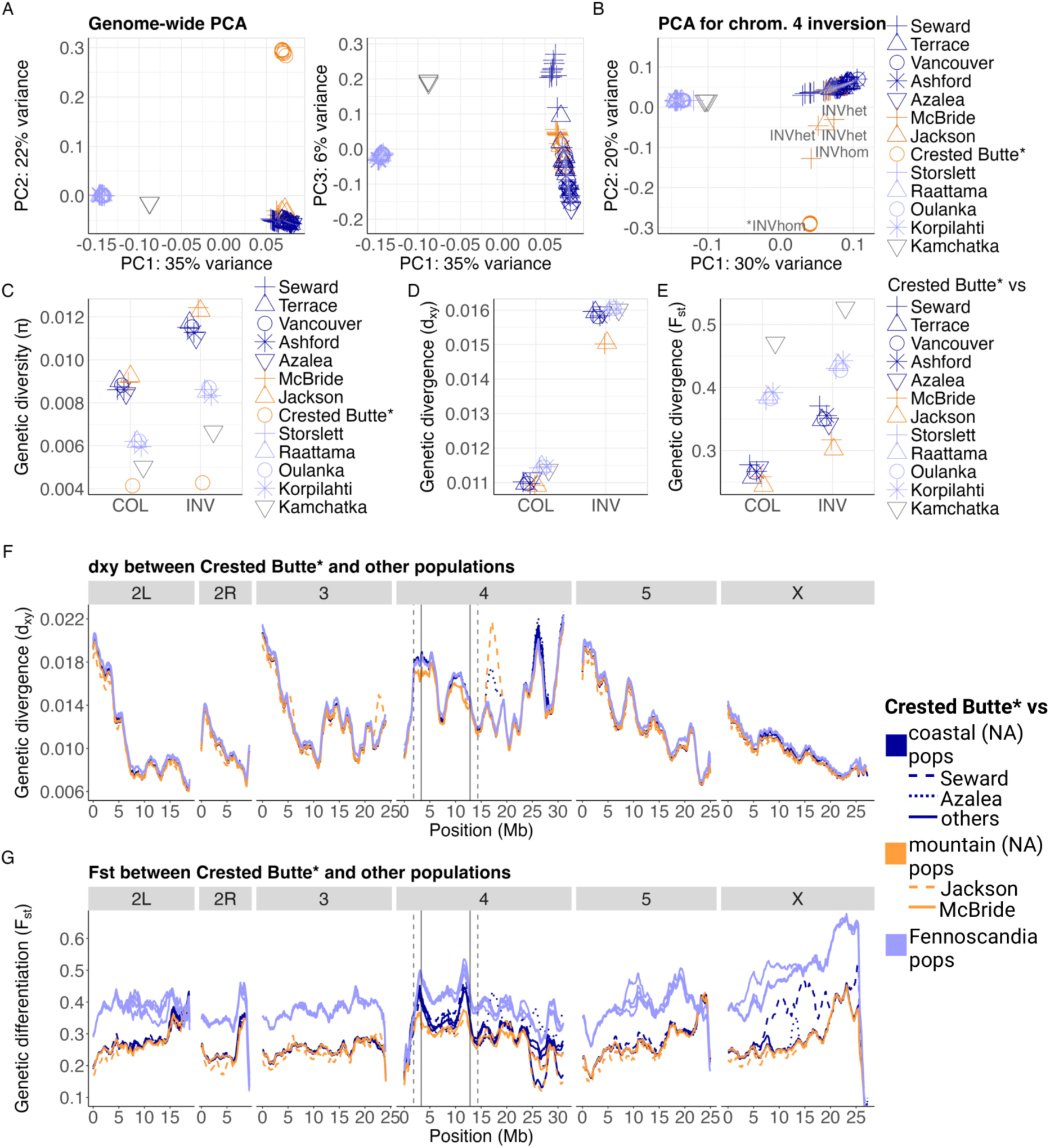
Genetic population structure of *D. montana*. Genetic variation was summarized in principal component analyses (PCA) including SNPs (A) from all chromosomes and (B) from the large inversion on chromosome 4. The inversion is fixed in Crested Butte (marked with *), i.e. all samples are homozygous for the inversion (*INVhom). The Jackson and McBride samples that carry the inversion at homozygous (INVhom) or heterozygous (INVhet) state are marked separately; see Table S11). (C) Mean genetic diversity (π) was calculated for each population and for inverted (INV) and non-inverted (colinear, COL) regions. Mean (D) genetic divergence (d_xy_) and (E) genetic differentiation (F_st_) were calculated between Crested Butte (inversion fixed) and other populations (inversion at lower frequencies in Jackson and McBride and absent in the other populations) for INV and COL regions. (F) d_xy_ and (G) F_st_ were plotted in sliding windows of a fixed block size between Crested Butte and the other populations (125 blocks ξ blocks of 256bp in length, with a step change of 25 blocks ξ blocks of 256bp in length). Populations are highlighted with dashed lines whenever their divergence strongly deviates from that of the other populations of the same geographic region. The inversion breakpoints are marked with vertical solid lines. Since recombination is often suppressed beyond the inverted region into the colinear region, 1.5Mb buffer region was marked at both ends of the inversion with vertical dashed lines. Kamchatka samples were excluded from the d_xy_ and F_st_ window-wise plots since they were collected from inbred laboratory strains.

We also measured genetic diversity (π) for each population, and divergence (d_xy_) and differentiation (F_st_) between populations (Fig. 4C-E, Table S14-S15, Fig. S4-S6). North American populations showed ∼40% higher π than Fennoscandian populations, except for Crested Butte (NA) which had the lowest diversity of all samples. Kamchatka had similarly low diversity, although this may be unreliable because these samples were obtained from isofemale strains, not directly from the wild. π was particularly high in the inverted region of Jackson and McBride, reflecting their mixed inverted and non-inverted haplotypes (Fig. 4B-C, Table S15). d_xy_ and F_st_ generally increased with geographic distance, following an isolation-by-distance model, except for Kamchatka and Crested Butte. d_xy_ and F_st_ were highest between North American and Fennoscandian populations, lower among North American populations, and lowest within Fennoscandia. Despite its geographic proximity to North America, Kamchatka was genetically closer to Fennoscandia, likely due to the isolation by the Bering Strait. Crested Butte exhibited the highest genetic divergence from other populations, regardless of geographic location, indicating that it is a relatively isolated population. Finally, d_xy_ and F_st_ between Crested Butte, where the inversion is fixed, and populations without the inversion were higher in the inverted regions compared to the colinear regions (Fig. 4D-G, Table S15). Although d_xy_ and F_st_ were also elevated in the inverted regions between Crested Butte and the Rocky Mountains populations (Jackson and McBride), the values were not as high due to the presence of the inversion at lower frequencies in Jackson and McBride (Fig. 4D-G, Table S15).

Taken together, genetic variation in Crested Butte, other North American populations, Fennoscandia, and Kamchatka contribute to large-scale genetic variation across *D. montana*’s range, while the chromosome 4 inversion is one of the key drivers of finer-scale genetic variation in *D. montana*. The phylogeographic history of the species has been uncertain, with hypotheses suggesting either a North American origin, followed by a spread to Asia and Europe, or an Asian origin, with subsequent spread to North America and Europe (Throckmorton 1982; Mirol et al. 2007; Garlovsky et al. 2020). Given the highest genetic diversity observed in North America, which gradually decreases towards Alaska (NA) and Northern Europe (Fennoscandia) (Table S14-S15), along with the presence of all *montana* complex species in North America (Throckmorton 1982; Hoikkala and Poikela 2022), it is likely that *D. montana* originated in North America and spread into Asia and Europe from there. The low genetic diversity in Crested Butte (NA) may indicate a founder effect or population bottleneck, which could explain the fixation of the inversion in that population, while not in Jackson or McBride (NA). This inversion may still play a key role in local adaptation, for example, by reducing recombination and gene flow between populations, as indicated by the increased genetic divergence within the inversion, a topic we explore in the following sections.

### SNPs associated with climate adaptation are enriched within the inversion

To explore the role of the chromosome 4 inversion in accumulating and preserving loci under local adaptation, we linked genomic data to climatic conditions and experimental cold tolerance data. This was achieved through a genotype-environment association (GEA) analysis using Illumina data from nine populations (Table S8), combined with population-specific environmental variables (PC1, PC2; Fig. 1B) and cold tolerance traits (CT_min_, CCRT; Fig. 2A-B). The inversion is fixed in the Crested Butte population and found at lower frequency in the McBride population, but absent in the other North American and all Fennoscandian populations. The recombination-suppressing effects of inversions can extend up to 2-3Mb beyond the inverted sequence into the colinear regions (Kulathinal et al. 2009; Stevison et al. 2011). Therefore, in addition to exploring SNPs strictly within the inversion breakpoints, we extended the inversion by a conservative estimate of 1.5Mb at both ends since there is no prior knowledge on the effect of inversions on recombination in *D. montana*.

Across different genomic regions, only 0.18-5.01% of SNPs were significantly associated with environmental variables (PC1, PC2) or cold tolerance phenotypes (CT_min_, CCRT; Fig. 5, Fig. S7, Table S16). The inversion exhibited a significantly higher enrichment of SNPs associated with PC1 and PC2 compared to colinear regions, and this enrichment persisted regardless of whether the regions analyzed were limited to the inversion breakpoints (Chi Square, PC1: X^2^=6.5, P=0.011, PC2: X^2^=2079.2, P<0.001) or extended 1.5Mb beyond them into the colinear regions (PC1: X^2^=500.7, P<0.001, PC2: X^2^=4754.8, P<0.001; Fig. 5A, Fig. S7A-B, Table S17).

**Figure 5.**
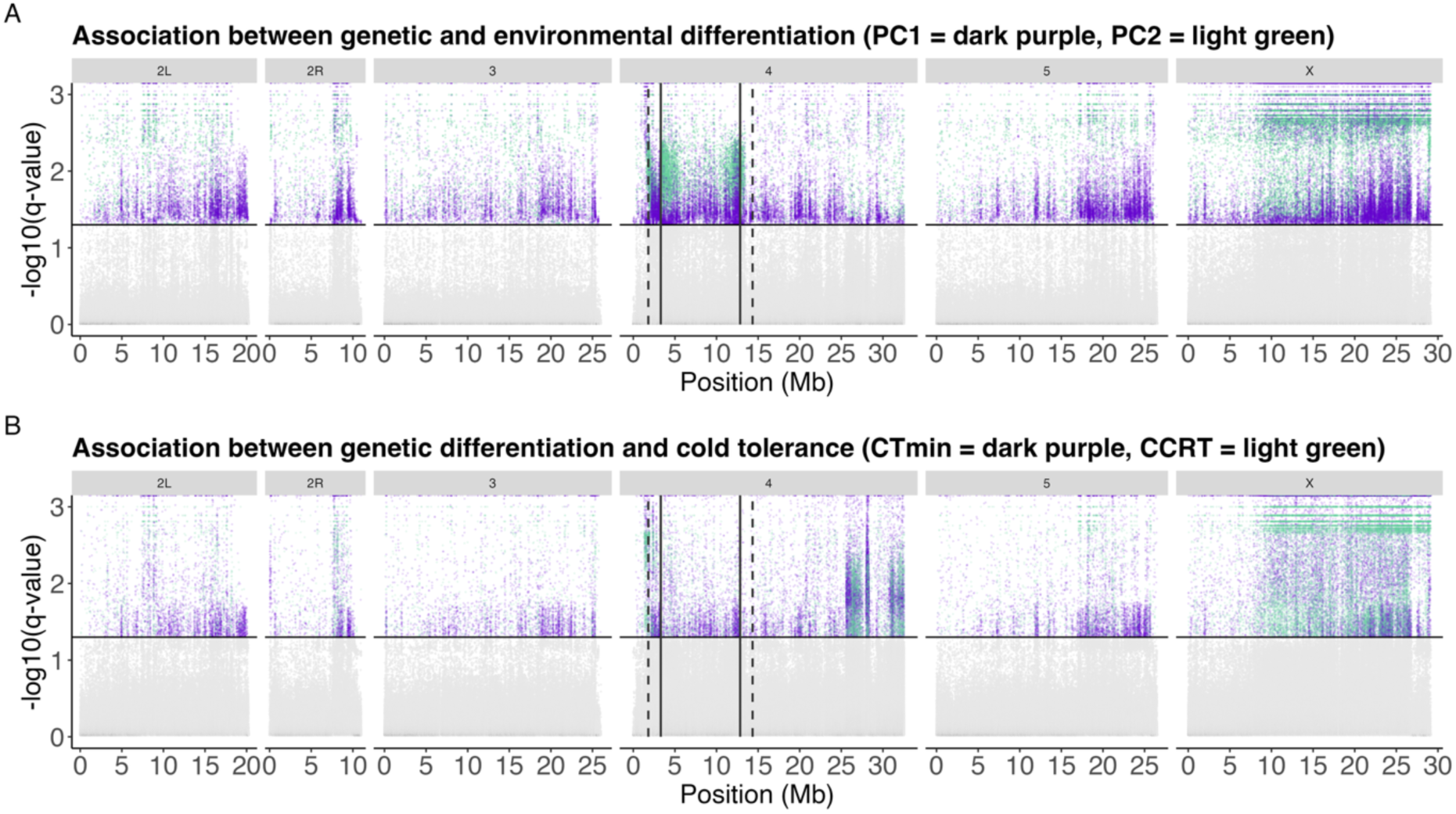
Genotype-environment association analysis (GEA; BayeScEnv). Points above the horizontal line indicate significant SNPs, describing significant association between (A) genetic and environmental (PC1=dark purple, PC2=light green) differentiation and (B) genetic differentiation and cold tolerance (CT_min_= dark purple, CCRT=light green; see separate plots in Fig. S7). q-values for the g parameter of two chains (Fig. S8-S11) were used to control the false positive rate at 0.05, i.e. SNPs with q-values>1.3, to obtain the final candidate SNPs. The solid vertical lines represent the inversion breakpoints, while the dashed vertical lines mark a conservative estimate of 1.5Mb beyond the breakpoints, where recombination is expected to remain reduced.

In contrast, the inversion did not exhibit a significantly higher enrichment of SNPs associated with CT_min_ and CCRT (Fig. 5B, Fig. S7C-D, Table S17). Instead, SNPs associated with CT_min_ and CCRT showed two narrow peaks near the telomeric end of chromosome 4 (Fig. 5B, Fig. S7C-D, Table S17). Additionally, the X chromosome displayed a strong enrichment of SNPs associated with all four variables (Fig. 5, Fig. S7, Table S16). The X chromosome showed some horizontal banding patterns in q-values, a computational artifact where multiple SNPs are assigned identical significance values, so these particular results should be interpreted with caution.

The significant SNPs inside the inversion were distributed across 146-507 genes, depending on the analyzed dataset (Table S18). Based on the gene ontology (GO) enrichment analysis, the genes linked to the four variables were significantly associated e.g. with epidermal growth factor-like domain, immunoglobulin (Ig)-like domain, calcium transport, and membrane and transmembrane functions (Table S18, Supplementary file 1). Many of these groups have been repeatedly linked to the ability of *D. montana* to withstand cold temperatures (Kauranen et al. 2019; Parker et al. 2021; Wiberg et al. 2021), and genes connected to membrane, transmembrane and immunoglobulins were among the fastest evolving when comparing several cold-tolerant *Drosophila* species for their rates of molecular evolution (Parker et al. 2018). Membrane proteins play a crucial role in membrane and cuticular functions, which are especially important in cold temperatures and desiccation tolerance (Gibbs 2002; Stinziano et al. 2015). Additionally, cell functioning at low temperatures is strongly dependent on changes in ion transport and fluidity (Hazel 1995; Teets and Denlinger 2013).

Overall, the accumulation of SNPs linked to climate adaptation within the inversion and near its breakpoints (Fig. 5), along with its exclusive presence in the Rocky Mountains populations (Fig. 3), suggest that the inversion may contribute to local adaptation in the Rocky Mountains. However, it remains unclear whether this local adaptation has driven the fixation of the inversion in the southern high-altitude Crested Butte population, or if the fixation is due to other factors not related to selection, such as a population bottleneck or founder effect. In contrast, the inversion is found in a polymorphic state in the more northern, lower-altitude Rocky Mountains populations (Jackson and McBride), possibly because it has not had time to reach fixation, or is maintained polymorphic through some form of balancing selection, such as heterozygote advantage, frequency-dependent selection, or environmental heterogeneity (Wellenreuther and Bernatchez 2018), but we lack direct fitness estimates of the genotypes. An intriguing question also remains as to why the inversion has not spread to Fennoscandia, despite the Fennoscandian climate resembling that of the Rocky Mountains (Fig. 1B), where the inversion could potentially be beneficial. Perhaps geographical or environmental barriers have prevented its spread to Fennoscandia, or the inversion is linked to other environmental factors or phenotypic traits not tested in this study. Future research could focus on comparing both phenotypic traits and gene expression profiles between different inversion genotypes (inverted homozygous, non-inverted homozygous and heterozygous) under stressful environments, to identify the potential drivers of the inversion evolution.

### The inversion is old and enriched with barrier loci

To investigate the evolutionary history of *D. montana* and the role of the chromosome 4 inversion in population divergence, we used a demographic modelling approach (gIMble; Laetsch et al. 2023). First, we compared and identified the best-fit demographic model between three genetically, climatically, and geographically isolated populations: Crested Butte (North America, inversion fixed), Vancouver (North America, inversion absent), and Storslett (Fennoscandia, inversion absent; Fig. 1, Fig. 3). We tested models both without (strict divergence, DIV) and with post-divergence gene flow in both directions (isolation with migration, IM). This initial model selection focused on colinear regions, as the evolutionary history of inversions may differ from other genomic regions. The best-fit demographic scenario for Storslett-Crested Butte and Vancouver-Crested Butte comparisons suggests post-divergence gene flow (IM model), with gene flow occurring from Storslett and Vancouver to Crested Butte (forward in time; Table S19). Our parametric bootstrap analysis showed that the IM model significantly improved the fit compared to the DIV model in these comparisons, indicating a genuine signal of post-divergence gene flow (Fig. S12). However, while the IM model had higher support than the DIV model in the Storslett-Vancouver comparison (Table S19), the parametric bootstrap analysis showed that the improvement in fit was not significant (see Fig. S12). Consequently, the DIV model could not be rejected for this comparison. Although *D. montana* likely originated in North America and later spread to Asia and Europe (as discussed earlier), post-divergence gene flow can occur in multiple directions, reflecting more recent gene exchange rather than the original migration patterns. The similar climate conditions in Northern Europe (Storslett) and the Rocky Mountains of North America (Crested Butte; see Fig. 1B) may explain the gene flow from Northern Europe to the Rocky Mountains. Conversely, the lack of gene flow from Storslett to Vancouver could be due to climate differences, as Vancouver is a low-latitude, low-altitude population (Fig. 1B). This is also supported by the observation that in Europe this species has not spread to southern locations.

Second, the best-fit model of each population pair (Table S19) was used to analyse the mean time of divergence (*T*) and migration rate (*m_e_*) separately for inverted and colinear chromosomal regions to estimate the age of the inversion and its effects on migration rate (Fig. 6A-B, Table 1). Based on the colinear background, the North American (Crested Butte, Vancouver) and Fennoscandian (Storslett) populations of *D. montana* diverged approximately 700,000 to 1,000,000 years ago (Fig. 6A, Table 1). The estimated divergence time for the inverted region was approximately 1.4 million years between the Crested Butte population, where the inversion is fixed, and the Vancouver and Storslett populations, where the inversion is absent (Fig. 6A, Table 1), suggesting that the inversion was already present in the ancestral population before *D. montana* populations began to diverge. Notably, the colinear and inverted regions between Vancouver and Storslett − neither of which carry the inversion − diverged around the same time (Fig. 6A, Table 1). Additionally, the mean migration rate estimate was significantly lower within the inverted compared to colinear regions between Crested Butte and Vancouver, where gene flow has been strongest and may still be ongoing (Fig. 6B, Table 1). Migration rate estimates were very low for both inverted and colinear regions between Crested Butte and Storslett (Fig. 6B, Table 1), likely due to strong geographic isolation reducing gene flow across the genome, not solely within the inversion. Since the strict divergence (DIV) scenario between Vancouver and Storslett could not be rejected, *m_e_* was estimated to be zero for both inverted and colinear regions (Fig. 6B, Table 1). Overall, the inversion likely has not played a major role in the divergence between North American and Fennoscandian populations, as none of the Fennoscandian populations and most North American populations do not carry the inversion, and the strong geographic isolation has reduced gene flow across the entire genome.

**Figure 6.**
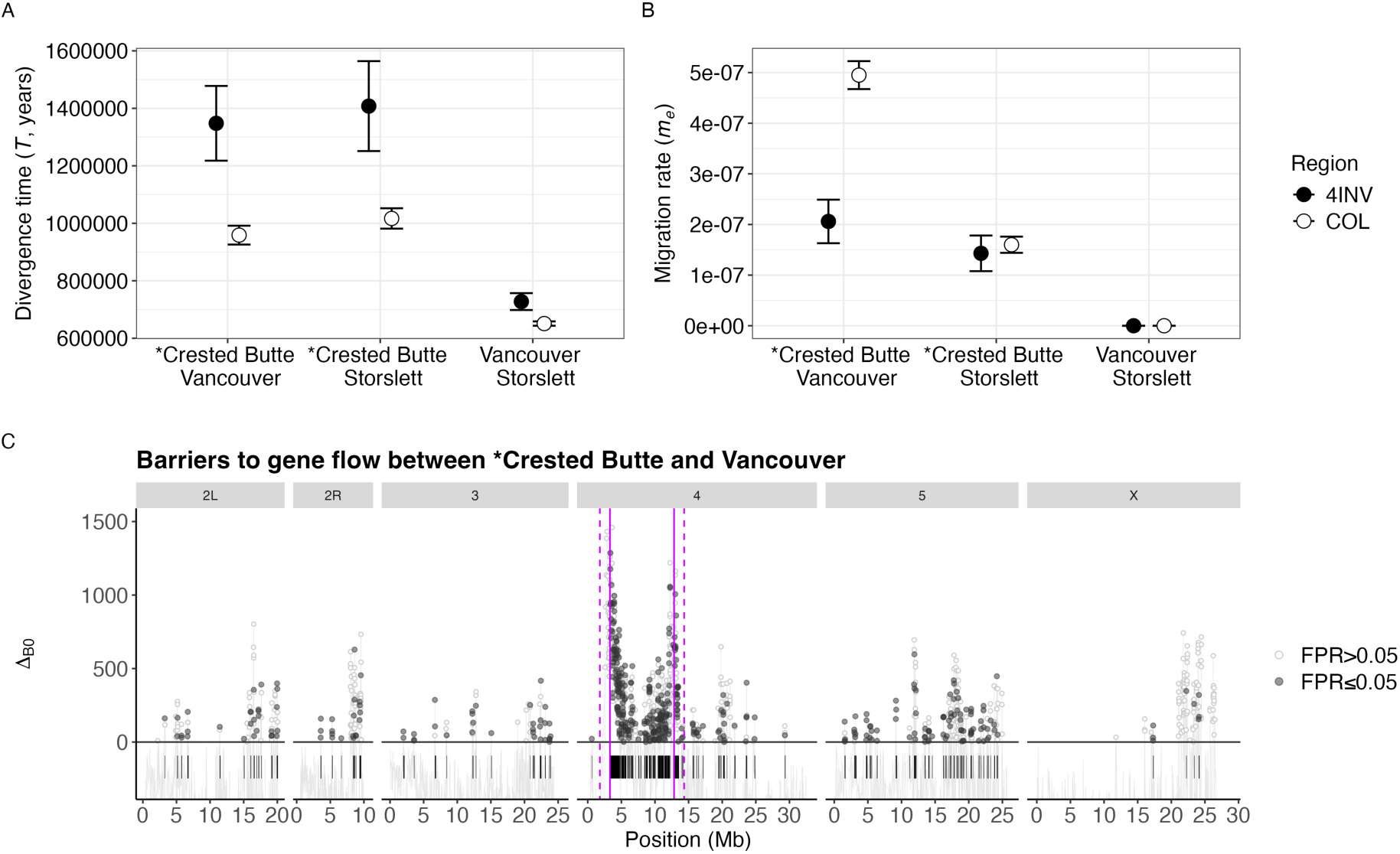
Evolutionary history of *D. montana* populations and the polymorphic inversion on chromosome 4. (A) Mean divergence time (*T*) and (B) mean migration rate (*m_e_*) of inverted (4INV) and colinear (COL) regions between three *D. montana* populations. The *m_e_* estimates were inferred under the best-fit IM model, except for Vancouver-Storslett comparison where the *m_e_* estimates were based on the DIV model that assumes no post-divergence gene flow (*m_e_*=0; see Table 1). Crested Butte, the population with the fixed inversion, is highlighted with * (absent in the other populations). Confidence intervals were estimated using the parametric bootstrap as +/- 2 SD across the 100 replicates under the best-fit models with recombination (see Methods). (C) Support for locally reduced *m_e_* between Crested Butte and Vancouver was measured via positive Δ_B0_ (data points above the dashed horizontal line) and a false positive rate at 5% (more detailed plot in Fig. S13). Barrier regions (Δ_B0_ > 0 and FPR ≤ 0.05) are marked with black segments. The solid vertical lines represent the inversion breakpoints, while the dashed vertical lines mark a conservative estimate of 1.5Mb beyond the breakpoints, where recombination is expected to remain reduced.

**Table 1.** Parameter for effective population sizes (N_e_), divergence times (*T* in years/generations) and migration rates (*m_e_*) for three *D. montana* population pairs estimated from 256bp blocks under the best-fit IM model (Table S19), including intergenic non-repetitive regions to minimize the direct effects of selection. *m_e_* estimates correspond to M (=4*N_e_m_e_*) individuals per generation (forwards in time). The chromosome 4 inversion is fixed in Crested Butte and absent in the other populations. The *m_e_* estimates were inferred under the best-fit IM model, except for the Vancouver-Storslett comparison where the *m_e_* estimates were based on the best-fit DIV model that assumes no post-divergence gene flow (*m_e_*=0).

| Population | Region | Ancestral $N_e$ | Pop. A $N_e$ | Pop. B $N_e$ | $T$ (years) | $m_e$ (M) |
| --- | --- | --- | --- | --- | --- | --- |
| Crested Butte (A) - Vancouver (B) | 4 INV | 646,563 | 179,131 | 1,137,026 | 1,347,905 | 2.06E-07 (0.15) |
| IM Vancouver --> Crested Butte | COL | 438,089 | 147,816 | 1,070,603 | 958,812 | 4.95E-07 (0.29) |
| Crested Butte (A) - Storslett (B) | 4 INV | 647,591 | 203,253 | 670,724 | 1,407,776 | 1.43E-07 (0.12) |
| IM Storslett --> Crested Butte | COL | 474,766 | 198,026 | 471,474 | 1,016,841 | 2.19E-07 (0.17) |
| Storslett (A) - Vancouver (B) | 4 INV | 808,211 | 561,850 | 1,159,870 | 727,619 | 0.0 (0.0) |
| IM Vancouver --> Storslett | COL | 527,819 | 413,484 | 1,228,026 | 651,134 | 0.0 (0.0) |

Finally, gIMble was used to identify long-term barrier loci between Crested Butte (inversion fixed) and Vancouver (inversion absent) in sliding windows to assess whether the inversion is enriched for barriers to gene flow (window-wise *m_e_* shown in Fig. S14). Local support for barriers to gene flow was defined by positive Δ_B0_ values with a false positive rate (FPR) of 5%, indicating that a strict divergence model (i.e. *m_e_*=0) fits better than the genome-wide background model of effective migration. As before, the barriers were explored strictly within the inversion breakpoints and by extending the inversion by 1.5Mb at both ends to account for the recombination-suppressing effects of inversions. Overall, 11.1% (1,050 out of 9,468) of the windows showed Δ_B0_>0, indicating potential barriers to gene flow (Fig. 6C). After accounting for an FPR of 5%, 44.4% (466 out of 1,050) of these windows passed the FPR criteria (Fig. 6C). By merging the remaining overlapping windows, a total of 10.7% (15.6Mb out of 145.5Mb) of the genome comprised barriers, distributed in 132 regions across the genome (Fig. 6C, Table S20). The barriers only partially overlapped with d_xy_ and F_st_ outliers (Fig. S15), a pattern also observed in *Heliconius melpomene* and *H. cydno* butterflies (Laetsch et al. 2023), highlighting the need for demographic modelling approaches to identify outlier barrier regions.

Notably, 62.9% (6.0 Mb out of 9.5 Mb) of the large inversion on chromosome 4 constituted barriers, compared to 7.0% (9.6 Mb out of 135.9 Mb) of the colinear regions across chromosomes. This difference was statistically significant, as determined by the number of barrier and non-barrier windows in inverted and colinear regions (Chi-Square: X² = 309.9, P < 0.0001; Fig. 6C, Table S20-S21). This pattern remained consistent when extending the inverted region beyond the breakpoints into the adjacent colinear regions (Chi-Square: X² = 337.4, P < 0.0001; Table S20-S21).

How closely associated are barrier and local adaptation loci? SNPs linked to PC1, PC2, CT_min_ and CCRT overlapped with the barrier regions more than expected by chance (Fig. 7). This suggests that loci underlying local adaptation may at least partly act as barrier loci, reducing gene flow between populations.

**Figure 7.**
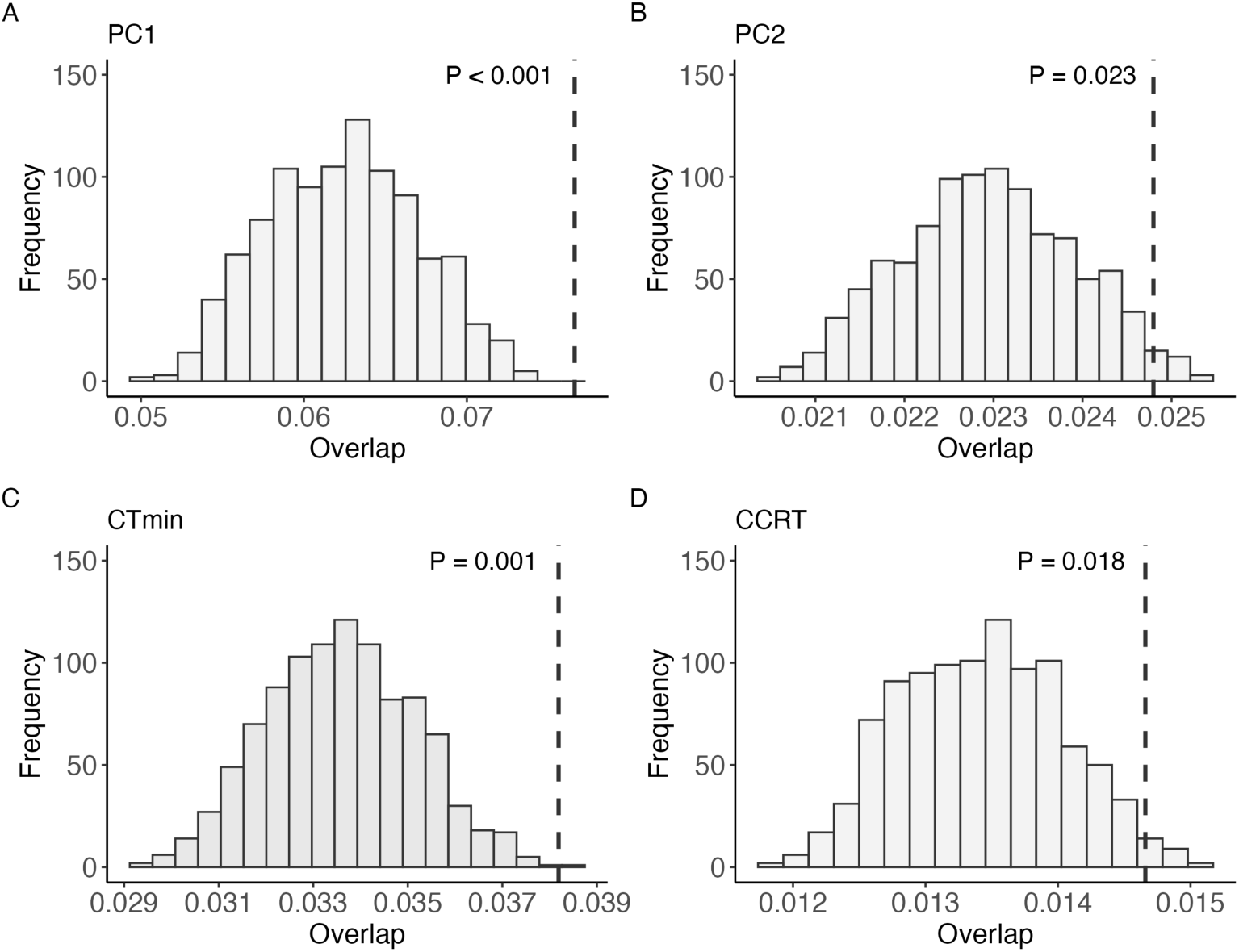
The observed (vertical dashed lines) and simulated (distributions) overlap between barrier windows and SNPs significantly associated with (A) PC1, (B) PC2, (C) CT_min_ and (D) CCRT. The significance of the circular resampling is indicated in each plot.

Our results suggest that *D. montana* populations from Crested Butte (NA), Vancouver (NA) and Storslett (Fennoscandia) diverged approximately 700,000-1,000,000 years ago. The estimate broadly aligns with a recent estimate of 1.75mya (Garlovsky et al. 2020). The origin of the chromosome 4 inversion predates these population split times based on the colinear background, indicating that the inversion originated in the ancestral *D. montana* population before it spread across North America, Asia, and Northern Europe. Similarly, species-specific inversions between closely related species *Drosophila persimilis* and *D. pseudoobscura* (Fuller et al. 2018) and *D. montana* and *D. flavomontana* (Poikela et al. 2024) originated in the ancestral population before species divergence. Interestingly, the polymorphic *D. montana* inversion, which has remained restricted to the North American Rocky Mountains populations, is enriched with both adaptive and barrier loci. This enrichment suggests that inversions can help maintain the association between multiple adaptive and barrier loci, effectively coupling them together. The significant overlap between the adaptive and barrier loci, in turn, suggests that genetic regions associated with climate adaptation may have reduced gene flow between populations, acting as a barrier, potentially due to ecologically-based divergent selection (Schluter 2001; Rundle and Nosil 2005). Other features of *D. montana* also act as potential barriers to gene flow and may contribute to the barrier loci we have identified. For instance, coastal and mountain *D. montana* populations exhibit partial sexual and postmating-prezygotic (PMPZ) isolation (Jennings, Snook, et al. 2014), likely driven by differences in courtship songs (Routtu et al. 2007), cuticular hydrocarbons (CHCs; Jennings, Etges, et al. 2014), and reproductive proteins (Garlovsky et al. 2020).

## Conclusions

Chromosomal inversions have been suggested to influence population adaptation and divergence by reducing gene flow and maintaining adaptive allelic combinations in linkage between populations or ecotypes with different arrangements. However, determining whether and how chromosomal inversions facilitate population adaptation and divergence is challenging due to sparse empirical evidence (Faria and Navarro 2010; Fuller et al. 2019) and the complexity of interacting processes that produce similar genomic signatures (Berdan et al. 2023). In this study, we characterize and investigate a large (∼9.5Mb) polymorphic inversion in *Drosophila montana*, a cold-adapted and widely distributed insect species.

The inversion’s origin predates the divergence of North American and Fennoscandian populations, suggesting it emerged in the ancestral *D. montana* population in the North American Rocky Mountains. Despite the species’ spread across the northern hemisphere, this inversion remains exclusive to the Rocky Mountains populations, being fixed in the southern high-altitude Crested Butte population and present at lower frequencies in the northern, lower-altitude Jackson and McBride populations. We independently mapped SNPs associated with climate adaptation and barriers to gene flow, and found that the inversion is a hotspot for both, with partial overlap between barrier and adaptive regions. This enrichment, along with its exclusive presence in the Rocky Mountain populations, further suggests that chromosomal inversions can play a role in environmental adaptation by restricting gene flow, facilitating adaptive divergence, and contributing to the coupling of different barriers to gene flow.

## Materials and methods

### *D. montana* sampling sites and overview of the data

*D. montana* samples were collected altogether from 13 geographically and climatically diverse sites in North America, Fennoscandia, and Asia in 2013−2020 (Fig. 1, Table S1). The newly collected females were brought to the fly laboratory in the University of Jyväskylä (Finland), and were kept in constant light, 19±1°C and ∼60% humidity. The females were allowed to lay eggs in malt vials for several days to produce the next generation, and to establish isofemale strains. The wild-caught females and their F_1_ daughters were then stored in 70% EtOH at −20°C. Furthermore, mass-bred populations were established from the F_4_-progenies of 16-20 isofemale strains (320-400 flies) per population.

The mass-bred populations or isofemale strains were used in the experiments measuring fly cold tolerance (Table S1). PacBio (Pacific BioSciences) long-read sequencing data was generated from pools of females from isofemale strains (Table S1) and used to characterize chromosomal inversions. Finally, Illumina short-read sequencing data was generated from wild-caught females or their F_1_-daughters collected and stored in 2013−2020 (Table S1). Illumina data were used to characterize chromosomal inversions and in genomic analyses. Details of the samples are described later in the text.

### Climatic conditions of *D. montana* sampling sites

A principal component analysis (PCA) was used to summarize differences in climatic conditions between the 13 *D. montana* sampling sites. For this, 19 bioclimatic variables of each sampling site were extracted from WorldClim database v2.1 (2.5min spatial resolution, dataset 1970-2000, http://www.worldclim.org, Fick and Hijmans 2017) using latitudinal and longitudinal coordinates of each site (Table S1) and “raster” R package v2.8-19 (Hijmans et al. 2015; Table S2). The PCA was performed for the 19 bioclimatic variables of each population using the “FactoMineR” R package (Lê et al. 2008). All R analyses were conducted in R (v4.2.0) and R studio (v2023.03.0).

### Assessment of *D. montana* cold tolerance

Variation in *D. montana* cold tolerance was assessed using two commonly used and ecologically relevant methods, chill coma temperature (CT_min_, also called critical thermal minimum) and chill coma recovery time (CCRT) (MacMillan and Sinclair 2011; Overgaard et al. 2011). CT_min_ describes insects’ ability to resist cold temperatures until falling into a reversible paralytic state, chill-coma, and CCRT their ability to recover from the chill-coma.

Fly cold tolerance was measured for nine *D. montana* populations: Seward, Terrace, Vancouver, Ashford, McBride, Crested Butte, Storslett, Oulanka and Korpilahti (Fig. 1, Table S1). Flies were collected from the mass-bred populations, apart from Oulanka for which the flies were collected from four isofemale strains. All flies had been maintained in the laboratory for 63−70 generations prior the experiments, except for Storslett and Oulanka which were maintained in the laboratory for 19 generations prior the experiments. To prevent the flies from entering reproductive diapause, they were maintained at constant light and 19±1℃ prior and throughout the experiments. Flies were provided fresh malt-medium once a week. Only female cold tolerance was measured, but variation between *D. montana* females and males has been studied in Wiberg et al. (2021). Females were collected using light CO_2_ anaesthesia within 72h after their emergence, placed in malt-vials, and used in cold tolerance experiments when sexually mature (18–23 days old, e.g. Jennings, Snook, et al. 2014).

A combined CT_min_-CCRT test was performed by placing females of each population individually in glass vials sealed with parafilm, which were then submerged into a 30% glycol bath. The CT_min_ test was initiated by decreasing the bath temperature from 19 °C to -6 °C at the rate of 0.5 °C/min, and the temperature at which a fly can no longer resist cold and loses neurophysiological activity and coordination and falls into a chill coma was scored (MacMillan and Sinclair 2011). After the CT_min_ test, the flies were left in -6 °C for 16 hours, after which they were returned into room temperature and the time required for each fly to recover from the chill coma and stand on its legs was measured (MacMillan and Sinclair 2011; Vesala et al. 2012). The tests were performed using Julabo F32-HL Refrigerated/Heating Circulator. The assays were done in 13 batches, each consisting of 19-42 females that were split between populations. For each population 40-44 females were tested, except for the Oulanka population where 16-26 females from each of the four isofemale strain were tested (altogether 413 females).

In statistical tests, CT_min_ data (Celsius degrees + 10°C because Gamma distribution does not allow negative values) and CCRT data (seconds) were used as response variables. Since the data was not normally distributed (Shapiro-Wilk test: CT_min_; W = 0.974, P < 0.001, CCRT; W = 0.851, P < 0.001), they were analysed with a generalized linear mixed model (GLMM) with Gamma distribution, using *glmmTMB* function from glmmTMB R package (Brooks et al. 2017). To define the relevant fixed effects (latitude, altitude and their interaction) and random effects (experimental batch and room temperature during the experiment), we performed model selection first on the fixed effects, and then for the random effects using the best-fit fixed effects. The best-fit model for CT_min_ and CCRT was chosen based on Akaike information criterion (AIC) value and Akaike weight using *aictab* function from AICcmodavg R package (Mazerolle and Mazerolle 2020). The best-fit model for CT_min_ included latitude, altitude, their interaction as fixed effects and experimental batch and room temperature as random effects, while the best-fit model for CCRT included latitude, altitude, their interaction as fixed effects, with no random effects.

### Genomic data

PacBio long-read sequencing data from 11 *D. montana* populations (60 females from one isofemale strain per population; Table S1, S7), Illumina short-read sequencing data of single females from 13 *D. montana* populations (3-11 females per population, totalling 101 samples; Table S1, S8) and Illumina pool-sequence data from six *D. montana* populations (each pool containing 49-50 individuals; Table S1, S9) were used in the genomic analyses. Six of the 11 PacBio data sets were generated in the current study, but other PacBio and all Illumina data sets were first published by Wiberg et al. (2021), Poikela et al. (2024), and Tahami et al. (2024). Details of DNA extraction methods and sequencing technologies are given in these publications, but the relevant information is summarized in Table S7-S9. For PacBio data, average raw read coverage for each sample varied between 14-77X (Table S7). For Illumina single female data, average coverage for each sample varied between 12-97X, except for monSE13F37 sample which was sequenced to 435X (Table S8). For the Illumina pool-sequence data the average coverage for each sample varied between 99-234X (Table S9).

*D. montana* chromosome-level reference genome (Seward) and the respective gene and repeat annotations were obtained from Poikela et al. (2024). To characterize inversions, contig-level assemblies of *D. montana* originating from Seward, Jackson, Crested Butte, Oulanka and Kamchatka were obtained from Poikela et al. (2024) and Tahami et al. (2024). Assembly statistics are given in Table S10.

### Characterization of inversions and their frequencies

The presence of polymorphic *D. montana* inversions was investigated i) by mapping-based approaches and ii) by aligning assembled genomes of different populations. First, the PacBio raw reads of each of the 11 samples (one per population, except no data from Vancouver and Korpilahti; Table S7) were mapped against the contig-level Seward genome with the highest contiguity (Table S10) using ngmlr v0.2.7 (Sedlazeck et al. 2018). The resulting BAM-files were sorted with SAMtools v1.16.1 (Li et al. 2009) and structural variant (SV) candidates were obtained from Sniffles v2.0.7 (Sedlazeck et al. 2018). Also, Illumina paired-end reads of 101 single wild-caught samples (3-11 samples per population; Table S8) and six pool-sequence samples (Table S9) were quality-checked using FastQC v0.11.8 (Andrews 2010), trimmed for adapter contamination and low-quality bases using fastp v0.20.0 (Chen et al. 2018) and mapped against the contig-level Seward genome using BWA mem v0.7.17 with read group information (Li and Durbin 2009). The resulting BAM-files were sorted with SAMtools and PCR duplicates marked with sambamba v0.7.0 (Tarasov et al. 2015). The BAM-files were given to Delly v0.8.7 for SV identification (Rausch et al. 2012). SV signals from Sniffles (PacBio) and Delly (Illumina) were combined using SURVIVOR (Jeffares et al. 2017) to identify inversion candidates. The putative breakpoints of inversions were confirmed visually with IGV v2.8.0 (Thorvaldsdottir et al. 2013) using both long- and short-read data (Example view in Fig. S2). Additionally, the Delly output and the visual inspection of the Illumina reads using IGV were used to determine whether individuals were homozygous or heterozygous inversion carriers. Second, alignments between *D. montana* contig-level assemblies (Table S10) were generated using nucmer of MUMmer package (Marçais et al. 2018) together with Dot plots (https://dot.sandbox.bio/). Finally, the breakpoints of reliable inversion candidates were mapped against the chromosome-level *D. montana* reference genome. Downstream analyses were focused on the large 9.5Mb inversion on chromosome 4 (Table S11), since the other characterized polymorphic inversion was tiny (∼1500bp; Table S12).

### Genetic diversity and divergence of *D. montana* populations

To define the genetic architecture of *D. montana*, the genetic variation of 101 Illumina single female samples was summarized using a principal component analysis (PCA). As explained in the previous paragraph, the Illumina raw reads were quality-checked, trimmed and mapped with read group information against the chromosome-level reference genome. The resulting BAM files were sorted, marked for PCR duplicates, and given to freebayes v1.3.6 for variant calling (Garrison and Marth 2012). The VCF file was filtered using preprocess function of gIMble with default settings (Laetsch et al. 2023). Variants with missing genotypes were excluded and the resulting VCF file was used to run the PCA i) for all chromosomes combined, ii) for each chromosome separately, iii) for the large chromosome 4 inversion (see Results), and iv) for the colinear regions of chromosome 4. All runs included coding and non-coding SNPs. The PCAs were run using PLINK’s (v1.9) --pca function (Chang et al. 2015), by converting the VCF-file to PLINK’s BED/BIM format and filtering SNPs and pruning them for linkage disequilibrium using PLINK’s --indep-pairwise function.

Genetic diversity (π) within *D. montana* populations, and genetic divergence (d_xy_) and differentiation (F_st_) between pairwise comparisons of *D. montana* populations were investigated using the filtered VCF-file and info and windows modules of gIMble. π, d_xy_ and F_st_ were first analysed for coding, intronic and intergenic regions across the chromosomes to investigate the genetic architecture of *D. montana* in general, and then for the inverted and colinear intergenic regions to estimate the effects of the inversion on genetic diversity and differentiation/divergence. Repetitive regions were excluded from all data partitions as they may bias the results. Window-wise heterozygosity (*H*), d_xy_ and F_st_ (including coding, intronic and intergenic regions) were also plotted across the genome.

### Signatures of local adaptation linked to climate conditions, cold tolerance and the chromosome 4 inversion

BayeScEnv is an F_st_ based genome-scan method that identifies loci linked to a given environmental variable or phenotype, while accounting for confounding effects of population structure (de Villemereuil and Gaggiotti 2015). We used the method to detect signatures of local adaptation linked to environmental conditions (PC1 and PC2 for each population) and cold phenotypes (scaled mean CT_min_ and CCRT for each population) of nine *D. montana* populations. Illumina paired-end reads of the nine populations, each consisting of 7-11 samples (Seward, Terrace, Vancouver, Ashford, McBride, Crested Butte, Storslett, Oulanka, and Korpilahti; Table S1, S8), were quality-checked and trimmed for adapter contamination and low-quality bases as described before. We then randomly resampled 3.6 million forward and reverse reads using seqtk (https://github.com/lh3/seqtk). We merged the resulting subsampled reads of each sample of a given population to obtain population-level forward and reverse reads. Reads of each population were mapped against the reference genome with read group information, and the alignments were sorted and marked for PCR duplicates, like before. Allele counts for each population at each genomic position were obtained with SAMtools mpileup using options to keep reads with a mapping quality >30 and sites with a base quality >30. The resulting mpileup file was used for variant calling using a heuristic variant calling software PoolSNP (Kapun et al. 2020). PoolSNP settings included a minimum count, coverage and frequency of 5, 10 and 0.05, respectively, a missing fraction of 0.05, and 95% coverage percentile for a given population and chromosome was considered as a maximum coverage. Variant calling detected a total of 3,547,377 biallelic SNPs.

The VCF-file was given to BayeScEnv v1.1, which was run twice for each dataset. Each run used the -pr_pref 0.5 setting and included five pilot runs of length of 1000, 2000 iterations and burn-in length of 2000. The MCMC chains were checked for convergence using coda R package v0.19-4.1 (Plummer et al. 2006), which was reached across the two chains for all analyses and parameters (potential scale reduction factors (PSRFs) of 1.00-1.03 in a Gelman-Rubin diagnostic test; Figures S7–S10). The union of significant SNPs, using q-values for the g parameter (FDR at 0.05, i.e. SNPs with q-values<0.05), were taken as the final candidate set. These SNPs describe the association between genetic differentiation and environmental or phenotypic differentiation.

The number of significant SNPs was compared between inverted and colinear regions using a Chi-square test to identify whether adaptive loci have enriched within the inversion. The effect of inversions in suppressing recombination may extend up to 2-3Mb beyond the inverted sequence (Kulathinal et al. 2009; Stevison et al. 2011). Thus, the effect of the inversion on adaptive loci was tested with a) the inversion ending strictly at the breakpoints and b) the inversion extending from each breakpoint into the colinear regions by a conservative estimate of 1.5Mb, as the impact of inversions on recombination has not been studied in *D. montana*.

Finally, we performed GO-term enrichment analysis on the significant SNPs using DAVID 2021 (Huang, Sherman, Tan, Kir, et al. 2007; Huang, Sherman, Tan, Collins, et al. 2007). We extracted all genes containing significant SNPs within the chromosome 4 inversion and blasted them against both *Drosophila virilis* and *Drosophila melanogaster* RefSeq proteins (BLASTp 2.15.0+; Camacho et al. 2009). While *D. montana* is more closely related to *D. virilis* (both belong to the *virilis* group; Yusuf et al. 2022) than to *D. melanogaster*, we wanted to take advantage of the superior annotation of *D. melanogaster*. Functional annotation clustering of the best *D. virilis* and *D. melanogaster* hits was carried out using DAVID.

### Evolutionary history of *D. montana* and the inversion on chromosome 4

We used gIMble (Laetsch et al. 2023) to resolve the evolutionary history of *D. montana* and to estimate when the large chromosome 4 inversion (see Results) originated and whether it has reduced gene flow between populations. gIMble is a composite likelihood approach that uses a joint distribution of mutation types in short sequence blocks, the blockwise site frequency spectrum (bSFS), across pairs of individual genomes.

First, a “global” mode of gIMble was used to investigate the likely evolutionary history of three genetically, geographically and climatically diverged *D. montana* populations, Vancouver (the western coast of North America), Crested Butte (the Rocky Mountains of North America) and Storslett (Fennoscandia), using the respective singly sequenced, wild-caught Illumina samples (Fig. 1, Table S1, Table S8). For each population pair, three demographic scenarios were fitted: a strict divergence without post-divergence gene flow (DIV, migration rate (*m_e_*) = 0), and isolation with migration with both gene flow directions between the populations (IM_monA→monB,_ IM_monB→monA_). This initial model selection was limited to intergenic, non-repetitive sequences of colinear chromosomes by excluding sequences with gene and repeat annotations to minimize the direct effects of selection. The large chromosome 4 inversion was excluded from the analysis due to its potentially different evolutionary history compared to colinear regions (Lohse et al. 2015; Fuller et al. 2018; Poikela et al. 2024), while colinear regions, ending at breakpoints, were combined across the genome as these regions are expected to share the same evolutionary history. Data was summarized by the bSFS for a block length of 256bp with kmax values of 2, and blocks were grouped in sliding windows (--blocks 12,500, --steps 2,500). The bSFS-based analyses assume no recombination within blocks and a constant mutation rate (μ) across them. Based on an estimate of the spontaneous mutation rate in *Drosophila melanogaster* (Keightley et al. 2014), μ = 2.8×10^-9^ per base and generation was used. The estimates of divergence time (*T*) were converted into absolute time using t = T×2N_e_×g, where N_e_ = θ/(4μ) and g is generation time. One generation per year was assumed for each population (Tyukmaeva et al. 2020). To evaluate the global support for an IM model versus a DIV model, we conducted 100 simulations using msprime (Baumdicker et al. 2022) through gIMble simulate, parameterised by the DIV history. Depending on the dataset, we simulated 8,343, 8,435 or 8,738 windows, matching the size of the real data, specifically 12,500 pair-blocks, for 10 diploid individuals per population, consistent with the empirical dataset. We assumed a constant recombination rate, based on one crossover per female meiosis per chromosome (recombination occurs only in females), resulting in map length of 25cM per chromosome. We fitted both a DIV model and an IM model to each simulation and compared the null distribution of lnCL to the observed empirical lnCL (see Fig. S12).

Second, a “global” mode of gIMble was used to estimate the time of divergence of the polymorphic inversion on chromosome 4 and the mean migration rate inside and outside of the inversion. For this, the best-fit model of each population pair was fitted separately for the inverted and colinear regions using the same settings as before. Here, a parametric bootstrap was performed to estimate the uncertainty of *T* and *M* parameters using gIMble simulate with 100 replicate datasets. These simulations used the same number of samples, block length, and number of blocks per window as the real data, and assumed the same mutation and recombination rates as before. Since the windows of the real data partially overlap and are in linkage, whereas the simulated windows are assumed to be statistically independent, we simulated datasets with 36 times fewer windows than the real data, resulting in approximately 500kb between windows, following Laetsch et al. (2023). Confidence intervals were estimated using the parametric bootstrap as +/- 2 SD across the 100 replicates.

Third, a “local” mode of gIMble was used to identify regions of reduced gene flow, referred to as barrier loci, across the genome between Crested Butte (inversion fixed) and Vancouver (no inversion). gIMble accounts for background selection through heterogeneity in effective population size (*N_e_*) and barriers to gene flow through heterogeneity in migration rate (*m_e_*). To quantify the heterogeneity in *m_e_* and *N_e_*, a grid of parameters was fitted to sliding windows of a fixed length of 12,500 pair-blocks with a step size of 2,500 pair-blocks along the genome. This resulted in 9,468 windows with a minimum span of 32 kb. Since only intergenic blocks (excluding repetitive sequences) were used, the window span was typically greater than 32 kb (median window span 68.6 kb; Fig. S16). Variation in *N_e_s* and *m_e_* across the windows were explored by searching for parameter combinations in a 12ξ12ξ12ξ16 grid (*N_e_* mon_A_, *N_e_* mon_B_, *N_e_* ancestral, *m_e_*; Fig. S17). Given that the inversion on chromosome 4 has diverged earlier, i.e. is older than the colinear regions, the value for *T* was allowed to be either 958,812 or 1,347,905 (COL vs. 4INV *T*; see Table 1). The mutation rate was set to μ = 2.9 × 10^-9^ per base and generation (Keightley et al. 2014). Windows were identified as barriers to gene flow when the support for a DIV model (i.e. *m_e_* = 0) was greater than the support for an IM model. The IM model was parameterized with the most likely inferred effective migration rate under a null model of *m_e_* variation, indicated by Δ_B0_ > 0. In other words, the local support for reduced gene flow was measured via Δ_B0_, where positive Δ_B0_ indicates strongly reduced gene flow. To estimate the uncertainty in these local estimates, we used a parametric bootstrap approach. Specifically, we simulated 100 datasets for each window under the best-fitting local background history using the msprime software (Baumdicker et al. 2022) via the gIMble simulate module, providing a null distribution of Δ_B0_ values. In these simulations, *m_e_* was fixed across windows at the global background level, while *N_e_* parameters were allowed to vary between windows based on the best composite log-likelihood *N_e_* parameters inferred for each window. Windows from the real data were labelled as barriers if their Δ_B0_ value was greater than the highest Δ_B0_ value observed in the simulated null distribution, ensuring a false positive rate (FPR) of ≤0.05. The same recombination rate was used in these simulations as before. Finally, windows with significantly reduced gene flow (Δ_B0_>0 and FPR≤0.05) that overlapped were merged to define barrier regions.

The number of the barrier and non-barrier windows (divided by five to obtain the number of non-overlapping windows) was compared between the inverted and colinear regions using a Chi-Square test to determine whether barriers are enriched within the inversion. Like before, the effect of the inversion on barrier loci was tested with a) the inversion ending strictly at the breakpoints and b) the inversion extending from each breakpoint into the colinear regions by a conservative estimate of 1.5Mb. Finally, to test whether SNPs significantly associated with PC1, PC2, CT_min_ or CCRT were enriched within barrier regions (proportion of SNPs in barrier vs non-barrier regions), we used a circular resampling approach (https://github.com/LohseLab/circular_bootstrap/tree/main) by Ebdon et al. (2025), where the observed estimate was compared to the distribution of means of 1000 resampled datasets.

## Acknowledgements

We want to thank Sara Lommi for her help with DNA extractions and cold tolerance experiments, and Oscar E. Gaggiotti for his advice on the BayeScEnv analyses. This work was supported by an Academy of Finland project 322980 to M.K., and grants from Jenny and Antti Wihuri Foundation to N.P. Fig. 1A was created with Biorender.com. Fig. 3 was created with MapMixture (https://tomjenkins.shinyapps.io/mapmixture/).

## Data Accessibility and Benefit-Sharing

### Data accessibility statement

The new PacBio raw sequence data are available at SRA under BioProject PRJNA1237710. The previously published raw sequence data and genome assemblies were obtained from BioProject accessions PRJNA939085, PRJNA828433, and PRJNA588720. Cold tolerance data and all scripts are available at https://github.com/noorlinnea/Dmontana_inversion_polymorphism.

### Benefit-Sharing Statement

Not applicable.

### Author contributions

MK and NP designed the study. NP and VH performed cold tolerance experiments. NP conducted statistical and genomic analyses, with input from VH and MGR. MK and MGR supervised, and MK funded the study. NP drafted the first version of the manuscript, and all authors edited and read the final draft.

### Ethics declaration

*D. montana* is not endangered, and the flies were collected along watersides on public lands outside National and State parks, where insect collecting does not require permits in the USA (The Wilderness Act of 1964, section 6302.15).

