## Supplementary material for "Examining the role of a chromosomal inversion in accumulating adaptive and barrier loci in a cold-adapted insect species"

### Supplementary Tables

Table S1. Summary of the *D. montana* data used in the study. Table shows sampling locations, coordinates and years, the data used in cold experiments, the isofemale strain ID sequenced with PacBio, the number of wild-caught individuals sequenced with Illumina, and the availability of poolseq data for each population. Details of the PacBio and Illumina samples are given in Tables S7-S9.

| Sampling location | Sampling year | Data used in cold experiments | Strain used in PacBio sequencing | Number of wild-caught females sequenced with Illumina | Illumina poolseq (49-50 single wild-caught females/males per pool) |
|---|---|---|---|---|---|
| **Western coast of North America** | | | | | |
| Seward, Alaska, USA<br>60°10'N; 149°27'W<br>Altitude 35 m | 2013 | mass-bred population | monSE13F37 | 11 | yes |
| Terrace, British Columbia, Canada<br>54°27'N; 128°34'W<br>Altitude 217 m | 2014 | mass-bred population | monTER14F11 | 11 | yes |
| Vancouver, British Columbia, Cana<br>49°15'N; 123°10'W<br>Altitude 4 m | 2014 | mass-bred population | - | 10 | no |
| Ashford, Washington, Usa<br>46°45'N; 121°57'W<br>Altitude 573 m | 2013 | mass-bred population | monASH13F9 | 10 | yes |
| Azalea, Oregon, USA<br>42°48'N; 123°13'W<br>Altitude 498 m | 2010 | - | monAZA2 | 3 | no |
| **Rocky Mountains of North America** | | | | | |
| McBride, British Columbia, Canad<br>53°18'N; 120°9'W<br>Altitude 720 m | 2014 | mass-bred population | monMB14F1 | 9 | no |
| Jackson, Wyoming, USA<br>43°26'N; 110°50'W<br>Altitude 1 857 m | 2013 | - | monJX13F48 | 3 | no |
| Crested Butte, Colorado, USA<br>39°49'N; 107°04'W<br>Altitude 2960 m | 2009 | mass-bred population | mon34CC5 | 10 | yes |
| **Fennoscandia** | | | | | |
| Storslett, Norway<br>69°36'N; 21°19'E<br>Altitude 2 m | 2020 | mass-bred population | monSTO20F2 | 10 | no |
| Raattama, Finland<br>68°01'N; 24°08'E<br>Altitude 343 m | 2020 | - | monRAA20F1 | 3 | no |
| Oulanka, Finland<br>66°22'N; 29°20'E<br>Altitude 341 m | 2013 | 5 isofemale strains: OU21F1, OU21F14, OU21F15, OU21F30 | monOU13F149 | 11 | yes |
| Korpilahti, Finland<br>62°00' N; 25°34'E<br>Altitude 85 m | 2013 | mass-bred population | - | 7 | yes |
| **Kamchatka** | | | | | |
| Kamchatka peninsula, Russia<br>56°12'N; 161°59'E<br>Altitude 21 m | 2013 | - | monKR1309 | 3 | no |

Table S2. Latitude, longitude, altitude and 19 bioclimatic variables and their description for each fly sampling site. The bioclimatic variables were extracted from WorldClim database v2.1 with 2.5 min spatial resolution using geographic coordinates (latitudes and longitudes) of each fly collecting site (Fick and Hijmans 2017). The darker (greener) the colour, the greater the value. NA = North America.

| Geographic region | Population | Latitude | Longitude | Altitude | bio1 | bio2 | bio3 | bio4 | bio5 | bio6 | bio7 | bio8 | bio9 | bio10 | bio11 | bio12 | bio13 | bio14 | bio15 | bio16 | bio17 | bio18 | bio19 |
|---|---|---|---|---|---|---|---|---|---|---|---|---|---|---|---|---|---|---|---|---|---|---|---|
| Western coast (NA) | Seward | 60.17 | -149.45 | 35 | 2.26 | 6.77 | 28.76 | 639.63 | 15.62 | -7.91 | 23.54 | 2.06 | 8.53 | 10.68 | -4.66 | 1266 | 197 | 54 | 44.92 | 505 | 178 | 213 | 342 |
| | Terrace | 54.45 | -128.57 | 217 | 6.41 | 7.72 | 27.57 | 730.96 | 21.38 | -6.63 | 28.01 | 1.75 | 13.43 | 15.32 | -2.71 | 1379 | 210 | 50 | 51.67 | 578 | 157 | 170 | 487 |
| | Vancouver | 49.25 | -123.17 | 4 | 10.28 | 6.76 | 31.70 | 533.84 | 22.28 | 0.95 | 21.34 | 4.59 | 16.87 | 17.01 | 4.04 | 1440 | 225 | 45 | 51.90 | 620 | 154 | 157 | 536 |
| | Ashford | 46.75 | -121.95 | 573 | 7.55 | 10.18 | 41.90 | 514.42 | 22.52 | -1.77 | 24.29 | 2.13 | 14.00 | 14.22 | 1.86 | 2025 | 309 | 43 | 57.67 | 917 | 178 | 189 | 832 |
| | Azalea | 42.80 | -123.22 | 498 | 11.05 | 13.74 | 48.03 | 555.79 | 28.90 | 0.30 | 28.60 | 5.03 | 18.11 | 18.20 | 4.80 | 1145 | 196 | 13 | 70.09 | 559 | 62 | 71 | 507 |
| Rocky Mountains (NA) | McBride | 53.30 | -120.16 | 720 | 4.07 | 11.31 | 32.24 | 850.10 | 22.13 | -12.94 | 35.07 | 4.04 | 0.10 | 14.23 | -6.66 | 706 | 75 | 39 | 21.11 | 210 | 127 | 201 | 163 |
| | Jackson | 43.43 | -110.83 | 1857 | 3.65 | 16.12 | 38.11 | 943.61 | 27.08 | -15.21 | 42.29 | 8.40 | -1.65 | 15.16 | -8.27 | 439 | 50 | 27 | 20.40 | 128 | 90 | 108 | 115 |
| | Crested Butte | 39.82 | -107.07 | 2960 | 3.58 | 17.14 | 42.68 | 848.91 | 25.25 | -14.92 | 40.16 | 13.79 | -6.56 | 14.09 | -6.56 | 430 | 45 | 29 | 13.34 | 127 | 97 | 122 | 97 |
| Fennoscandia | Storslett | 69.60 | 21.32 | 95 | 1.42 | 5.85 | 22.07 | 757.24 | 15.73 | -10.80 | 26.53 | 10.34 | -4.12 | 11.32 | -7.17 | 482 | 62 | 26 | 28.02 | 163 | 83 | 153 | 110 |
| | Raattama | 68.02 | 24.13 | 343 | -1.74 | 9.03 | 24.14 | 995.77 | 17.76 | -19.64 | 37.41 | 11.12 | -8.01 | 11.12 | -13.60 | 523 | 85 | 26 | 44.73 | 221 | 79 | 221 | 87 |
| | Oulanka | 66.37 | 29.33 | 341 | -0.81 | 9.74 | 24.54 | 1037.88 | 19.78 | -19.91 | 39.69 | 12.43 | -7.76 | 12.43 | -13.48 | 571 | 76 | 29 | 31.99 | 206 | 92 | 206 | 101 |
| | Korpilahti | 62.00 | 25.57 | 85 | 3.51 | 7.58 | 23.02 | 899.21 | 21.16 | -11.75 | 32.91 | 13.17 | -3.19 | 14.98 | -7.34 | 614 | 86 | 29 | 34.21 | 225 | 100 | 220 | 116 |
| Kamchatka | Kamchatka | 56.20 | 161.98 | 21 | -0.46 | 7.27 | 22.74 | 928.28 | 15.79 | -16.18 | 31.97 | -9.46 | 2.60 | 10.79 | -11.69 | 691 | 77 | 35 | 26.39 | 218 | 110 | 171 | 211 |

| | |
|---|---|
| bio1 | Annual mean temperature |
| bio2 | Mean diurnal range (mean of monthly (max - min temperature)) |
| bio3 | Isothermality (bio2/bio7*100) |
| bio4 | Temperature seasonality (standard deviation * 100) |
| bio5 | Max temperature of the warmest month |
| bio6 | Min temperature of the coldest month |
| bio7 | Annual temperature range (bio5-bio6) |
| bio8 | Mean temperature of the wettest quarter |
| bio9 | Mean temperature of the driest quarter |
| bio10 | Mean temperature of the warmest quarter |
| bio11 | Mean temperature of the coldest quarter |
| bio12 | Annual precipitation |
| bio13 | Precipitation of the wettest month |
| bio14 | Precipitation of the driest month |
| bio15 | Precipitation seasonality (coefficient of variation) |
| bio16 | Precipitation of the wettest quarter |
| bio17 | Precipitation of the driest quarter |
| bio18 | Precipitation of the warmest quarter |
| bio19 | Precipitation of the coldest quarter |

Table S3. Principal components and their Eigenvalues, variance, and cumulative variance for the bioclimatic variables.

| PC | Eigenvalue | Variance (%) | Cumulative variance (%) |
|---|---|---|---|
| PC1 | 10.6 | 55.9 | 55.9 |
| PC2 | 4.8 | 25.4 | 81.2 |
| PC3 | 1.2 | 6.4 | 87.6 |
| PC4 | 0.9 | 4.9 | 92.5 |
| PC5 | 0.7 | 3.7 | 96.2 |
| PC6 | 0.5 | 2.5 | 98.7 |
| PC7 | 0.1 | 0.7 | 99.4 |
| PC8 | 0.1 | 0.5 | 99.9 |
| PC9 | 0.0 | 0.0 | 100.0 |
| PC10 | 0.0 | 0.0 | 100.0 |
| PC11 | 0.0 | 0.0 | 100.0 |
| PC12 | 0.0 | 0.0 | 100.0 |

Table S4. Contributions (loadings) of the bioclimatic variables (Table S2) on each principal component (PC). The darker (greener) the shading, the greater the contribution of a given variable on each PC.

| Variable | PC1 | PC2 | PC3 | PC4 | PC5 |
|---|---|---|---|---|---|
| **bio1** | 7.0 | 3.9 | 0.1 | 0.1 | 7.2 |
| **bio2** | 0.2 | 15.3 | 16.8 | 0.0 | 3.1 |
| **bio3** | 2.3 | 11.6 | 7.9 | 0.1 | 3.5 |
| **bio4** | 8.4 | 0.0 | 0.7 | 0.1 | 2.8 |
| **bio5** | 0.8 | 17.1 | 3.1 | 1.6 | 0.0 |
| **bio6** | 8.6 | 0.3 | 2.5 | 0.0 | 5.1 |
| **bio7** | 5.9 | 4.2 | 7.4 | 0.9 | 5.7 |
| **bio8** | 1.6 | 3.3 | 2.9 | 59.2 | 3.7 |
| **bio9** | 8.7 | 0.0 | 0.2 | 4.4 | 0.0 |
| **bio10** | 3.0 | 9.6 | 0.3 | 2.7 | 6.4 |
| **bio11** | 8.2 | 1.6 | 0.6 | 0.0 | 5.1 |
| **bio12** | 8.3 | 1.3 | 1.5 | 1.1 | 3.9 |
| **bio13** | 8.4 | 0.9 | 0.1 | 2.2 | 5.7 |
| **bio14** | 1.6 | 9.8 | 21.0 | 0.4 | 9.3 |
| **bio15** | 5.9 | 0.0 | 12.8 | 2.2 | 20.5 |
| **bio16** | 8.4 | 0.6 | 0.2 | 1.9 | 7.4 |
| **bio17** | 3.5 | 7.0 | 20.8 | 1.3 | 3.4 |
| **bio18** | 0.5 | 13.3 | 0.7 | 21.8 | 1.1 |
| **bio19** | 8.6 | 0.1 | 0.6 | 0.0 | 6.1 |

Table S5. The best-fit model for $CT_{min}$ and CCRT was defined based on Akaike Information Criterion (AIC) and it was performed in two steps. First, the best-fit fixed effects were defined (highlighted with light grey). Then, the best-fit random effects were defined using the best-fit fixed effects, i.e. this is the final best-fit model used for further analyses (highlighted with dark grey). The model-fit was established using the highest Akaike weight (AICcWt), i.e. probability of being the best model. T refers to temperature during the experiment.

| Experiment | Model | | df | AICc | ΔAICc | AICcWt |
|---|---|---|---|---|---|---|
| $CT_{min}$ | Model selection for fixed effects | latitude + altitude + latitude:altitude | 5 | 776.51 | 0.00 | 0.99 |
| | | latitude + altitude | 4 | 786.56 | 10.05 | 0.01 |
| | | latitude | 3 | 794.84 | 18.33 | 0.00 |
| | Model selection for randon effects using the best-fit fixed effects, i.e. the final best-fit model | latitude + altitude + latitude:altitude + experimental batch + T | 7 | 737.54 | 0.00 | 0.75 |
| | | latitude + altitude + latitude:altitude + experimental batch | 6 | 770.24 | 32.70 | 0.00 |
| | | latitude + altitude + latitude:altitude + T | 6 | 739.77 | 2.23 | 0.25 |
| | | latitude + altitude + latitude:altitude (no random effects) | 5 | 776.51 | 38.97 | 0.00 |
| CCRT | Model selection for fixed effects | latitude + altitude + latitude:altitude | 5 | 4906.63 | 0 | 0.91 |
| | | latitude + altitude | 4 | 4913.96 | 7.32 | 0.02 |
| | | latitude | 3 | 4911.93 | 5.3 | 0.06 |
| | Model selection for randon effects using the best-fit fixed effects, i.e. the final best-fit model | latitude + altitude + latitude:altitude + experimental batch + T | 7 | 4910.65 | 4.02 | 0.07 |
| | | latitude + altitude + latitude:altitude + experimental batch | 6 | 4908.57 | 1.94 | 0.2 |
| | | latitude + altitude + latitude:altitude + T | 6 | 4908.7 | 2.07 | 0.19 |
| | | latitude + altitude + latitude:altitude (no random effects) | 5 | 4906.63 | 0 | 0.53 |

Table S6. Pearson correlation test between $CT_{min}$ and CCRT across all populations and separately for each population. Significant correlations are highlighted.

| Geographical region | Population | Pearson correlation coefficient | t statistic | df | P value |
|---|---|---|---|---|---|
| All regions together | All populations together | 0.217 | 4.04 | 330 | **6.65E-05** |
| Western coast of NA | Seward | 0.387 | 2.30 | 30 | **0.029** |
| | Terrace | 0.421 | 2.59 | 31 | **0.015** |
| | Vancouver | 0.190 | 1.10 | 32 | 0.281 |
| | Ashford | 0.034 | 0.19 | 33 | 0.848 |
| Rocky Mountains of NA | McBride | -0.112 | -0.62 | 30 | 0.541 |
| | Crested Butte | 0.301 | 1.73 | 30 | 0.094 |
| Fennoscandia | Storslett | 0.111 | 0.62 | 31 | 0.540 |
| | Oulanka | -0.102 | -0.82 | 64 | 0.417 |
| | Korpilahti | 0.233 | 1.37 | 33 | 0.179 |

Table S7. Details of *D. montana* PacBio raw read datasets, which were used to characterize inversions. N50 raw reads = half of the raw reads are larger than or equal to the N50 raw read value.

| PacBio reads | | | | # SMRT | | Estimated | Max | Average | | Number of reads | | | Sequencing | |
|---|---|---|---|---|---|---|---|---|---|---|---|---|---|---|
| **Region** | **Population** | **Strain ID** | **# reads** | **cells** | **Bases (Gb)** | **Mean coverage** | **read length (bp)** | **read length (bp)** | **N50 raw read (bp)** | **>= 80 000 bp** | **Technique** | **Sequencing facility** | **year** | **First published** |
| Western coast | Seward | monSE13F37 | 1,670,083 | 2 | 9.9 | 54.8 | 168,303 | 5,906 | 11,366 | 132 | PacBio Sequel I | Norwegian sequencing center | 2018 | Poikela et al. (2024) |
| of North America | Terrace | monTER14F11 | 328,728 | 1 | 3.2 | 17.9 | 154,030 | 9,782 | 14,605 | 28 | PacBio Sequel II | Novogene | 2021 | current study |
| | Ashford | monASH13F9 | 426,503 | 1 | 4.1 | 23.0 | 131,309 | 9,706 | 13,952 | 51 | PacBio Sequel II | Novogene | 2021 | current study |
| | Azalea | monAZA1 | 233,260 | 1 | 2.5 | 13.8 | 107,310 | 10,623 | 15,435 | 29 | PacBio Sequel II | Novogene | 2021 | current study |
| Rocky Mountains | McBride | monMB14F1 | 347,232 | 1 | 3.2 | 17.6 | 113,197 | 9,131 | 13,176 | 36 | PacBio Sequel II | Novogene | 2021 | current study |
| of North America | Jackson | monJX13F48 | 2,161,617 | 1 | 13.9 | 77.1 | 111,203 | 6,422 | 8,045 | 19 | PacBio Sequel I | BGI | 2019 | Poikela et al. (2024) |
| | Crested Butte | mon34CC5 | 1,281,079 | 3 | 7.2 | 39.9 | 145,085 | 5,608 | 10,260 | 248 | PacBio Sequel II | Novogene | 2021 | Tahami et al. (2024) |
| Fennoscandia | Storslett | monSTO20F2 | 917,273 | 2 | 2.8 | 15.5 | 146,182 | 3,046 | 3,789 | 177 | PacBio Sequel II | Novogene | 2021 | current study |
| | Raattama | monRAA20F1 | 900,722 | 2 | 3.0 | 16.5 | 133,674 | 3,304 | 4,436 | 190 | PacBio Sequel II | Novogene | 2021 | current study |
| | Oulanka | monOU13F149 | 1,431,344 | 1 | 8.3 | 46.4 | 118,876 | 5,830 | 8,836 | 33 | PacBio Sequel I | BGI | 2019 | Tahami et al. (2024) |
| Asia | Kamchatka | monKR1309 | 879,152 | 1 | 5.5 | 30.6 | 132,961 | 6,269 | 9,233 | 53 | PacBio Sequel I | BGI | 2019 | Tahami et al. (2024) |

Table S8. Details of Illumina paired-end raw reads of 101 singly sequenced wild-caught *D. montana* females or their $F_1$ daughters (continues the next page), used in genomic analyses. The data were first published by Poikela et al. (2024) and Tahami et al. (2024).

| Illumina 150bp paired-end reads | | | | | | Mean | | | | Sequencing | Sequenced | |
|---|---|---|---|---|---|---|---|---|---|---|---|---|
| Region | Population | Strain ID | # reads (M) | Bases (Gb) | # lanes | coverage | Library | Technique | Sequencing facility | year | generation | First published |
| Western coast of North America | Seward | monSE13F11 | 80.3 | 12.0 | 1 | 55.3 | Truseq | HiSeq X-Ten | Novogene | 2021 | $F_1$ | Tahami et al. (2024) |
| | | monSE13F12 | 110.4 | 16.6 | 1 | 60.3 | Truseq | HiSeq X-Ten | Novogene | 2021 | $F_1$ | Tahami et al. (2024) |
| | | monSE13F16 | 77.3 | 11.6 | 1 | 51.1 | Truseq | HiSeq X-Ten | Novogene | 2021 | $F_1$ | Tahami et al. (2024) |
| | | monSE13F20 | 80.2 | 12.0 | 1 | 55.5 | Truseq | HiSeq X-Ten | Novogene | 2021 | $F_1$ | Tahami et al. (2024) |
| | | monSE13F21 | 79.3 | 11.9 | 1 | 55.2 | Truseq | HiSeq X-Ten | Novogene | 2021 | $F_1$ | Tahami et al. (2024) |
| | | monSE13F22 | 68.2 | 10.2 | 1 | 39.7 | Truseq | HiSeq X-Ten | Novogene | 2021 | $F_1$ | Tahami et al. (2024) |
| | | monSE13F3 | 71.6 | 10.7 | 1 | 51.1 | Truseq | HiSeq X-Ten | Novogene | 2021 | $F_1$ | Tahami et al. (2024) |
| | | monSE13F37 | 630.7 | 95.2 | 1 | 434.5 | Truseq | HiSeq4000 | Norwegian sequencing center | 2018 | $F_1$ | Poikela et al. (2024) |
| | | monSE13F5 | 78.6 | 11.8 | 1 | 53.8 | Truseq | HiSeq X-Ten | Novogene | 2021 | $F_1$ | Tahami et al. (2024) |
| | | monSE13F7 | 79.5 | 11.9 | 1 | 50.9 | Truseq | HiSeq X-Ten | Novogene | 2021 | $F_1$ | Tahami et al. (2024) |
| | | monSE13F9 | 87.3 | 13.1 | 1 | 49.6 | Truseq | HiSeq X-Ten | Novogene | 2021 | $F_1$ | Tahami et al. (2024) |
| | Terrace | monTER14F10 | 76.2 | 11.4 | 1 | 41.6 | Truseq | HiSeq X-Ten | Novogene | 2021 | wild-caught | Tahami et al. (2024) |
| | | monTER14F11 | 96.8 | 14.5 | 1 | 67.3 | Truseq | HiSeq X-Ten | BGI | 2019 | wild-caught | Poikela et al. (2024) |
| | | monTER14F12 | 93.4 | 14.0 | 1 | 27.1 | Truseq | HiSeq X-Ten | Novogene | 2021 | wild-caught | Tahami et al. (2024) |
| | | monTER14F13 | 68.6 | 10.3 | 1 | 44.1 | Truseq | HiSeq X-Ten | Novogene | 2021 | wild-caught | Tahami et al. (2024) |
| | | monTER14F16 | 88.8 | 13.3 | 1 | 55.5 | Truseq | HiSeq X-Ten | Novogene | 2021 | wild-caught | Tahami et al. (2024) |
| | | monTER14F17 | 135.6 | 20.3 | 2 | 68.8 | Truseq | HiSeq X-Ten | Novogene | 2021 | wild-caught | Tahami et al. (2024) |
| | | monTER14F19 | 65.3 | 9.8 | 1 | 34.5 | Truseq | HiSeq X-Ten | Novogene | 2021 | wild-caught | Tahami et al. (2024) |
| | | monTER14F2 | 76.2 | 11.4 | 1 | 42.3 | Truseq | HiSeq X-Ten | Novogene | 2021 | wild-caught | Tahami et al. (2024) |
| | | monTER14F21 | 94.0 | 14.1 | 1 | 63.6 | Truseq | HiSeq X-Ten | Novogene | 2021 | wild-caught | Tahami et al. (2024) |
| | | monTER14F24 | 129.9 | 19.5 | 2 | 72.3 | Truseq | HiSeq X-Ten | Novogene | 2021 | wild-caught | Tahami et al. (2024) |
| | | monTER14F26 | 65.8 | 9.9 | 1 | 44.5 | Truseq | HiSeq X-Ten | Novogene | 2021 | wild-caught | Tahami et al. (2024) |
| | Vancouver | monVAN14F1 | 121.5 | 18.2 | 2 | 70.6 | Nextera | HiSeq4000 | Edinburgh Genomics | 2017 | wild-caught | Poikela et al. (2024) |
| | | monVAN14F10 | 80.5 | 12.1 | 1 | 48.4 | Truseq | HiSeq X-Ten | Novogene | 2021 | wild-caught | Tahami et al. (2024) |
| | | monVAN14F11 | 65.6 | 9.8 | 1 | 46.4 | Truseq | HiSeq X-Ten | Novogene | 2021 | wild-caught | Tahami et al. (2024) |
| | | monVAN14F17 | 68.1 | 10.2 | 1 | 43.2 | Truseq | HiSeq X-Ten | Novogene | 2021 | wild-caught | Tahami et al. (2024) |
| | | monVAN14F24 | 66.6 | 10.0 | 1 | 33.3 | Truseq | HiSeq X-Ten | Novogene | 2021 | wild-caught | Tahami et al. (2024) |
| | | monVAN14F4 | 86.9 | 13.0 | 1 | 41.7 | Truseq | HiSeq X-Ten | Novogene | 2021 | wild-caught | Tahami et al. (2024) |
| | | monVAN14F5 | 79.5 | 11.9 | 1 | 44.4 | Truseq | HiSeq X-Ten | Novogene | 2021 | wild-caught | Tahami et al. (2024) |
| | | monVAN14F6 | 73.8 | 11.1 | 1 | 33.4 | Truseq | HiSeq X-Ten | Novogene | 2021 | wild-caught | Tahami et al. (2024) |
| | | monVAN14F7 | 84.0 | 12.6 | 1 | 56.3 | Truseq | HiSeq X-Ten | Novogene | 2021 | wild-caught | Tahami et al. (2024) |
| | | monVAN14F8 | 75.9 | 11.4 | 1 | 40.6 | Truseq | HiSeq X-Ten | Novogene | 2021 | wild-caught | Tahami et al. (2024) |
| | Ashford | monASH13F11 | 77.2 | 11.6 | 1 | 38.1 | Truseq | HiSeq X-Ten | Novogene | 2021 | $F_1$ | Tahami et al. (2024) |
| | | monASH13F12 | 75.8 | 11.4 | 1 | 48.9 | Truseq | HiSeq X-Ten | Novogene | 2021 | $F_1$ | Tahami et al. (2024) |
| | | monASH13F13 | 167.7 | 25.2 | 2 | 96.3 | Nextera | HiSeq4000 | Edinburgh Genomics | 2017 | $F_1$ | Poikela et al. (2024) |
| | | monASH13F17 | 97.6 | 14.6 | 1 | 71.3 | Truseq | HiSeq X-Ten | Novogene | 2021 | $F_1$ | Tahami et al. (2024) |
| | | monASH13F18 | 91.9 | 13.8 | 1 | 59.3 | Truseq | HiSeq X-Ten | Novogene | 2021 | $F_1$ | Tahami et al. (2024) |
| | | monASH13F20 | 92.7 | 13.9 | 1 | 66.7 | Truseq | HiSeq X-Ten | Novogene | 2021 | $F_1$ | Tahami et al. (2024) |
| | | monASH13F4 | 75.0 | 11.2 | 1 | 52.6 | Truseq | HiSeq X-Ten | Novogene | 2021 | $F_1$ | Tahami et al. (2024) |
| | | monASH13F5 | 77.7 | 11.7 | 1 | 55.8 | Truseq | HiSeq X-Ten | Novogene | 2021 | $F_1$ | Tahami et al. (2024) |
| | | monASH13F7 | 76.8 | 11.5 | 1 | 46.4 | Truseq | HiSeq X-Ten | Novogene | 2021 | $F_1$ | Tahami et al. (2024) |
| | | monASH13F9 | 82.8 | 12.4 | 1 | 50.3 | Truseq | HiSeq X-Ten | Novogene | 2021 | $F_1$ | Tahami et al. (2024) |
| | Azalea | monAZA1 | 72.8 | 10.9 | 1 | 52.1 | Truseq | HiSeq X-Ten | Novogene | 2021 | $F_1$ | Tahami et al. (2024) |
| | | monAZA2 | 94.6 | 14.2 | 1 | 70.7 | Truseq | HiSeq X-Ten | BGI | 2019 | $F_1$ | Poikela et al. (2024) |
| | | monAZA3 | 90.0 | 13.5 | 1 | 64.1 | Truseq | HiSeq X-Ten | Novogene | 2021 | $F_1$ | Tahami et al. (2024) |
| Rocky Mountains of North America | McBride | monMB14F1 | 129.5 | 19.4 | 1 | 53.8 | Truseq | HiSeq X-Ten | BGI | 2019 | wild-caught | Poikela et al. (2024) |
| | | monMB14F15 | 74.1 | 11.1 | 1 | 52.4 | Truseq | HiSeq X-Ten | Novogene | 2021 | wild-caught | Tahami et al. (2024) |
| | | monMB14F16 | 65.5 | 9.8 | 1 | 43.5 | Truseq | HiSeq X-Ten | Novogene | 2021 | wild-caught | Tahami et al. (2024) |
| | | monMB14F19 | 78.9 | 11.8 | 1 | 56.1 | Truseq | HiSeq X-Ten | Novogene | 2021 | wild-caught | Tahami et al. (2024) |
| | | monMB14F22 | 85.3 | 12.8 | 1 | 61.8 | Truseq | HiSeq X-Ten | Novogene | 2021 | wild-caught | Tahami et al. (2024) |
| | | monMB14F5 | 67.7 | 10.2 | 1 | 43.5 | Truseq | HiSeq X-Ten | Novogene | 2021 | wild-caught | Tahami et al. (2024) |
| | | monMB14F6 | 71.4 | 10.7 | 1 | 50.3 | Truseq | HiSeq X-Ten | Novogene | 2021 | wild-caught | Tahami et al. (2024) |
| | | monMB14F7 | 85.2 | 12.8 | 1 | 60.6 | Truseq | HiSeq X-Ten | Novogene | 2021 | wild-caught | Tahami et al. (2024) |
| | | monMB14F9 | 74.2 | 11.1 | 1 | 40.0 | Truseq | HiSeq X-Ten | Novogene | 2021 | wild-caught | Tahami et al. (2024) |
| | Jackson | monJX13F3 | 130.0 | 19.5 | 2 | 83.4 | Nextera | HiSeq4000 | Edinburgh Genomics | 2017 | $F_1$ | Poikela et al. (2024) |
| | | monJX13F41 | 79.6 | 11.9 | 1 | 51.3 | Truseq | HiSeq X-Ten | Novogene | 2021 | $F_1$ | Tahami et al. (2024) |
| | | monJX13F48 | 68.0 | 10.2 | 1 | 23.8 | Truseq | HiSeq X-Ten | Novogene | 2021 | $F_1$ | Tahami et al. (2024) |
| | Crested Butte | mon17CC5 | 112.2 | 16.8 | 2 | 82.2 | Truseq | HiSeq X-Ten | Novogene | 2021 | $F_1$ | Tahami et al. (2024) |
| | | mon26CC5 | 79.2 | 11.9 | 1 | 57.1 | Truseq | HiSeq X-Ten | Novogene | 2021 | $F_1$ | Tahami et al. (2024) |
| | | mon29CC5 | 69.6 | 10.4 | 1 | 49.2 | Truseq | HiSeq X-Ten | Novogene | 2021 | $F_1$ | Tahami et al. (2024) |
| | | mon37CC5 | 69.5 | 10.4 | 1 | 48.7 | Truseq | HiSeq X-Ten | Novogene | 2021 | $F_1$ | Tahami et al. (2024) |
| | | mon48CC5 | 94.5 | 14.2 | 1 | 65.3 | Truseq | HiSeq X-Ten | Novogene | 2021 | $F_1$ | Tahami et al. (2024) |
| | | mon4CC5 | 133.5 | 20.0 | 2 | 95.0 | Truseq | HiSeq X-Ten | Novogene | 2021 | $F_1$ | Tahami et al. (2024) |
| | | mon68CC5 | 69.9 | 10.5 | 1 | 45.6 | Truseq | HiSeq X-Ten | Novogene | 2021 | $F_1$ | Tahami et al. (2024) |
| | | mon77CC5 | 76.2 | 11.4 | 1 | 48.7 | Truseq | HiSeq X-Ten | Novogene | 2021 | $F_1$ | Tahami et al. (2024) |
| | | mon81CC5 | 87.2 | 13.1 | 1 | 18.9 | Truseq | HiSeq X-Ten | Novogene | 2021 | $F_1$ | Tahami et al. (2024) |
| | | mon8CC5 | 133.3 | 20.0 | 2 | 96.5 | Truseq | HiSeq X-Ten | Novogene | 2021 | $F_1$ | Tahami et al. (2024) |

| | | | | | | | | | | | | |
|---|---|---|---|---|---|---|---|---|---|---|---|---|
| Fennoscandia | Storslett | monSTO20F10 | 71.9 | 10.8 | 1 | 51.5 | Truseq | HiSeq X-Ten | Novogene | 2021 | F1 | Tahami et al. (2024) |
| | | monSTO20F11 | 109.5 | 16.4 | 1 | 72.5 | Truseq | HiSeq X-Ten | Novogene | 2021 | F1 | Tahami et al. (2024) |
| | | monSTO20F13 | 69.1 | 10.4 | 1 | 50.4 | Truseq | HiSeq X-Ten | Novogene | 2021 | F1 | Tahami et al. (2024) |
| | | monSTO20F14 | 74.9 | 11.2 | 1 | 54.1 | Truseq | HiSeq X-Ten | Novogene | 2021 | F1 | Tahami et al. (2024) |
| | | monSTO20F15 | 66.0 | 9.9 | 1 | 47.6 | Truseq | HiSeq X-Ten | Novogene | 2021 | F1 | Tahami et al. (2024) |
| | | monSTO20F2 | 69.1 | 10.4 | 1 | 11.7 | Truseq | HiSeq X-Ten | Novogene | 2021 | F1 | Tahami et al. (2024) |
| | | monSTO20F3 | 81.3 | 12.2 | 1 | 25.5 | Truseq | HiSeq X-Ten | Novogene | 2021 | F1 | Tahami et al. (2024) |
| | | monSTO20F7 | 92.4 | 13.9 | 1 | 26.5 | Truseq | HiSeq X-Ten | Novogene | 2021 | F1 | Tahami et al. (2024) |
| | | monSTO20F8 | 86.2 | 12.9 | 1 | 33.4 | Truseq | HiSeq X-Ten | Novogene | 2021 | F1 | Tahami et al. (2024) |
| | | monSTO20F9 | 114.6 | 17.2 | 1 | 37.7 | Truseq | HiSeq X-Ten | Novogene | 2021 | F1 | Tahami et al. (2024) |
| | Raattama | monRAA20F1 | 79.0 | 11.8 | 1 | 51.8 | Truseq | HiSeq X-Ten | Novogene | 2021 | F1 | Tahami et al. (2024) |
| | | monRAA20F4 | 69.7 | 10.5 | 1 | 50.3 | Truseq | HiSeq X-Ten | Novogene | 2021 | F1 | Tahami et al. (2024) |
| | | monRAA20F5 | 65.4 | 9.8 | 1 | 46.4 | Truseq | HiSeq X-Ten | Novogene | 2021 | F1 | Tahami et al. (2024) |
| | Oulanka | monOU13F140 | 70.7 | 10.6 | 1 | 49.4 | Truseq | HiSeq X-Ten | Novogene | 2021 | F1 | Tahami et al. (2024) |
| | | monOU13F149 | 98.8 | 14.8 | 1 | 72.5 | Truseq | HiSeq X-Ten | BGI | 2019 | F1 | Tahami et al. (2024) |
| | | monOU13F163 | 79.4 | 11.9 | 1 | 33.3 | Truseq | HiSeq X-Ten | Novogene | 2021 | F1 | Tahami et al. (2024) |
| | | monOU13F180 | 77.6 | 11.6 | 1 | 39.3 | Truseq | HiSeq X-Ten | Novogene | 2021 | F1 | Tahami et al. (2024) |
| | | monOU13F182 | 121.9 | 18.3 | 1 | 33.1 | Truseq | HiSeq X-Ten | Novogene | 2021 | F1 | Tahami et al. (2024) |
| | | monOU13F199 | 78.2 | 11.7 | 1 | 34.0 | Truseq | HiSeq X-Ten | Novogene | 2021 | F1 | Tahami et al. (2024) |
| | | monOU13F203 | 68.9 | 10.3 | 1 | 49.5 | Truseq | HiSeq X-Ten | Novogene | 2021 | F1 | Tahami et al. (2024) |
| | | monOU13F212 | 85.3 | 12.8 | 1 | 53.2 | Truseq | HiSeq X-Ten | Novogene | 2021 | F1 | Tahami et al. (2024) |
| | | monOU13F216 | 69.6 | 10.4 | 1 | 44.9 | Truseq | HiSeq X-Ten | Novogene | 2021 | F1 | Tahami et al. (2024) |
| | | monOU13F3 | 70.1 | 10.5 | 1 | 31.6 | Truseq | HiSeq X-Ten | Novogene | 2021 | F1 | Tahami et al. (2024) |
| | | monOU13F95 | 67.9 | 10.2 | 1 | 40.2 | Truseq | HiSeq X-Ten | Novogene | 2021 | F1 | Tahami et al. (2024) |
| | Korpilahti | monKL13F130 | 110.0 | 16.5 | 1 | 66.5 | Truseq | HiSeq X-Ten | Novogene | 2021 | F1 | Tahami et al. (2024) |
| | | monKL13F14 | 107.2 | 16.1 | 1 | 43.6 | Truseq | HiSeq X-Ten | Novogene | 2021 | F1 | Tahami et al. (2024) |
| | | monKL13F20 | 77.9 | 11.7 | 1 | 25.9 | Truseq | HiSeq X-Ten | Novogene | 2021 | F1 | Tahami et al. (2024) |
| | | monKL13F21 | 68.6 | 10.3 | 1 | 49.8 | Truseq | HiSeq X-Ten | Novogene | 2021 | F1 | Tahami et al. (2024) |
| | | monKL13F25 | 72.7 | 10.9 | 2 | 46.1 | Truseq | HiSeq X-Ten | Novogene | 2021 | F1 | Tahami et al. (2024) |
| | | monKL13F50 | 74.1 | 11.1 | 1 | 50.4 | Truseq | HiSeq X-Ten | Novogene | 2021 | F1 | Tahami et al. (2024) |
| | | monKL13F9 | 93.2 | 14.0 | 1 | 47.6 | Truseq | HiSeq X-Ten | Novogene | 2021 | F1 | Tahami et al. (2024) |
| Asia | Kamchatka | monKR1309 | 97.9 | 14.7 | 1 | 58.0 | Truseq | HiSeq X-Ten | Novogene | 2021 | ~F50 | Tahami et al. (2024) |
| | | monKR1314 | 90.6 | 13.6 | 1 | 45.1 | Truseq | HiSeq X-Ten | Novogene | 2021 | ~F50 | Tahami et al. (2024) |
| | | monKR1323 | 70.5 | 10.6 | 1 | 47.7 | Truseq | HiSeq X-Ten | Novogene | 2021 | ~F50 | Tahami et al. (2024) |

Table S9. Details of Illumina pool-sequenced *D. montana* samples (49-50 wild-caught females/males per pool), which were used in characterizing inversion frequences. The data was first published by Wiberg et al. (2021).

| Illumina pool-sequence 150bp paired-end reads<br>Region | Population | Males/females in a pool | # reads (M) | Bases (Gb) | Mean coverage | Technique | Sequencing facility | Sequencing year | Sequenced generation | First published |
|---|---|---|---|---|---|---|---|---|---|---|
| Western coast of North America | Seward | 30/20 | 406.6 | 61.0 | 233.7 | Illumina HiSeq3000 | Finnish Functional Genomics Centre (Turku, Finland) | 2016 | wild-caught | Wiberg et al. 2021 |
| | Terrace | 22/27 | 231.9 | 34.8 | 129.6 | Illumina HiSeq3000 | Finnish Functional Genomics Centre (Turku, Finland) | 2016 | wild-caught | Wiberg et al. 2021 |
| | Ashford | 16/34 | 157.1 | 23.6 | 99.4 | Illumina HiSeq3000 | Finnish Functional Genomics Centre (Turku, Finland) | 2016 | wild-caught | Wiberg et al. 2021 |
| Rocky Mountains of North America | Crested Butte | 36/13 | 204.0 | 30.6 | 116.1 | Illumina HiSeq3000 | Finnish Functional Genomics Centre (Turku, Finland) | 2016 | wild-caught | Wiberg et al. 2021 |
| Fennoscandia | Oulanka | 25/25 | 168.7 | 25.3 | 99.3 | Illumina HiSeq3000 | Finnish Functional Genomics Centre (Turku, Finland) | 2016 | wild-caught | Wiberg et al. 2021 |
| | Korpilahti | 27/23 | 179.0 | 26.9 | 106.2 | Illumina HiSeq3000 | Finnish Functional Genomics Centre (Turku, Finland) | 2016 | wild-caught | Wiberg et al. 2021 |

Table S10. *D. montana* assembly statistics. The chromosome-level genome was used as a reference genome in the genomic analyses, and the contig-level genomes of five *D. montana* populations were used to characterize inversions. N50 = half of the genome is in contigs larger than or equal to the N50 contig size. N50 count = half of the genome consists of the number indicated in N50 count. The data was first published by Poikela et al. (2024) and Tahami et al. (2024).

| Statistic | chromosome-level genome (Seward) (Poikela et al. 2024) | Seward, Alaska monSE13F37 (Poikela et al. 2024) | Jackson, USA monJX13F48 (Poikela et al. 2024) | Crested Butte, USA mon34CC5 (Tahami et al. 2024) | Oulanka, Finland monOU13F149 (Tahami et al. 2024) | Kamchatka, Russia monKR1309 (Tahami et al. 2024) |
|---|---|---|---|---|---|---|
| **Genome size (Mb)** | 145.5 | 184.3 | 181.0 | 175.0 | 178.9 | 176.7 |
| **Total contigs** | 6 | 324 | 796 | 739 | 1097 | 1675 |
| **Longest contig (Mb)** | 32.5 | 29.1 | 15.2 | 4.5 | 3.7 | 1.2 |
| **N50 (Mb)** | 26.5 | 11.0 | 1.3 | 0.8 | 0.5 | 0.2 |
| **N50 count** | 3 | 5 | 20 | 55 | 97 | 253 |
| **GC level (%)** | 40.3 | 40.2 | 40.2 | 40.2 | 40.1 | 40.1 |
| **Complete BUSCOs (%; n: 3285)** | 91.9 | 98.1 | 98.5 | 97.2 | 96.4 | 96.0 |
| **Single copy BUSCOs (%)** | 91.7 | 97.6 | 98.0 | 96.7 | 96.0 | 95.6 |
| **Duplicated BUSCOs (%)** | 0.2 | 0.5 | 0.5 | 0.5 | 0.4 | 0.4 |

Table S11. Coordinates and frequencies of the large polymorphic inversion on chromosome 4 (9.5Mb in size). The populations that carry the inversion are highlighted with grey colour.

| Coordinates: chromosome 4, 3302717 (proximal breakpoint), 12837900 (distal breakpoint) | | | | Number of individuals with | |
|---|---|---|---|---|---|
| Region | Population | Illumina sample | Genotype | the inversion (0/1 or 1/1) | Inversion frequency |
| Rocky Mountains of North America | McBride | Singly sequenced female: MB14F6 | 0/1 | 0.33 (3/9) | 0.22 (4/18) |
| | | Singly sequenced female: MB14F7 | 0/1 | | |
| | | Singly sequenced female: MB14F19 | 1/1 | | |
| | | Other singly sequenced females (6) | 0/0 | | |
| | Jackson | Singly sequenced female: JX13F3 | 0/1 | 0.33 (1/3) | 0.17 (1/6) |
| | | Other singly sequenced females (2) | 0/0 | | |
| | Crested Butte | Singly sequenced female: mon4CC5 | 1/1 | 1.00 (59/59) | 1.00 (118/118) |
| | | Singly sequenced female: mon8CC5 | 1/1 | | |
| | | Singly sequenced female: mon17CC5 | 1/1 | | |
| | | Singly sequenced female: mon26CC5 | 1/1 | | |
| | | Singly sequenced female: mon29CC5 | 1/1 | | |
| | | Singly sequenced female: mon37CC5 | 1/1 | | |
| | | Singly sequenced female: mon48CC5 | 1/1 | | |
| | | Singly sequenced female: mon68CC5 | 1/1 | | |
| | | Singly sequenced female: mon77CC5 | 1/1 | | |
| | | Singly sequenced female: mon81CC5 | 1/1 | | |
| | | Pool-sequence data (36 males, 13 females) | 1/1 | | |
| Western coast of North America | Seward | Singly sequenced females (11) | 0/0 | 0.0 (0/61) | 0.0 (0/122) |
| | | Pool-sequence data (30 males, 20 females) | 0/0 | | |
| | Terrace | Singly sequenced females (11) | 0/0 | 0.0 (0/60) | 0.0 (0/120) |
| | | Pool-sequence data (22 males, 27 females) | 0/0 | | |
| | Vancouver | Singly sequenced females (10) | 0/0 | 0.0 (0/10) | 0.0 (0/20) |
| | Ashford | Singly sequenced females (10) | 0/0 | 0.0 (0/60) | 0.0 (0/120) |
| | | Pool-sequence data (16 males, 34 females) | 0/0 | | |
| | Azalea | Singly sequenced females (3) | 0/0 | 0.0 (0/3) | 0.0 (0/6) |
| Fennoscandia | Storslett | Singly sequenced females (10) | 0/0 | 0.0 (0/10) | 0.0 (0/20) |
| | Raattama | Singly sequenced females (3) | 0/0 | 0.0 (0/3) | 0.0 (0/6) |
| | Oulanka | Singly sequenced females (11) | 0/0 | 0.0 (0/61) | 0.0 (0/122) |
| | | Pool-sequence data (25 males, 25 females) | 0/0 | | |
| | Korpilahti | Singly sequenced females (7) | 0/0 | 0.0 (0/57) | 0.0 (0/114) |
| | | Pool-sequence data (27 males, 23 females) | 0/0 | | |
| Asia | Kamchatka | Singly sequenced females (3) | 0/0 | 0.0 (0/3) | 0.0 (0/6) |

Table S12. Coordinates and frequencies of the small polymorphic inversion on chromosome 4 (~1500bp in size). The populations that carry the inversion are highlighted with grey colour.

| Coordinates: chromosome 4, 1748864 (proximal breakpoint), 1750399 (distal breakpoint) | | | | Number of individuals with | |
|---|---|---|---|---|---|
| Region | Population | Illumina sample | Genotype | the inversion (0/1 or 1/1) | Inversion frequency |
| Rocky Mountains of North America | McBride | Singly sequenced females (9) | 0/0 | 0.0 (0/9) | 0.0 (0/18) |
| | Jackson | Singly sequenced females (3) | 0/0 | 0.0 (0/3) | 0.0 (0/6) |
| | Crested Butte | Singly sequenced females (10) | 0/0 | 0.0 (0/59) | 0.0 (0/118) |
| | | Pool-sequence data (36 males, 13 females) | 0/0 | | |
| Western coast of North America | Seward | Singly sequenced females (11) | 0/0 | 0.0 (0/61) | 0.0 (0/122) |
| | | Pool-sequence data (30 males, 20 females) | 0/0 | | |
| | Terrace | Singly sequenced females (11) | 0/0 | 0.0 (0/60) | 0.0 (0/120) |
| | | Pool-sequence data (22 males, 27 females) | 0/0 | | |
| | Vancouver | Singly sequenced females (10) | 0/0 | 0.0 (0/10) | 0.0 (0/20) |
| | Ashford | Singly sequenced females (10) | 0/0 | 0.0 (0/60) | 0.0 (0/120) |
| | | Pool-sequence data (16 males, 34 females) | 0/0 | | |
| | Azalea | Singly sequenced females (3) | 0/0 | 0.0 (0/3) | 0.0 (0/6) |
| Fennoscandia | Storslett | Singly sequenced females (10) | 1/1 | 1.0 (10/10) | 1.0 (20/20) |
| | Raattama | Singly sequenced females (3) | 1/1 | 1.0 (3/3) | 1.0 (6/6) |
| | Oulanka | Singly sequenced females (11) | 1/1 | 1.0 (61/61) | 1.0 (122/122) |
| | | Pool-sequence data (25 males, 25 females) | 1/1 | | |
| | Korpilahti | Singly sequenced females (7) | 1/1 | 1.0 (57/57) | 1.0 (114/114) |
| | | Pool-sequence data (27 males, 23 females) | 1/1 | | |
| Asia | Kamchatka | Singly sequenced females (3) | 1/1 | 1.0 (3/3) | 1.0 (6/6) |

Table S13. Eigenvalues, variance (%) and cumulative variance (%) of each principal component (PC) were calculated for filtered and pruned SNPs obtained from 101 Illumina re-sequence *D. montana* samples. The analyses were performed on different data partitions: all chromosomes, the polymorphic chromosome 4 inversion, non-inverted chromosome 4, and each chromosome separately.

| Genomic region | # SNPs | | PC1 | PC2 | PC3 | PC4 | PC5 | PC6 | PC7 | PC8 | PC9 | PC10 | PC11 | PC12 | PC13 | PC14 | PC15 | PC16 | PC17 | PC18 | PC19 | PC20 |
|---|---|---|---|---|---|---|---|---|---|---|---|---|---|---|---|---|---|---|---|---|---|---|
| **All chromosomes** | 1,986,534 | **Eigenvalue** | 19.9 | 12.3 | 3.3 | 2.6 | 1.8 | 1.6 | 1.5 | 1.4 | 1.2 | 1.2 | 1.1 | 1.1 | 1.1 | 1.0 | 1.0 | 1.0 | 1.0 | 1.0 | 1.0 | 1.0 |
| | | **Variance (%)** | 34.9 | 21.6 | 5.7 | 4.6 | 3.2 | 2.8 | 2.7 | 2.4 | 2.1 | 2.0 | 2.0 | 1.9 | 1.8 | 1.8 | 1.8 | 1.8 | 1.8 | 1.8 | 1.7 | 1.7 |
| | | **Cumulative variance (%)** | 34.9 | 56.5 | 62.2 | 66.8 | 69.9 | 72.8 | 75.4 | 77.8 | 79.9 | 81.9 | 83.9 | 85.7 | 87.6 | 89.4 | 91.2 | 93.0 | 94.8 | 96.5 | 98.3 | 100.0 |
| **Chromosome 4 inversion** | 166,193 | **Eigenvalue** | 19.8 | 13.3 | 6.4 | 5.3 | 2.9 | 2.6 | 2.3 | 1.3 | 1.2 | 1.2 | 1.1 | 1.1 | 1.0 | 1.0 | 1.0 | 1.0 | 1.0 | 1.0 | 1.0 | 1.0 |
| | | **Variance (%)** | 29.8 | 20.0 | 9.6 | 8.0 | 4.3 | 4.0 | 3.4 | 2.0 | 1.8 | 1.8 | 1.7 | 1.7 | 1.6 | 1.5 | 1.5 | 1.5 | 1.5 | 1.5 | 1.5 | 1.5 |
| | | **Cumulative variance (%)** | 29.8 | 49.8 | 59.4 | 67.3 | 71.6 | 75.6 | 79.0 | 81.0 | 82.8 | 84.6 | 86.3 | 88.0 | 89.5 | 91.1 | 92.6 | 94.1 | 95.6 | 97.1 | 98.5 | 100.0 |
| **Chromosome 4 colinear** | 314,473 | **Eigenvalue** | 21.5 | 11.7 | 7.6 | 3.0 | 2.5 | 2.4 | 2.0 | 1.8 | 1.6 | 1.5 | 1.3 | 1.2 | 1.2 | 1.1 | 1.1 | 1.1 | 1.0 | 1.0 | 1.0 | 1.0 |
| | | **Variance (%)** | 32.3 | 17.5 | 11.4 | 4.5 | 3.8 | 3.6 | 3.0 | 2.8 | 2.3 | 2.2 | 2.0 | 1.9 | 1.7 | 1.7 | 1.7 | 1.6 | 1.5 | 1.5 | 1.5 | 1.5 |
| | | **Cumulative variance (%)** | 32.3 | 49.8 | 61.2 | 65.8 | 69.6 | 73.2 | 76.2 | 78.9 | 81.3 | 83.5 | 85.4 | 87.3 | 89.0 | 90.7 | 92.4 | 94.0 | 95.6 | 97.1 | 98.5 | 100.0 |
| **Chromosome 2L** | 266,896 | **Eigenvalue** | 20.0 | 12.4 | 3.0 | 1.9 | 1.7 | 1.5 | 1.4 | 1.3 | 1.1 | 1.1 | 1.1 | 1.1 | 1.0 | 1.0 | 1.0 | 1.0 | 1.0 | 1.0 | 1.0 | 1.0 |
| | | **Variance (%)** | 35.8 | 22.3 | 5.4 | 3.3 | 3.1 | 2.8 | 2.5 | 2.3 | 2.0 | 1.9 | 1.9 | 1.9 | 1.9 | 1.9 | 1.9 | 1.8 | 1.8 | 1.8 | 1.8 | 1.8 |
| | | **Cumulative variance (%)** | 35.8 | 58.1 | 63.5 | 66.9 | 70.0 | 72.8 | 75.3 | 77.6 | 79.5 | 81.5 | 83.4 | 85.3 | 87.1 | 89.0 | 90.9 | 92.7 | 94.5 | 96.4 | 98.2 | 100.0 |
| **Chromosome 2R** | 131,003 | **Eigenvalue** | 19.7 | 12.3 | 2.8 | 1.9 | 1.3 | 1.2 | 1.1 | 1.1 | 1.1 | 1.0 | 1.0 | 1.0 | 1.0 | 1.0 | 1.0 | 1.0 | 1.0 | 1.0 | 1.0 | 1.0 |
| | | **Variance (%)** | 36.7 | 23.0 | 5.3 | 3.5 | 2.4 | 2.3 | 2.0 | 2.0 | 2.0 | 2.0 | 1.9 | 1.9 | 1.9 | 1.9 | 1.9 | 1.9 | 1.9 | 1.9 | 1.9 | 1.9 |
| | | **Cumulative variance (%)** | 36.7 | 59.6 | 64.9 | 68.4 | 70.9 | 73.1 | 75.1 | 77.1 | 79.1 | 81.1 | 83.0 | 84.9 | 86.8 | 88.7 | 90.6 | 92.5 | 94.4 | 96.3 | 98.1 | 100.0 |
| **Chromosome 3** | 398,598 | **Eigenvalue** | 19.1 | 12.2 | 2.6 | 1.6 | 1.3 | 1.2 | 1.2 | 1.1 | 1.1 | 1.0 | 1.0 | 1.0 | 1.0 | 1.0 | 1.0 | 1.0 | 1.0 | 1.0 | 1.0 | 1.0 |
| | | **Variance (%)** | 36.2 | 23.1 | 5.0 | 3.0 | 2.5 | 2.4 | 2.2 | 2.1 | 2.0 | 2.0 | 2.0 | 2.0 | 2.0 | 2.0 | 2.0 | 2.0 | 2.0 | 1.9 | 1.9 | 1.9 |
| | | **Cumulative variance (%)** | 36.2 | 59.3 | 64.2 | 67.2 | 69.7 | 72.1 | 74.3 | 76.4 | 78.4 | 80.4 | 82.4 | 84.4 | 86.3 | 88.3 | 90.3 | 92.2 | 94.2 | 96.1 | 98.1 | 100.0 |
| **Chromosome 4** | 502,781 | **Eigenvalue** | 20.7 | 12.1 | 6.3 | 3.7 | 3.2 | 2.5 | 2.0 | 1.6 | 1.5 | 1.3 | 1.3 | 1.2 | 1.2 | 1.2 | 1.1 | 1.0 | 1.0 | 1.0 | 1.0 | 1.0 |
| | | **Variance (%)** | 31.4 | 18.4 | 9.6 | 5.7 | 4.8 | 3.8 | 3.0 | 2.5 | 2.3 | 2.0 | 2.0 | 1.8 | 1.8 | 1.7 | 1.7 | 1.6 | 1.5 | 1.5 | 1.5 | 1.4 |
| | | **Cumulative variance (%)** | 31.4 | 49.8 | 59.4 | 65.1 | 69.9 | 73.7 | 76.7 | 79.2 | 81.4 | 83.5 | 85.5 | 87.3 | 89.1 | 90.9 | 92.5 | 94.1 | 95.6 | 97.1 | 98.6 | 100.0 |
| **Chromosome 5** | 403,222 | **Eigenvalue** | 19.7 | 12.5 | 2.8 | 1.6 | 1.3 | 1.2 | 1.2 | 1.2 | 1.1 | 1.0 | 1.0 | 1.0 | 1.0 | 1.0 | 1.0 | 1.0 | 1.0 | 1.0 | 1.0 | 1.0 |
| | | **Variance (%)** | 36.6 | 23.3 | 5.2 | 2.9 | 2.5 | 2.3 | 2.2 | 2.1 | 2.0 | 1.9 | 1.9 | 1.9 | 1.9 | 1.9 | 1.9 | 1.9 | 1.9 | 1.9 | 1.9 | 1.9 |
| | | **Cumulative variance (%)** | 36.6 | 59.8 | 65.1 | 68.0 | 70.5 | 72.8 | 75.0 | 77.1 | 79.1 | 81.0 | 82.9 | 84.9 | 86.8 | 88.7 | 90.6 | 92.5 | 94.4 | 96.2 | 98.1 | 100.0 |
| **Chromosome X** | 284,034 | **Eigenvalue** | 21.8 | 12.6 | 6.6 | 2.8 | 1.9 | 1.8 | 1.3 | 1.2 | 1.1 | 1.1 | 1.1 | 1.1 | 1.1 | 1.1 | 1.1 | 1.1 | 1.1 | 1.1 | 1.1 | 1.1 |
| | | **Variance (%)** | 34.5 | 19.9 | 10.4 | 4.4 | 3.0 | 2.8 | 2.1 | 2.0 | 1.8 | 1.8 | 1.8 | 1.7 | 1.7 | 1.7 | 1.7 | 1.7 | 1.7 | 1.7 | 1.7 | 1.7 |
| | | **Cumulative variance (%)** | 34.5 | 54.4 | 64.8 | 69.3 | 72.2 | 75.0 | 77.1 | 79.1 | 80.9 | 82.7 | 84.4 | 86.2 | 87.9 | 89.6 | 91.4 | 93.1 | 94.8 | 96.6 | 98.3 | 100.0 |

Table S14. Mean genetic diversity (π) of *D. montana* populations. The darker (greener) the colour, the greater the value.

| | | Intergenic | | Intronic | | Coding | |
|---|---|---|---|---|---|---|---|
| **Continent** | **Region** | **Population** | **π** | **Population** | **π** | **Population** | **π** |
| **North America** | **Western coast** | Seward | 0.00923 | Seward | 0.00994 | Seward | 0.00533 |
| | | Terrace | 0.00977 | Terrace | 0.01050 | Terrace | 0.00558 |
| | | Vancouver | 0.00951 | Vancouver | 0.01022 | Vancouver | 0.00543 |
| | | Ashford | 0.00947 | Ashford | 0.01015 | Ashford | 0.00541 |
| | | Azalea | 0.00951 | Azalea | 0.01015 | Azalea | 0.00542 |
| | **Rocky Mountains** | McBride** | 0.00967 | McBride** | 0.01036 | McBride** | 0.00546 |
| | | Jackson** | 0.01005 | Jackson** | 0.01077 | Jackson** | 0.00577 |
| | | Crested Butte* | 0.00436 | Crested Butte* | 0.00463 | Crested Butte* | 0.00256 |
| **Asia** | **Kamchatka** | Kamchatka† | 0.00447 | Kamchatka† | 0.00477 | Kamchatka† | 0.00255 |
| **Europe** | **North Europe** | Storslett | 0.00679 | Storslett | 0.00723 | Storslett | 0.00400 |
| | | Raattama | 0.00675 | Raattama | 0.00719 | Raattama | 0.00397 |
| | | Oulanka | 0.00678 | Oulanka | 0.00723 | Oulanka | 0.00399 |
| | | Korpilahti | 0.00652 | Korpilahti | 0.00700 | Korpilahti | 0.00388 |

* Inversion fixed

** Inversion at lower frequency

† Samples were collected from isofemale strains kept in the laboratory for ~50 generations.

Table S15. Mean genetic diversity ($\pi$) of *D. montana* populations and mean genetic divergence ($d_{xy}$) and genetic differentiation ($f_{st}$) between Crested Butte (inversion fixed*) and the other populations (inversion absent, or at lower frequency**) in inverted and colinear regions. The darker the colour, the greater the value.

| Genetic diversity | | π INV | π COL |
|---|---|---|---|
| | Seward | 0.01151 | 0.00861 |
| | Terrace | 0.01173 | 0.00901 |
| | Vancouver | 0.01153 | 0.00882 |
| | Ashford | 0.01127 | 0.00861 |
| | Azalea | 0.01103 | 0.00844 |
| | McBride** | 0.01243 | 0.00895 |
| | Jackson** | 0.01229 | 0.00924 |
| | Crested Butte* | 0.00428 | 0.00414 |
| | Kamchatka | 0.00666 | 0.00502 |
| | Storslett | 0.00862 | 0.0062 |
| | Raattama | 0.00852 | 0.00619 |
| | Oulanka | 0.00871 | 0.00621 |
| | Korpilahti | 0.00833 | 0.00596 |
| **Genetic divergence** | | **dxy INV** | **dxy COL** |
| Crested Butte* vs | Seward | 0.01596 | 0.01102 |
| | Terrace | 0.01594 | 0.01097 |
| | Vancouver | 0.01581 | 0.01097 |
| | Ashford | 0.01583 | 0.01097 |
| | Azalea | 0.01593 | 0.01112 |
| | McBride** | 0.01502 | 0.01091 |
| | Jackson** | 0.01505 | 0.01091 |
| | Kamchatka | 0.01601 | 0.01137 |
| | Storslett | 0.01599 | 0.01143 |
| | Raattama | 0.01607 | 0.01147 |
| | Oulanka | 0.01605 | 0.01145 |
| | Korpilahti | 0.01603 | 0.01146 |
| **Genetic differentiation** | | **fst INV** | **fst COL** |
| Crested Butte* vs | Seward | 0.371 | 0.277 |
| | Terrace | 0.349 | 0.257 |
| | Vancouver | 0.350 | 0.266 |
| | Ashford | 0.356 | 0.267 |
| | Azalea | 0.343 | 0.274 |
| | McBride** | 0.317 | 0.259 |
| | Jackson** | 0.302 | 0.244 |
| | Kamchatka | 0.527 | 0.471 |
| | Storslett | 0.430 | 0.380 |
| | Raattama | 0.436 | 0.384 |
| | Oulanka | 0.428 | 0.383 |
| | Korpilahti | 0.442 | 0.392 |

Table S16. BayeScEnv was used to identify loci associated with environmental variables (PC1 and PC2) and cold phenotypes ($CT_{min}$, CCRT). The number and proportion of significant SNPs associated with a given variable were obtained from significance level 0.05 and are shown separately for each chromosome and chromosome arm, and for the large inversion on chromosome 4 (4INV = inversion using the strict inversion breakpoints, 4INV+1.5Mb = inversion extends 1.5Mb beyond the breakpoints into the colinear regions), and for colinear regions (COL = colinear regions using the strict inversion breakpoints, COL-1.5Mb = colinear regions except 1.5Mb regions beyond the breakpoints that were included as part of the inversion).

| Variable | Genomic region | # Significant SNPs | # Non-significant SNPs | # SNPs total | % Significant SNPs |
|---|---|---|---|---|---|
| **PC1** | **All** | 66852 | 3049251 | 3546748 | 1.88 |
| | **2L** | 6131 | 430645 | 436776 | 1.40 |
| | **2R** | 2955 | 224029 | 226984 | 1.30 |
| | **3** | 5186 | 682366 | 687552 | 0.75 |
| | **4** | 15658 | 965111 | 980769 | 1.60 |
| | **5** | 9033 | 648757 | 657790 | 1.37 |
| | **X** | 27889 | 528988 | 556877 | 5.01 |
| | **4INV** | 6181 | 311867 | 318048 | 1.94 |
| | **COL** | 60671 | 3168029 | 3228700 | 1.88 |
| | **4INV+1.5Mb** | 9095 | 378563 | 387658 | 2.35 |
| | **COL-1.5Mb** | 57757 | 3101333 | 3159090 | 1.83 |
| **PC2** | **All** | 34481 | 3075920 | 3545139 | 0.97 |
| | **2L** | 1766 | 434738 | 436504 | 0.40 |
| | **2R** | 804 | 226105 | 226909 | 0.35 |
| | **3** | 1709 | 685594 | 687303 | 0.25 |
| | **4** | 9188 | 971395 | 980583 | 0.94 |
| | **5** | 2205 | 655236 | 657441 | 0.34 |
| | **X** | 18809 | 537590 | 556399 | 3.38 |
| | **4INV** | 5501 | 312525 | 318026 | 1.73 |
| | **COL** | 28980 | 3198133 | 3227113 | 0.90 |
| | **4INV+1.5Mb** | 7746 | 379849 | 387595 | 2.00 |
| | **COL-1.5Mb** | 26735 | 3130809 | 3157544 | 0.85 |
| **CTmin** | **All** | 54789 | 3051662 | 3538127 | 1.55 |
| | **2L** | 4015 | 431676 | 435691 | 0.92 |
| | **2R** | 1802 | 224731 | 226533 | 0.80 |
| | **3** | 3345 | 683367 | 686712 | 0.49 |
| | **4** | 15267 | 963093 | 978360 | 1.56 |
| | **5** | 5004 | 651129 | 656133 | 0.76 |
| | **X** | 25356 | 529342 | 554698 | 4.57 |
| | **4INV** | 2802 | 314419 | 317221 | 0.88 |
| | **COL** | 51987 | 3168919 | 3220906 | 1.61 |
| | **4INV+1.5Mb** | 4380 | 382076 | 386456 | 1.13 |
| | **COL-1.5Mb** | 50409 | 3101262 | 3151671 | 1.60 |
| **CCRT** | **All** | 32464 | 3078639 | 3546464 | 0.92 |
| | **2L** | 1396 | 435361 | 436757 | 0.32 |
| | **2R** | 776 | 226190 | 226966 | 0.34 |
| | **3** | 1235 | 686300 | 687535 | 0.18 |
| | **4** | 6374 | 974456 | 980830 | 0.65 |
| | **5** | 1894 | 655863 | 657757 | 0.29 |
| | **X** | 20789 | 535830 | 556619 | 3.73 |
| | **4INV** | 905 | 317301 | 318206 | 0.28 |
| | **COL** | 31559 | 3196699 | 3228258 | 0.98 |
| | **4INV+1.5Mb** | 1494 | 386358 | 387852 | 0.39 |
| | **COL-1.5Mb** | 30970 | 3127642 | 3158612 | 0.98 |

Table S17. Chi Square ($X^2$) test was used to analyse BayeScEnv results between the large inversion on chromosome 4 (4INV = inversion using the strict inversion breakpoints, 4INV+1.5Mb = inversion extends 1.5Mb beyond the breakpoints into the colinear regions) and colinear regions (COL = colinear regions using the strict inversion breakpoints, COL-1.5Mb = colinear regions except 1.5Mb regions beyond the breakpoints that were included as part of the inversion). Significant P-values are bolded, and significant comparisons in the expected direction (4INV>COL) are highlighted with grey shading.

| Chi Square test | | | | | | |
|---|---|---|---|---|---|---|
| | Genomic region | # Significant SNPs | # Non-significant SNPs | % significant SNPs | $X^2$ | P-value |
| PC1 | COL | 60671 | 3168029 | 1.88 | 6.47 | **0.011** |
| | 4INV | 6181 | 311867 | 1.94 | | |
| | COL-1.5Mb | 57757 | 3101333 | 1.83 | 500.71 | **<0.001** |
| | 4INV+1.5Mb | 9095 | 378563 | 2.35 | | |
| PC2 | COL | 28980 | 3198133 | 0.90 | 2079.18 | **<0.001** |
| | 4INV | 5501 | 312525 | 1.73 | | |
| | COL-1.5Mb | 26735 | 3130809 | 0.85 | 4754.75 | **<0.001** |
| | 4INV+1.5Mb | 7746 | 379849 | 2.00 | | |
| CTmin | COL | 51987 | 3168919 | 1.61 | 1011.50 | **<0.001** |
| | 4INV | 2802 | 314419 | 0.88 | | |
| | COL-1.5Mb | 50409 | 3101262 | 1.60 | 490.47 | **<0.001** |
| | 4INV+1.5Mb | 4380 | 382076 | 1.13 | | |
| CCRT | COL | 31559 | 3196699 | 0.98 | 1534.47 | **<0.001** |
| | 4INV | 905 | 317301 | 0.28 | | |
| | COL-1.5Mb | 30970 | 3127642 | 0.98 | 1349.64 | **<0.001** |
| | 4INV+1.5Mb | 1494 | 386358 | 0.39 | | |

Table S18. The table shows the number of genes that were associated with the significant SNPs linked to climate variability (PC1, PC2) and cold tolerance ($CT_{min}$, CCRT) within the inversion (Table S17, full lists of genes in Supplementary file 1). It also lists the GO-terms in which the genes showed a significant enrichment. We used orthologs from both *D. virilis* and *D. melanogaster*, as *D. virilis* and *D. montana* are more closely related, but *D. melanogaster* has superior annotation. 4INV indicates the genes located within the inversion using the strict inversion breakpoints, and 4INV+1.5Mb the genes located within the inversion or within 1.5Mb regions beyond the breakpoints into the colinear regions.

| Variable | Region | # genes | GO-term | Species | Benjamini corrected P-value |
|---|---|---|---|---|---|
| PC1 | 4INV | 392 | EGF-like_domain | *D. virilis* | 1.12E-02 |
| | | | EGF | *D. virilis* | 2.66E-02 |
| | | | EGF-like_domain | *D. melanogaster* | 2.82E-05 |
| | | | membrane | *D. melanogaster* | 1.78E-04 |
| | | | EGF | *D. melanogaster* | 4.00E-04 |
| | | | phagocytosis | *D. melanogaster* | 2.19E-03 |
| | | | EGF-like_dom | *D. melanogaster* | 3.95E-03 |
| | | | voltage-gated calcium channel complex | *D. melanogaster* | 2.98E-02 |
| | | | DOMAIN:EGF-like | *D. melanogaster* | 3.45E-02 |
| | | | sodium ion transport | *D. melanogaster* | 4.02E-02 |
| | | | Transmembrane | *D. melanogaster* | 4.05E-02 |
| | | | DOMAIN:Ig-like | *D. melanogaster* | 4.43E-02 |
| | | | Signal-anchor | *D. melanogaster* | 4.51E-02 |
| PC1 | 4INV+1.5Mb | 507 | EGF-like_domain | *D. melanogaster* | 1.68E-04 |
| | | | phagocytosis | *D. melanogaster* | 1.12E-03 |
| | | | EGF | *D. melanogaster* | 5.25E-03 |
| | | | EGF-like_dom | *D. melanogaster* | 4.61E-02 |
| PC2 | 4INV | 332 | EGF-like_domain | *D. virilis* | 1.10E-04 |
| | | | EGF | *D. virilis* | 1.15E-03 |
| | | | EGF-like_dom | *D. virilis* | 6.81E-03 |
| | | | IG | *D. virilis* | 2.44E-02 |
| | | | Ig-like_dom | *D. virilis* | 3.84E-02 |
| | | | Ig-like_dom_sf | *D. virilis* | 3.84E-02 |
| | | | Neural/epithelial_adhesion | *D. virilis* | 3.84E-02 |
| | | | Ig_sub | *D. virilis* | 3.84E-02 |
| | | | DOMAIN:EGF-like | *D. virilis* | 4.75E-02 |
| | | | DOMAIN:Ig-like | *D. virilis* | 4.75E-02 |
| | | | EGF-like_domain | *D. melanogaster* | 8.22E-08 |
| | | | EGF | *D. melanogaster* | 9.44E-06 |
| | | | phagocytosis | *D. melanogaster* | 2.21E-05 |
| | | | EGF-like_dom | *D. melanogaster* | 6.37E-05 |
| | | | membrane | *D. melanogaster* | 4.91E-04 |
| | | | DOMAIN:EGF-like | *D. melanogaster* | 9.84E-04 |
| | | | Growth_fac_rcpt_cys_sf | *D. melanogaster* | 1.28E-02 |
| | | | DOMAIN:Ig-like | *D. melanogaster* | 4.13E-02 |
| | | | Ig-like_dom_sf | *D. melanogaster* | 4.26E-02 |
| | | | Ig-like_dom | *D. melanogaster* | 4.26E-02 |
| | | | Neural/epithelial_adhesion | *D. melanogaster* | 4.26E-02 |
| PC2 | 4INV+1.5Mb | 423 | EGF-like_domain | *D. virilis* | 6.32E-04 |
| | | | EGF | *D. virilis* | 1.16E-02 |
| | | | EGF-like_domain | *D. melanogaster* | 6.15E-07 |
| | | | membrane | *D. melanogaster* | 9.56E-05 |
| | | | EGF | *D. melanogaster* | 1.66E-04 |
| | | | phagocytosis | *D. melanogaster* | 1.87E-04 |
| | | | EGF-like_dom | *D. melanogaster* | 9.83E-04 |
| | | | Growth_fac_rcpt_cys_sf | *D. melanogaster* | 6.70E-03 |
| | | | DOMAIN:EGF-like | *D. melanogaster* | 1.03E-02 |

| Variable | Region | # genes | GO-term | Species | Benjamini corrected P-value |
|---|---|---|---|---|---|
| $CT_{min}$ | 4INV | 307 | EGF | *D. virilis* | 2.67E-02 |
| | | | IG | *D. virilis* | 4.88E-02 |
| | | | membrane | *D. melanogaster* | 5.61E-05 |
| | | | Transmembrane | *D. melanogaster* | 1.99E-03 |
| | | | Ion channel | *D. melanogaster* | 2.62E-03 |
| | | | EGF | *D. melanogaster* | 3.49E-03 |
| | | | voltage-gated calcium channel complex | *D. melanogaster* | 1.13E-02 |
| | | | TRANSMEM:Helical | *D. melanogaster* | 2.70E-02 |
| | | | DOMAIN:EGF-like | *D. melanogaster* | 2.70E-02 |
| $CT_{min}$ | 4INV+1.5Mb | 400 | membrane | *D. melanogaster* | 1.95E-05 |
| | | | Transmembrane | *D. melanogaster* | 3.64E-03 |
| | | | voltage-gated calcium channel complex | *D. melanogaster* | 3.15E-02 |
| | | | EGF | *D. melanogaster* | 3.16E-02 |
| | | | TRANSMEM:Helical | *D. melanogaster* | 3.32E-02 |
| CCRT | 4INV | 146 | voltage-gated calcium channel complex | *D. virilis* | 2.84E-03 |
| | | | Calcium transport | *D. virilis* | 3.92E-03 |
| | | | FA58C | *D. virilis* | 3.00E-02 |
| | | | IG | *D. virilis* | 3.00E-02 |
| | | | DOMAIN:F5/8 type C | *D. virilis* | 3.39E-02 |
| | | | DOMAIN:Ig-like | *D. virilis* | 3.39E-02 |
| | | | TRANSMEM:Helical | *D. virilis* | 3.39E-02 |
| | | | Transmembrane | *D. virilis* | 3.86E-02 |
| | | | Membrane | *D. virilis* | 4.71E-02 |
| | | | membrane | *D. melanogaster* | 2.47E-04 |
| | | | Transmembrane | *D. melanogaster* | 4.19E-04 |
| | | | voltage-gated calcium channel complex | *D. melanogaster* | 7.58E-04 |
| | | | TRANSMEM:Helical | *D. melanogaster* | 1.03E-03 |
| | | | voltage-gated calcium channel activity | *D. melanogaster* | 4.83E-03 |
| | | | Receptor | *D. melanogaster* | 9.74E-03 |
| | | | Transmembrane helix | *D. melanogaster* | 1.41E-02 |
| | | | Membrane | *D. melanogaster* | 1.71E-02 |
| | | | Calcium transport | *D. melanogaster* | 1.72E-02 |
| | | | EGF | *D. melanogaster* | 3.40E-02 |
| | | | FA58C | *D. melanogaster* | 3.51E-02 |
| CCRT | 4INV+1.5Mb | 186 | Calcium transport | *D. virilis* | 1.20E-02 |
| | | | voltage-gated calcium channel complex | *D. virilis* | 4.80E-03 |
| | | | Transmembrane | *D. virilis* | 9.63E-04 |
| | | | TRANSMEM:Helical | *D. virilis* | 6.57E-03 |
| | | | DOMAIN:Sodium/calcium exchanger mem | *D. virilis* | 1.62E-02 |
| | | | Antiport | *D. virilis* | 2.36E-02 |
| | | | DOMAIN:F5/8 type C | *D. virilis* | 3.8E-02 |
| | | | Transmembrane | *D. melanogaster* | 7.62E-05 |
| | | | membrane | *D. melanogaster* | 4.46E-04 |
| | | | TRANSMEM:Helical | *D. melanogaster* | 6.83E-04 |
| | | | voltage-gated calcium channel complex | *D. melanogaster* | 2.01E-03 |
| | | | Calcium transport | *D. melanogaster* | 4.54E-03 |
| | | | Receptor | *D. melanogaster* | 7.62E-03 |
| | | | Transmembrane helix | *D. melanogaster* | 7.67E-03 |
| | | | voltage-gated calcium channel activity | *D. melanogaster* | 1.19E-02 |

Table S19. Support, measured as ΔlnCL, and parameter estimates for divergence time (*T* in years/generations), migration rate (*m*) and effective population sizes ($N_e$) for three *D. montana* populations and their common ancestral population under strict divergence model (DIV, *m*=0) and isolation with migration models with both gene flow directions ($IM_{A\to B}$ or $IM_{B\to A}$). Migration rate (*m*) estimates correspond to M (=$4N_e m$) individuals per generation (forwards in time). The model comparisons are based on 256bp blocks and were performed for colinear non-repetitive intergenic regions to minimize the direct effects of selection. Grey shading indicates the best-fit model for each comparison (see Fig. S12).

| Population | Model | Ancestral *Ne* | Pop. *A Ne* | Pop. *B Ne* | T (years) | *m (M = 4Nem)* | ΔlnCL |
|---|---|---|---|---|---|---|---|
| Crested Butte (A) - Vancouver (B) | DIV | 620,765 | 205,734 | 1,181,914 | 594,424 | - | 1,093,392 |
| | IM *D. monA --> D. monB* | 617,355 | 207,830 | 1,158,007 | 610,170 | 2.60E-08 (0.12) | 1,089,932 |
| | IM *D. monB --> D. monA* | 438,089 | 147,816 | 1,070,603 | 958,812 | 4.95E-07 (0.29) | 0 |
| Crested Butte (A) - Storslett (B) | DIV | 601,013 | 228,554 | 437,696 | 729,495 | - | 646,726 |
| | IM *D. monA --> D. monB* | 586,550 | 232,831 | 431,550 | 772,769 | 3.92E-08 (0.07) | 615,447 |
| | IM *D. monB --> D. monA* | 474,766 | 198,026 | 471,474 | 1,016,841 | 2.19E-07 (0.17) | 0 |
| Storslett (A) - Vancouver (B) | DIV | 527,819 | 413,484 | 1,228,026 | 651,134 | - | 403,580 |
| | IM *D. monA --> D. monB* | 460,630 | 432,335 | 917,162 | 899,273 | 3.88E-07 (1.42) | 17,628 |
| | IM *D. monB --> D. monA* | 441,187 | 311,827 | 1,096,621 | 861,097 | 4.83E-07 (0.60) | 0 |

Table S20. Barrier regions (Mb) and barrier fraction (%) for different chromosomes, for the large inversion on chromosome 4 (4INV = inversion using the strict inversion breakpoints, 4INV+1.5Mb = inversion extends 1.5Mb beyond the breakpoints into the colinear regions), and for colinear regions (COL = colinear regions using the strict inversion breakpoints, COL-1.5Mb = colinear regions except 1.5Mb regions beyond the breakpoints that were included as part of the inversion).

| Genomic region | Barrier (Mb) | Total (Mb) | Barrier fraction (%) |
|---|---|---|---|
| **All** | 15.57 | 145.45 | 10.71 |
| **X** | 0.32 | 29.14 | 1.09 |
| **2L** | 1.28 | 20.25 | 6.32 |
| **2R** | 0.78 | 11.00 | 7.12 |
| **3** | 1.33 | 26.02 | 5.11 |
| **4** | 8.20 | 32.54 | 25.19 |
| **5** | 3.67 | 26.51 | 13.84 |
| **4INV** | 6.00 | 9.54 | 62.91 |
| **4INV + 1.5Mb** | 6.83 | 12.54 | 54.52 |
| **COL** | 9.58 | 135.92 | 7.05 |
| **COL - 1.5MB** | 8.74 | 132.92 | 6.58 |

Table S21. A Chi-Square ($X^2$) test was used to estimate barrier enrichment in inverted regions (4INV = inversion using the strict inversion breakpoints, 4INV+1.5Mb = inversion extends 1.5Mb beyond the breakpoints into the colinear regions) compared to colinear regions (COL = colinear regions using the strict inversion breakpoints, COL-1.5Mb = colinear regions except 1.5Mb regions beyond the breakpoints that were included as part of the inversion). The test was based on the number of non-overlapping barrier and non-barrier windows (divided by five to obtain non-overlapping windows) in each region.

| Chi-square test | | #barrier windows | #non-barrier windows | $X^2$-statistic | P-value |
|---|---|---|---|---|---|
| **Genomic region** | **4INV** | 45.6 | 62 | 309.9 | **<0.0001** |
| | **COL** | 47.6 | 1621.6 | | |
| | **4INV+1.5MB** | 51.2 | 73.8 | 337.4 | **<0.0001** |
| | **COL-1.5MB** | 42 | 1609.8 | | |

### Supplementary Figures

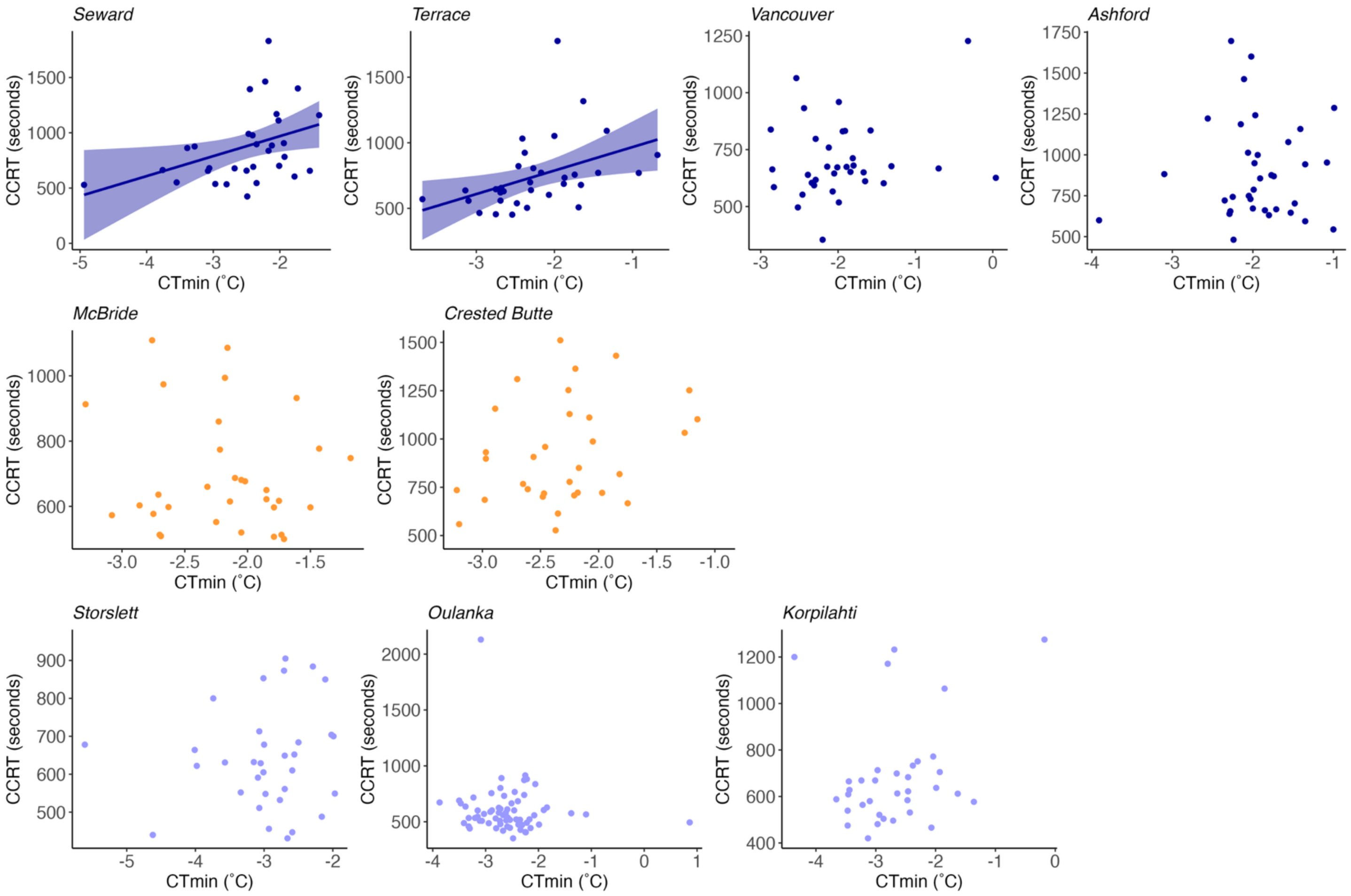

Figure S1. Pearson correlation between $CT_{min}$ and CCRT in nine *D. montana* populations (black dashed line). Regression lines indicate significant correlations. The respective statistics are shown in Table S6.

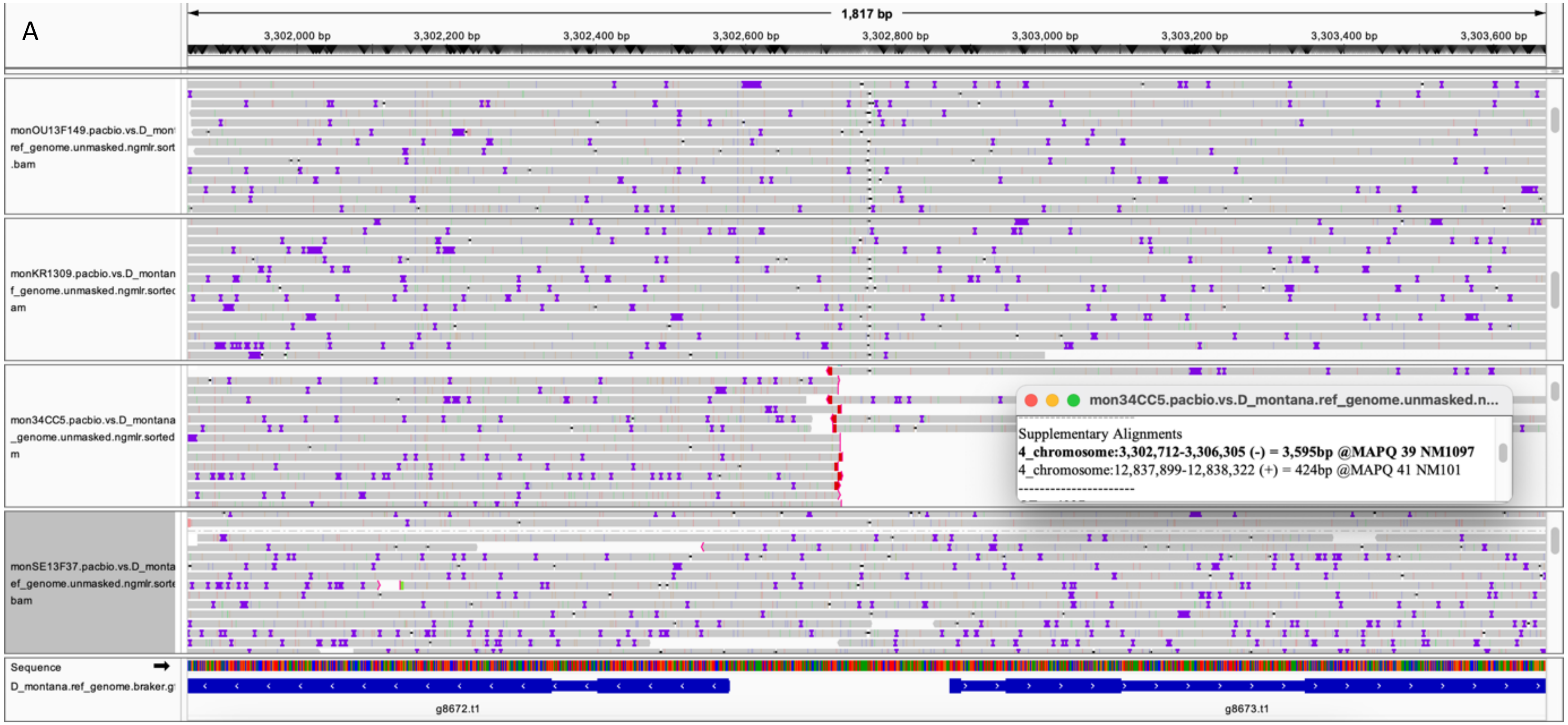
A
1,817 bp
3,302,000 bp
3,302,200 bp
3,302,400 bp
3,302,600 bp
3,302,800 bp
3,303,000 bp
3,303,200 bp
3,303,400 bp
3,303,600 bp
monOU13F149.pacbio.vs.D_mon
ref_genome.unmasked.ngmlr.sort
.bam
monKR1309.pacbio.vs.D_montan
f_genome.unmasked.ngmlr.sortec
am
mon34CC5.pacbio.vs.D_montana
_genome.unmasked.ngmlr.sorted
m
monSE13F37.pacbio.vs.D_monta
ef_genome.unmasked.ngmlr.sorte
bam
mon34CC5.pacbio.vs.D_montana.ref_genome.unmasked.n...
Supplementary Alignments
4_chromosome:3,302,712-3,306,305 (-) = 3,595bp @MAPQ 39 NM1097
4_chromosome:12,837,899-12,838,322 (+) = 424bp @MAPQ 41 NM101
Sequence
D_montana.ref_genome.braker.g
g8672.t1
g8673.t1

B

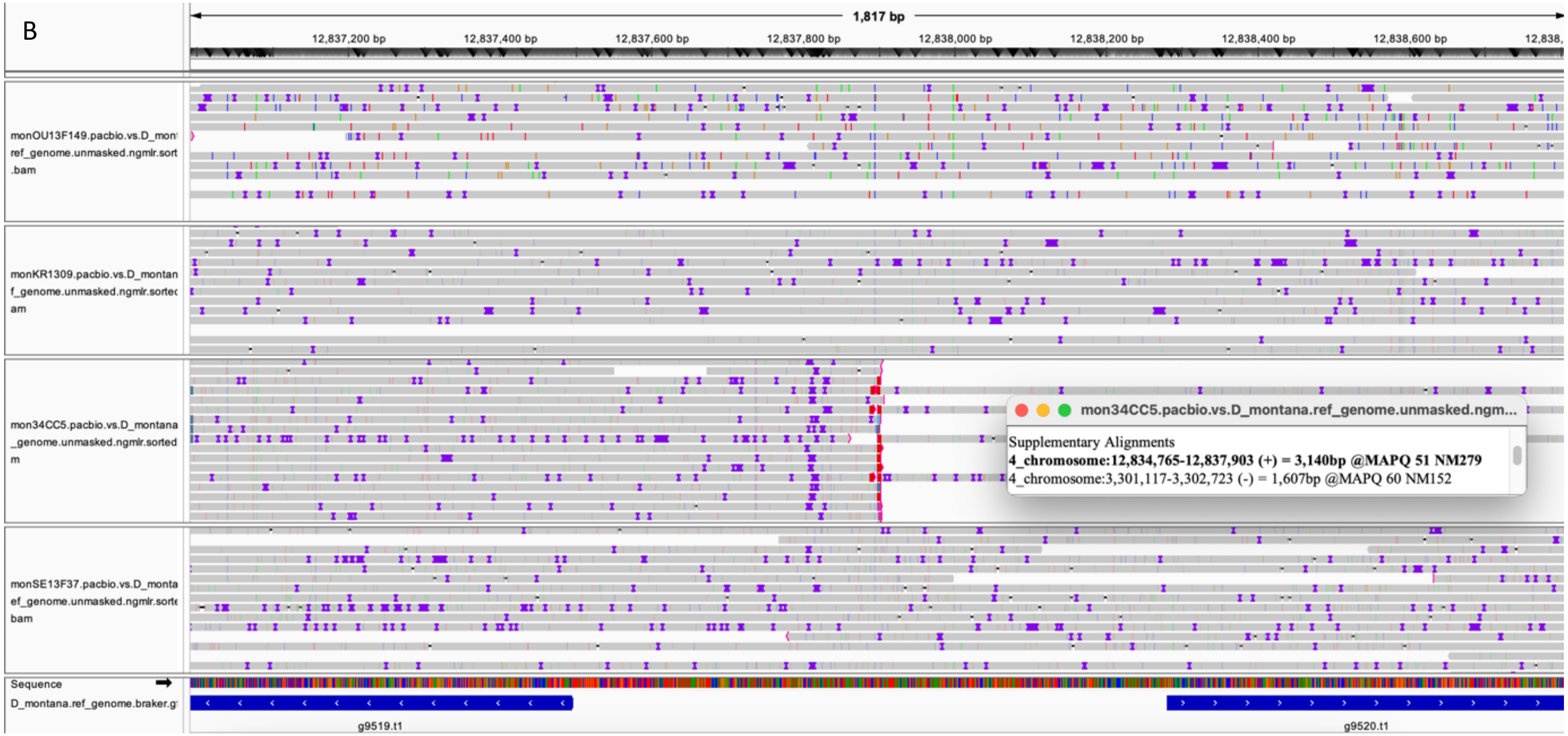
1,817 bp
12,837,200 bp
12,837,400 bp
12,837,600 bp
12,837,800 bp
12,838,000 bp
12,838,200 bp
12,838,400 bp
12,838,600 bp
monOU13F149.pacbio.vs.D_mon ref_genome.unmasked.ngmlr.sort .bam
monKR1309.pacbio.vs.D_montan f_genome.unmasked.ngmlr.sortec am
mon34CC5.pacbio.vs.D_montana _genome.unmasked.ngmlr.sorted m
mon34CC5.pacbio.vs.D_montana.ref_genome.unmasked.ngm...
Supplementary Alignments
4_chromosome:12,834,765-12,837,903 (+) = 3,140bp @MAPQ 51 NM279
4_chromosome:3,301,117-3,302,723 (-) = 1,607bp @MAPQ 60 NM152
monSE13F37.pacbio.vs.D_monta ef_genome.unmasked.ngmlr.sorte bam
Sequence
D_montana.ref_genome.braker.g
g9519.t1
g9520.t1

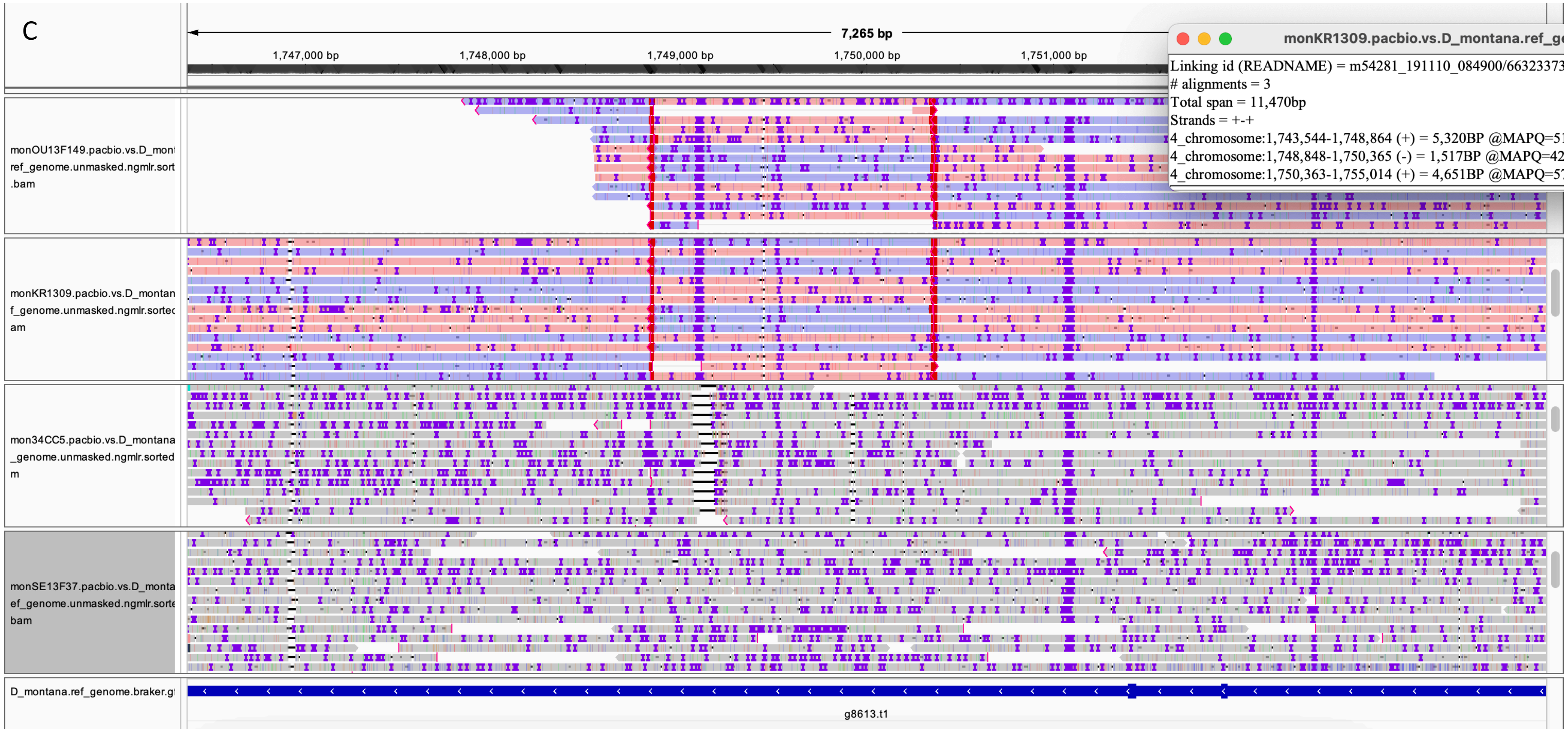

Figure S2. Inversion breakpoints illustrated with Integrative Genomics Viewer (IGV) v2.18.0 (Thorvaldsdottir et al. 2013). PacBio reads from different samples were mapped against the *D. montana* reference genome (Seward). (A) Proximal and (B) distal breakpoints of the large polymorphic inversion on chromosome 4. This inversion is fixed in the Crested Butte population (mon34CC5), as indicated by split reads (red tip of the read) and reversed orientation relative to the reference genome. (C) The small polymorphic inversion on chromosome 4, which is fixed in Fennoscandian and Asian populations (monOU13F149, monKR1309), shows split and inverted reads, with the inverted region shown in a different color compared to the rest of the reads. Gene IDs: g8672.t1 = tRNA-specific adenosine deaminase 1, g8673.t1 = UPF0428 protein CG16865, g9519.t1 = sorting nexin-17, g9520.t1 = tRNA-dihydrouridine(20a/20b) synthase [NAD(P)+]-like, g8613.t1 = leucine-rich repeat transmembrane neuronal protein 3.

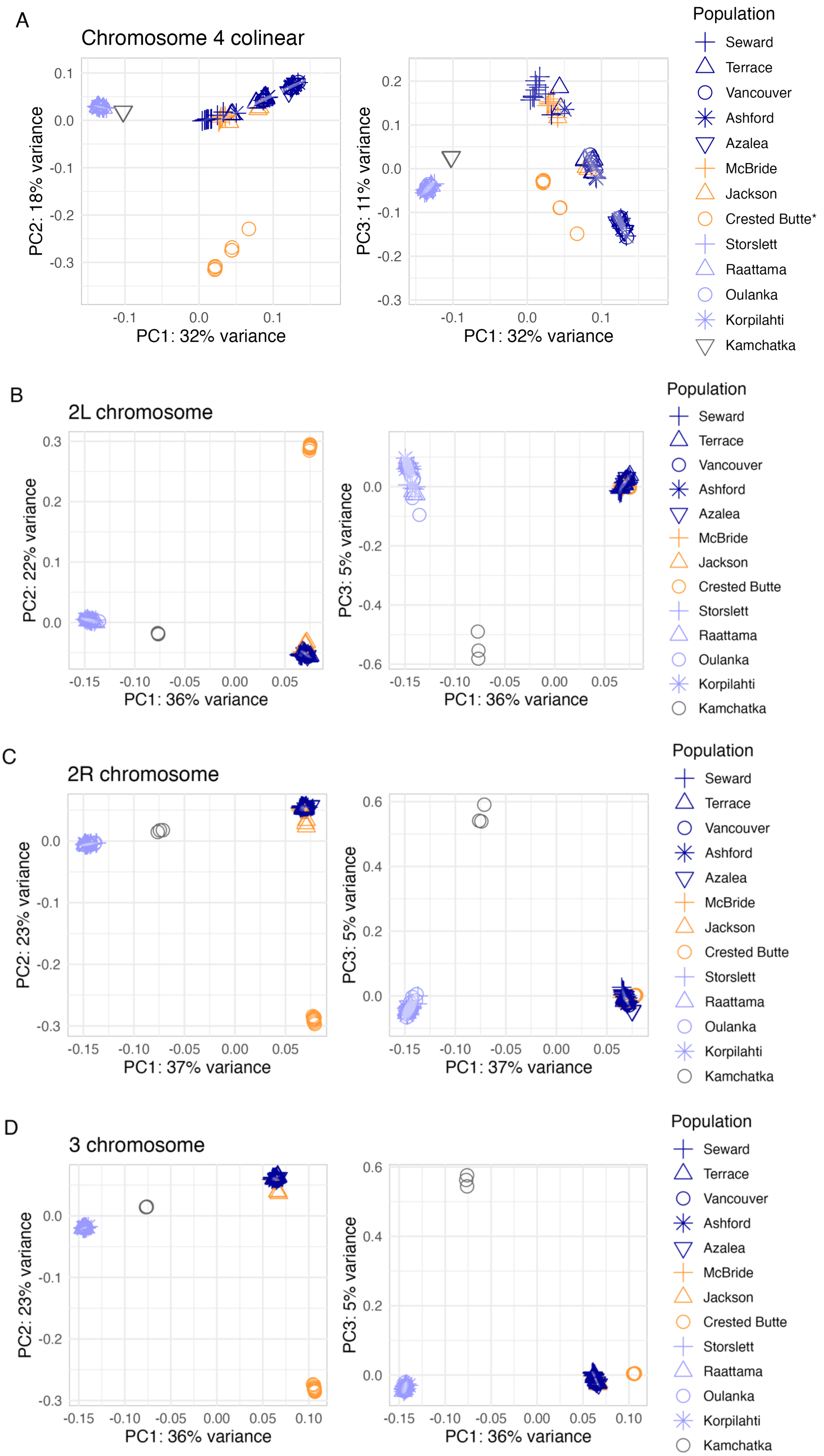
A
Chromosome 4 colinear
PC2: 18% variance
PC1: 32% variance
PC3: 11% variance
Population
Seward
Terrace
Vancouver
Ashford
Azalea
McBride
Jackson
Crested Butte*
Storslett
Raattama
Oulanka
Korpilahti
Kamchatka
B
2L chromosome
PC2: 22% variance
PC1: 36% variance
PC3: 5% variance
Crested Butte
C
2R chromosome
PC2: 23% variance
PC1: 37% variance
PC3: 5% variance
D
3 chromosome
PC2: 23% variance
PC1: 36% variance
PC3: 5% variance

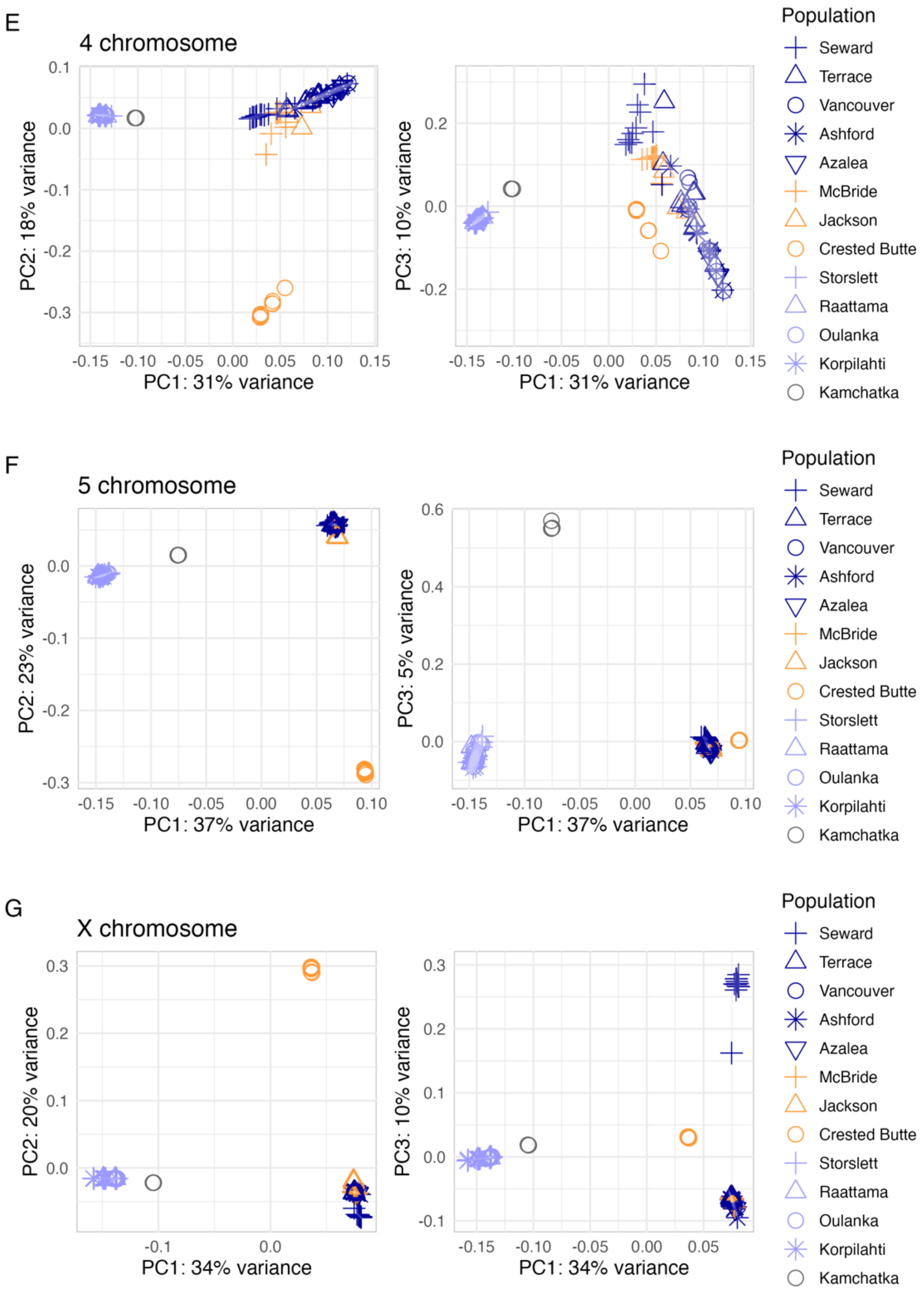

Figure S3. Genomic variation (SNPs from coding and non-coding regions) among *D. montana* populations (3-11 Illumina samples/population) summarized in a principal component analysis (PCA) for (A) colinear chromosome 4, (B) left arm of chromosome 2 (2L), (C) right arm of chromosome 2 (2R), (D) chromosome 3, (E) chromosome 4, (F) chromosome 5, and (G) the X chromosome. See details in Table S13.

A

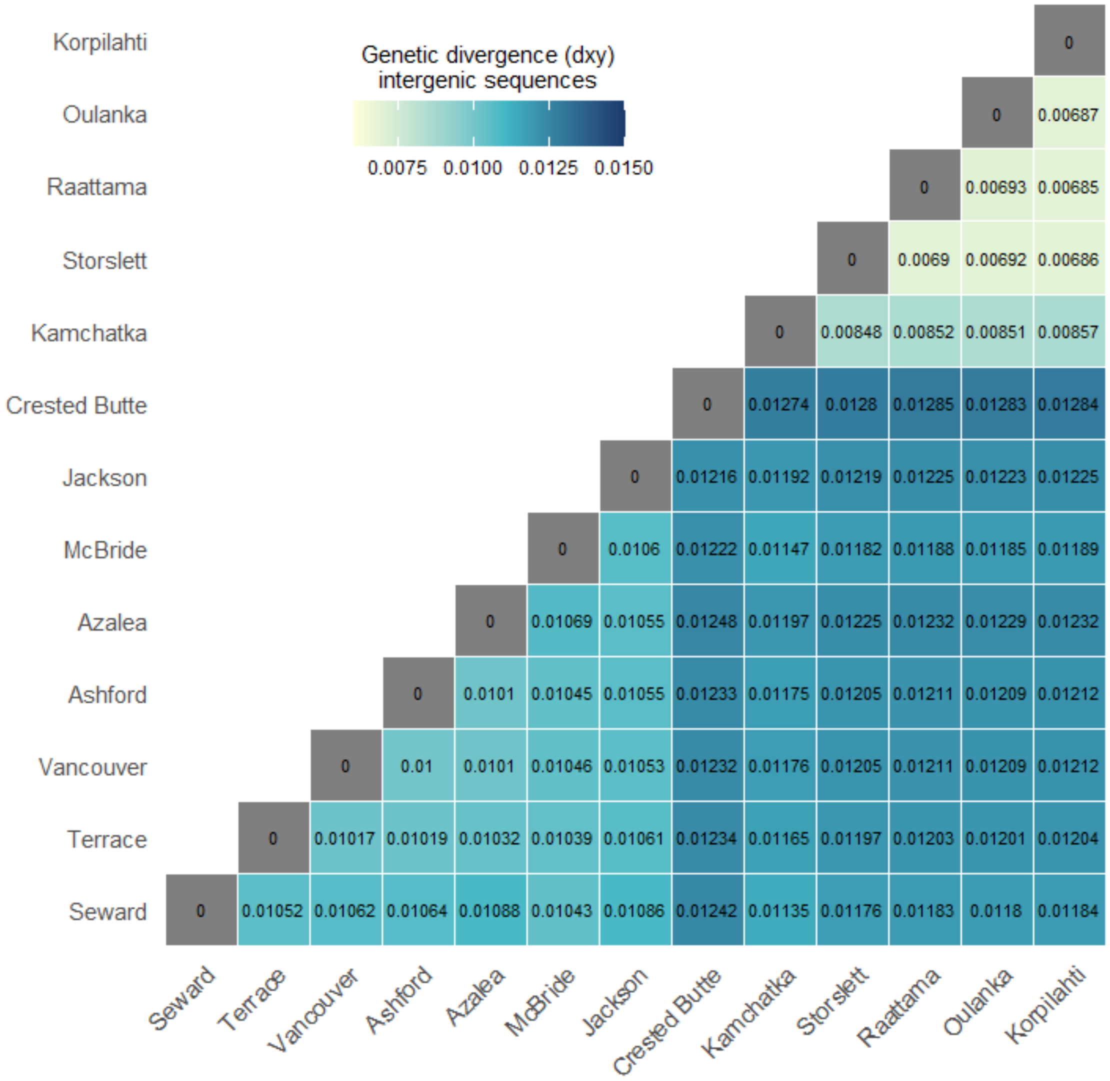
Genetic divergence (dxy)
intergenic sequences
0.0075 0.0100 0.0125 0.0150
Korpilahti
Oulanka
Raattama
Storslett
Kamchatka
Crested Butte
Jackson
McBride
Azalea
Ashford
Vancouver
Terrace
Seward
0
0 0.00687
0 0.00693 0.00685
0 0.0069 0.00692 0.00686
0 0.00848 0.00852 0.00851 0.00857
0 0.01274 0.0128 0.01285 0.01283 0.01284
0 0.01216 0.01192 0.01219 0.01225 0.01223 0.01225
0 0.0106 0.01222 0.01147 0.01182 0.01188 0.01185 0.01189
0 0.01069 0.01055 0.01248 0.01197 0.01225 0.01232 0.01229 0.01232
0 0.0101 0.01045 0.01055 0.01233 0.01175 0.01205 0.01211 0.01209 0.01212
0 0.01 0.0101 0.01046 0.01053 0.01232 0.01176 0.01205 0.01211 0.01209 0.01212
0 0.01017 0.01019 0.01032 0.01039 0.01061 0.01234 0.01165 0.01197 0.01203 0.01201 0.01204
0 0.01052 0.01062 0.01064 0.01088 0.01043 0.01086 0.01242 0.01135 0.01176 0.01183 0.0118 0.01184
Seward
Terrace
Vancouver
Ashford
Azalea
McBride
Jackson
Crested Butte
Kamchatka
Storslett
Raattama
Oulanka
Korpilahti

B

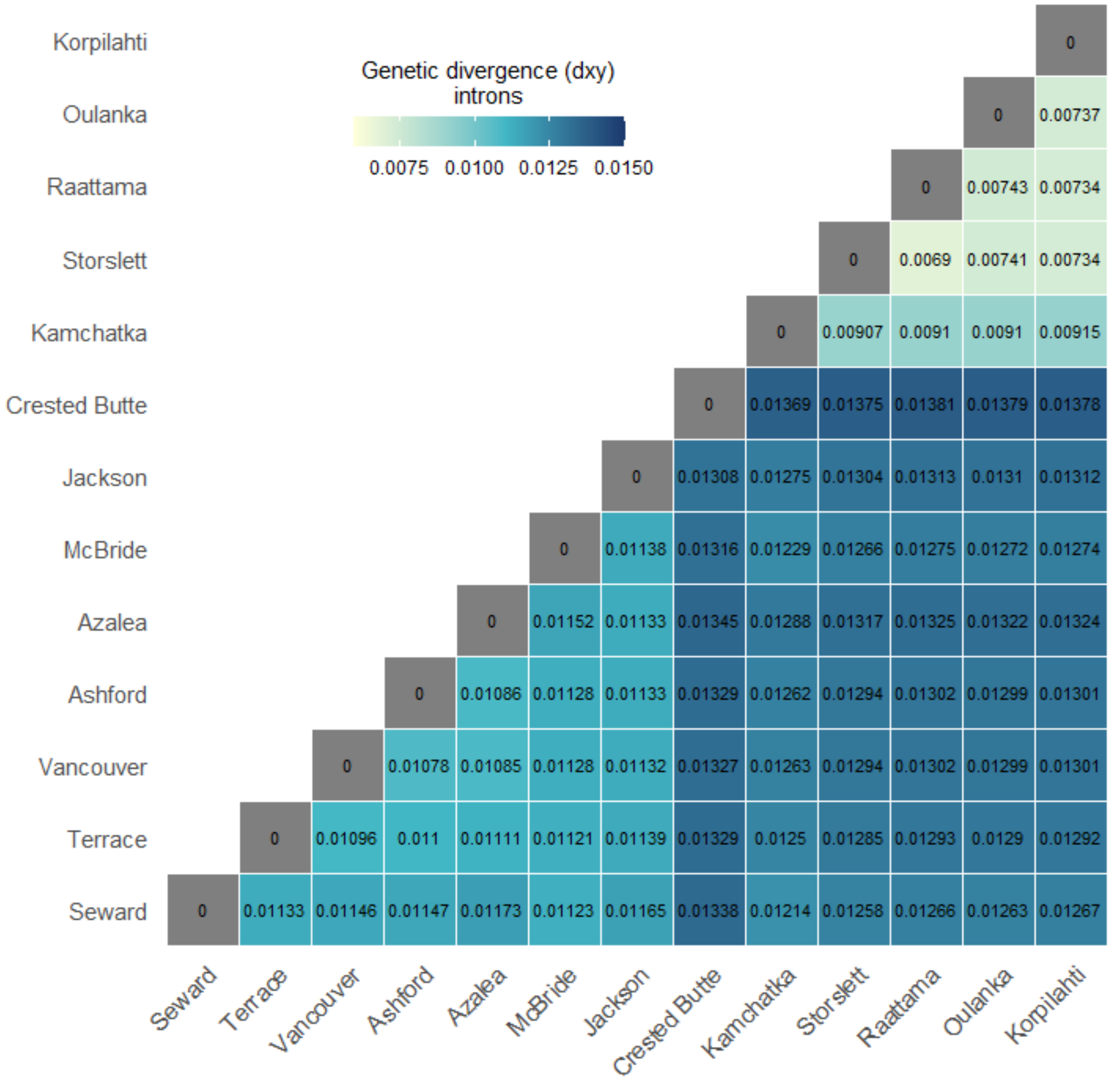
Genetic divergence (dxy)
introns
0.0075 0.0100 0.0125 0.0150
Korpilahti
Oulanka
Raattama
Storslett
Kamchatka
Crested Butte
Jackson
McBride
Azalea
Ashford
Vancouver
Terrace
Seward
0
0 0.00737
0 0.00743 0.00734
0 0.0069 0.00741 0.00734
0 0.00907 0.0091 0.0091 0.00915
0 0.01369 0.01375 0.01381 0.01379 0.01378
0 0.01308 0.01275 0.01304 0.01313 0.0131 0.01312
0 0.01138 0.01316 0.01229 0.01266 0.01275 0.01272 0.01274
0 0.01152 0.01133 0.01345 0.01288 0.01317 0.01325 0.01322 0.01324
0 0.01086 0.01128 0.01133 0.01329 0.01262 0.01294 0.01302 0.01299 0.01301
0 0.01078 0.01085 0.01128 0.01132 0.01327 0.01263 0.01294 0.01302 0.01299 0.01301
0 0.01096 0.011 0.01111 0.01121 0.01139 0.01329 0.0125 0.01285 0.01293 0.0129 0.01292
0 0.01133 0.01146 0.01147 0.01173 0.01123 0.01165 0.01338 0.01214 0.01258 0.01266 0.01263 0.01267
Seward
Terrace
Vancouver
Ashford
Azalea
McBride
Jackson
Crested Butte
Kamchatka
Storslett
Raattama
Oulanka
Korpilahti

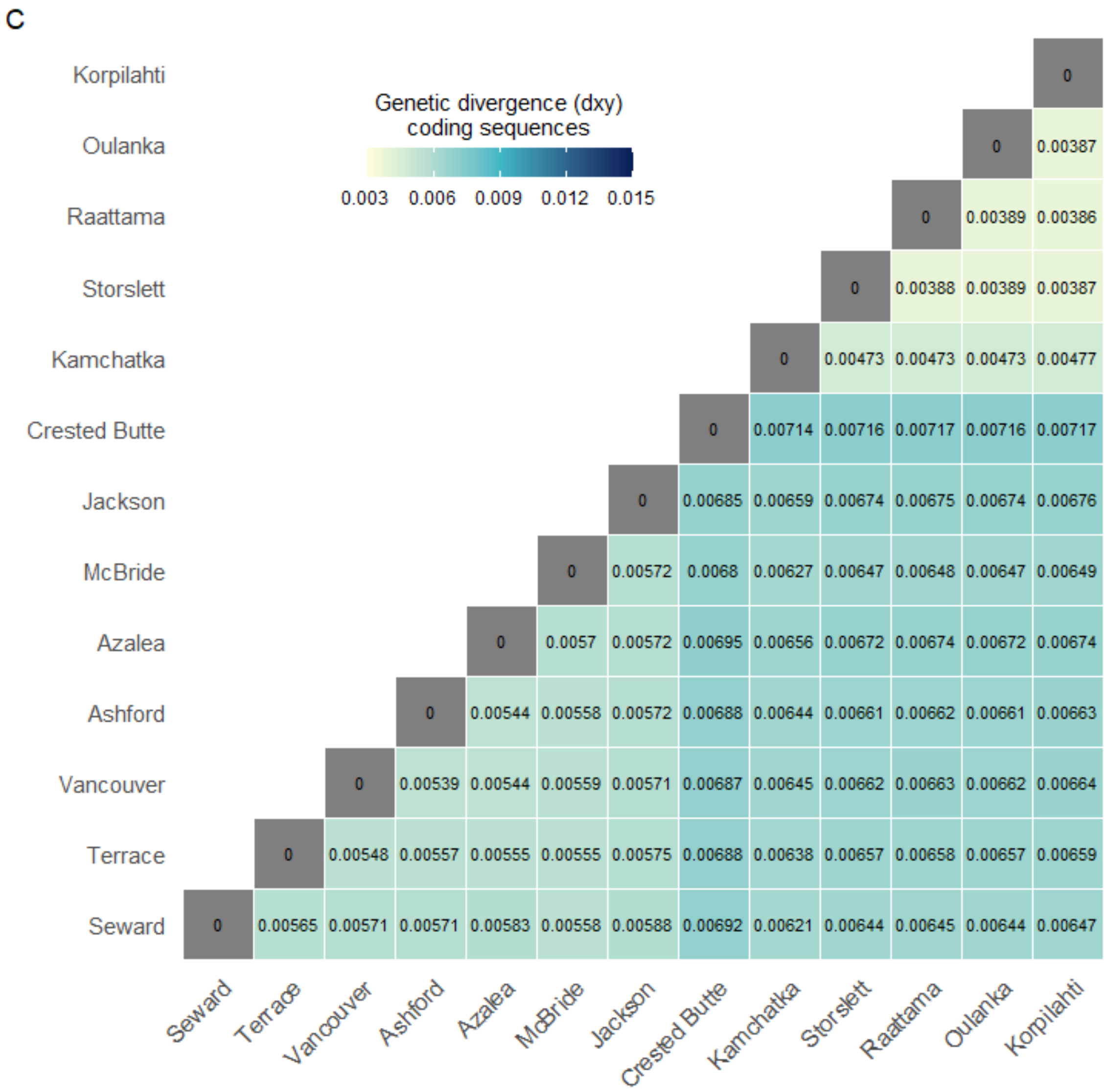

Figure S4. Mean genetic divergence ($d_{xy}$) of all population pairs for (A) intergenic, (B), intronic and (C) coding sequences.

A

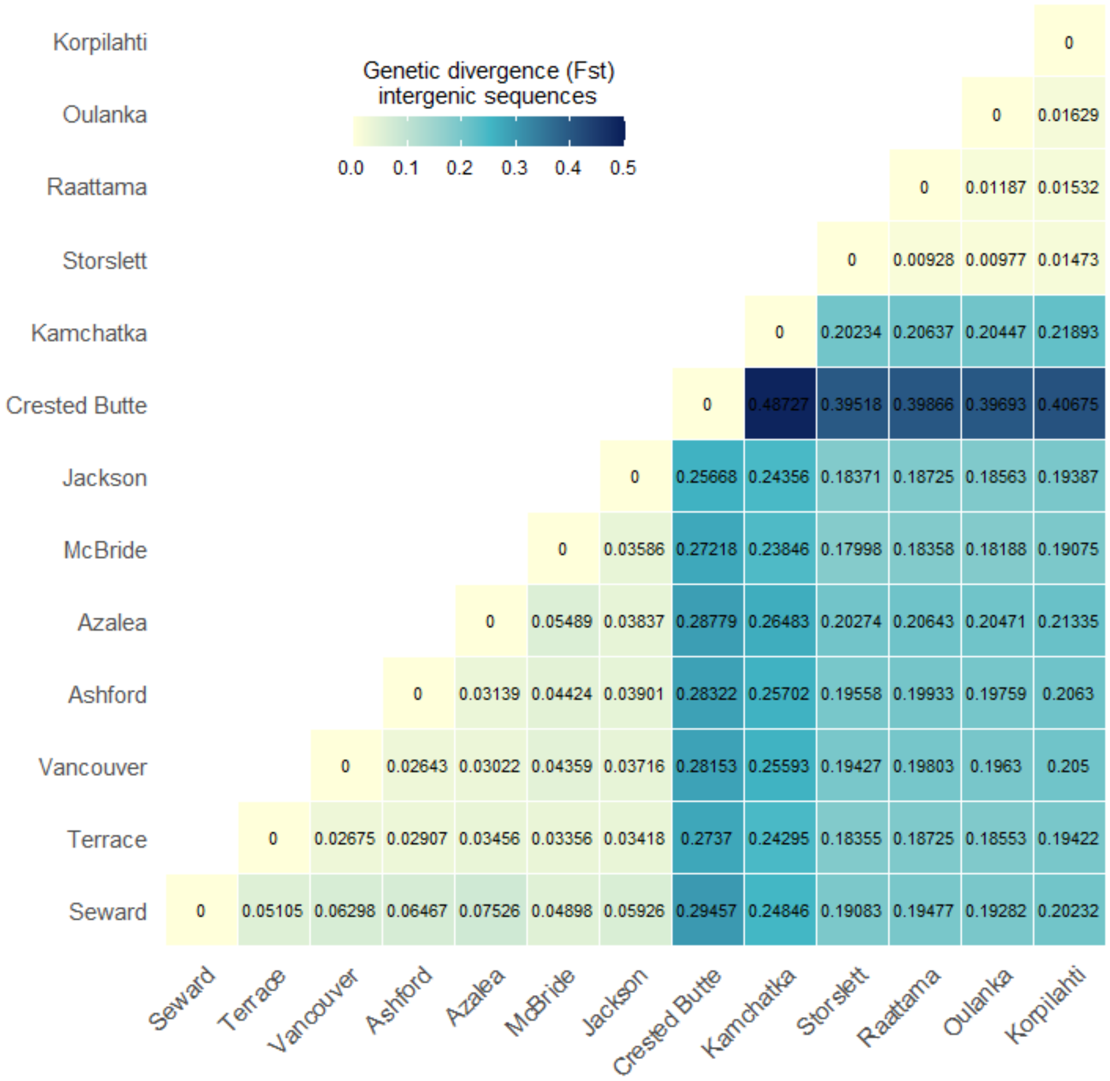
Genetic divergence (Fst)
intergenic sequences
0.0 0.1 0.2 0.3 0.4 0.5
Korpilahti 0
Oulanka 0 0.01629
Raattama 0 0.01187 0.01532
Storslett 0 0.00928 0.00977 0.01473
Kamchatka 0 0.20234 0.20637 0.20447 0.21893
Crested Butte 0 0.48727 0.39518 0.39866 0.39693 0.40675
Jackson 0 0.25668 0.24356 0.18371 0.18725 0.18563 0.19387
McBride 0 0.03586 0.27218 0.23846 0.17998 0.18358 0.18188 0.19075
Azalea 0 0.05489 0.03837 0.28779 0.26483 0.20274 0.20643 0.20471 0.21335
Ashford 0 0.03139 0.04424 0.03901 0.28322 0.25702 0.19558 0.19933 0.19759 0.2063
Vancouver 0 0.02643 0.03022 0.04359 0.03716 0.28153 0.25593 0.19427 0.19803 0.1963 0.205
Terrace 0 0.02675 0.02907 0.03456 0.03356 0.03418 0.2737 0.24295 0.18355 0.18725 0.18553 0.19422
Seward 0 0.05105 0.06298 0.06467 0.07526 0.04898 0.05926 0.29457 0.24846 0.19083 0.19477 0.19282 0.20232
Seward Terrace Vancouver Ashford Azalea McBride Jackson Crested Butte Kamchatka Storslett Raattama Oulanka Korpilahti

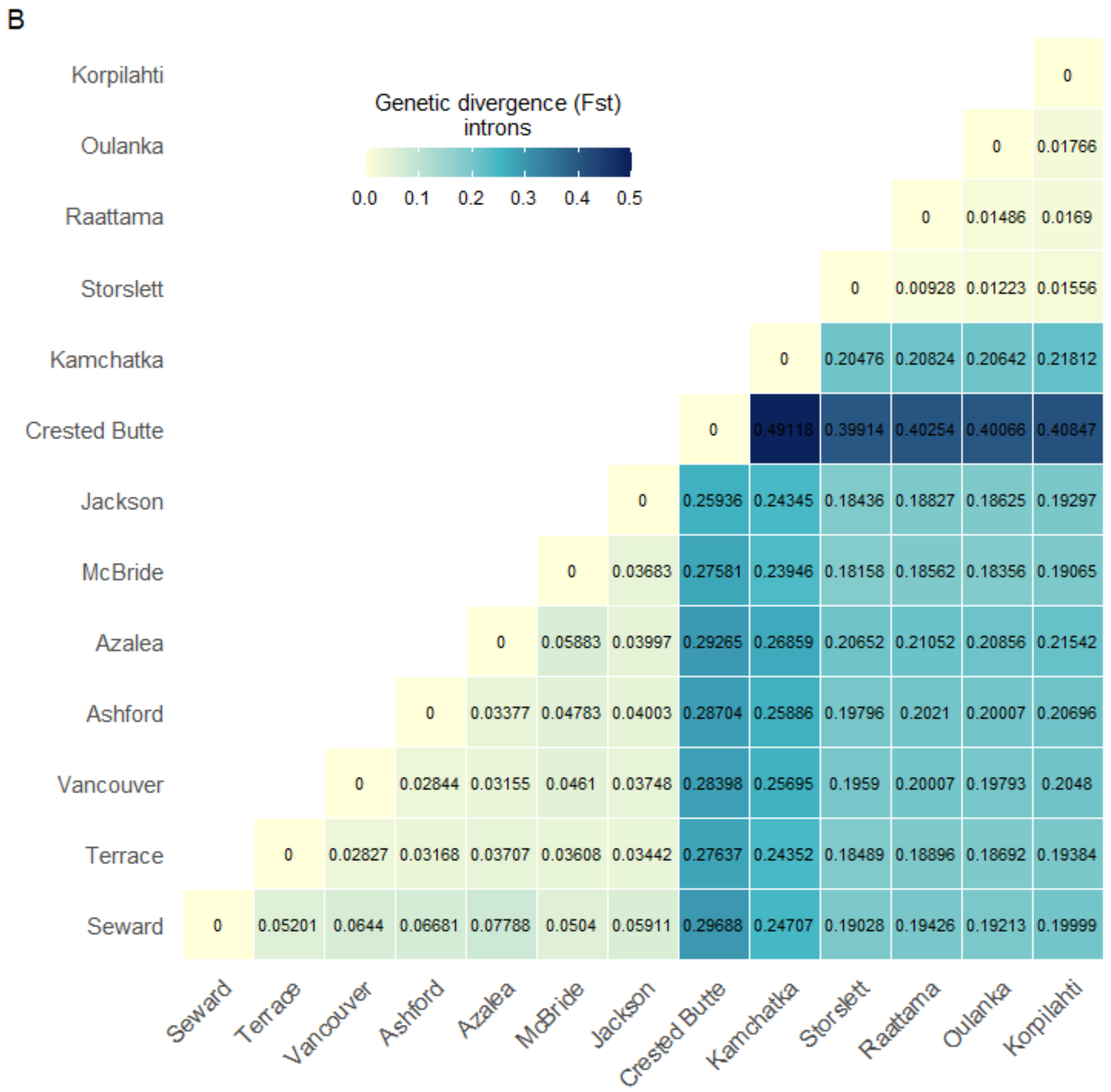
B
Genetic divergence (Fst)
introns
0.0 0.1 0.2 0.3 0.4 0.5
Korpilahti 0
Oulanka 0 0.01766
Raattama 0 0.01486 0.0169
Storslett 0 0.00928 0.01223 0.01556
Kamchatka 0 0.20476 0.20824 0.20642 0.21812
Crested Butte 0 0.49118 0.39914 0.40254 0.40066 0.40847
Jackson 0 0.25936 0.24345 0.18436 0.18827 0.18625 0.19297
McBride 0 0.03683 0.27581 0.23946 0.18158 0.18562 0.18356 0.19065
Azalea 0 0.05883 0.03997 0.29265 0.26859 0.20652 0.21052 0.20856 0.21542
Ashford 0 0.03377 0.04783 0.04003 0.28704 0.25886 0.19796 0.2021 0.20007 0.20696
Vancouver 0 0.02844 0.03155 0.0461 0.03748 0.28398 0.25695 0.1959 0.20007 0.19793 0.2048
Terrace 0 0.02827 0.03168 0.03707 0.03608 0.03442 0.27637 0.24352 0.18489 0.18896 0.18692 0.19384
Seward 0 0.05201 0.0644 0.06681 0.07788 0.0504 0.05911 0.29688 0.24707 0.19028 0.19426 0.19213 0.19999
Seward Terrace Vancouver Ashford Azalea McBride Jackson Crested Butte Kamchatka Storslett Raattama Oulanka Korpilahti

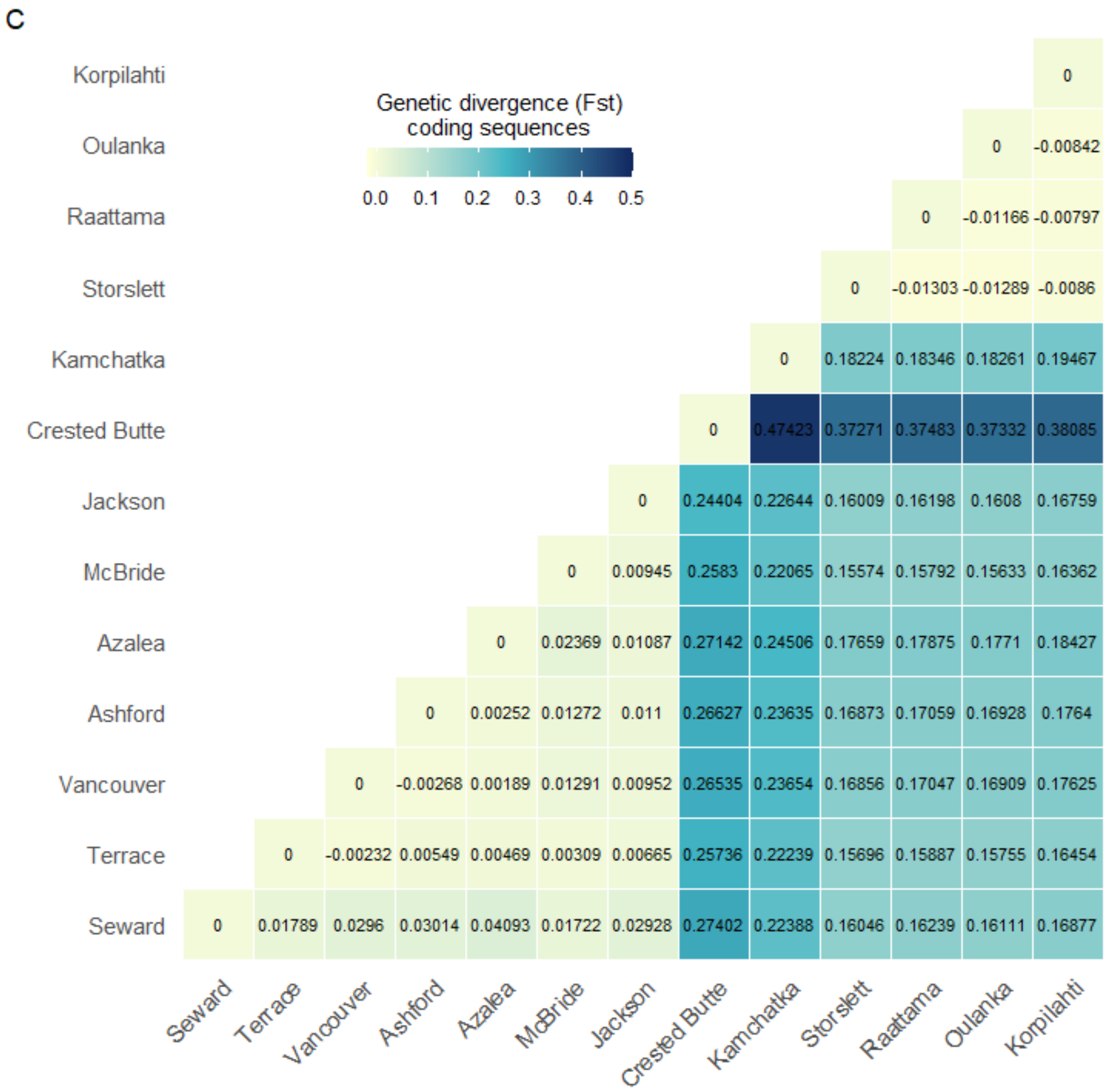

Figure S5. Mean genetic divergence ($F_{st}$) of all population pairs for (A) intergenic, (B), intronic and (C) coding sequences.

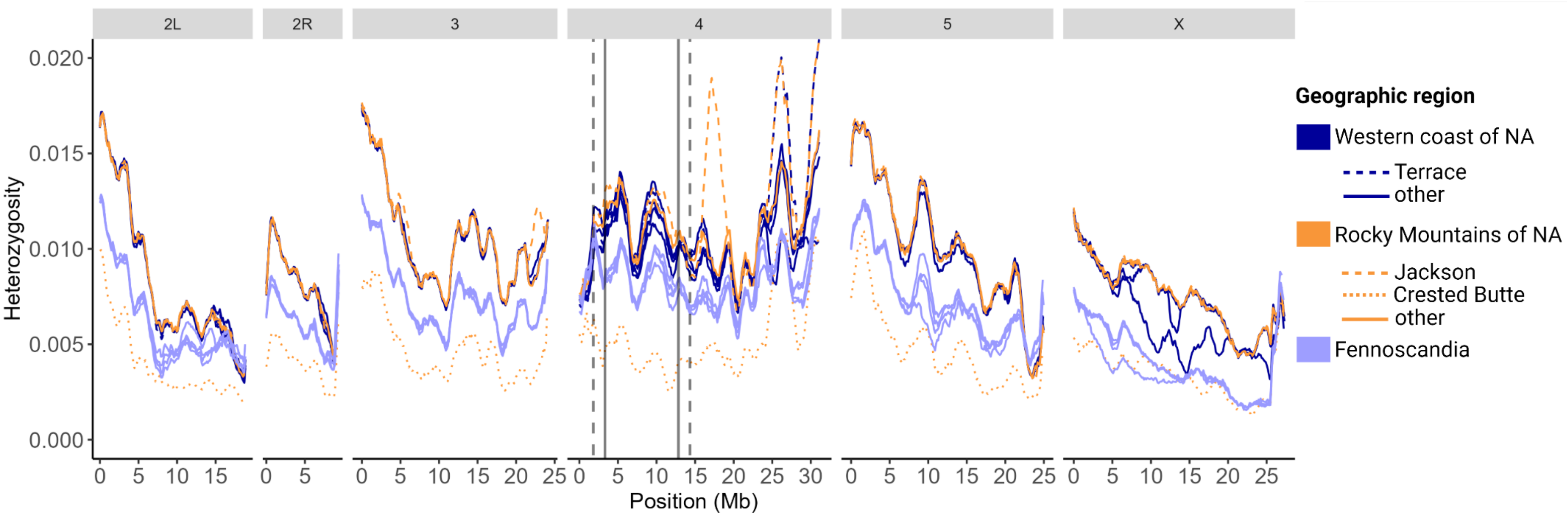

Figure S6. Genome-wide heterozygosity (*H*) of *D. montana* populations originating from the western coast and Rocky Mountains of North America (NA), and Fennoscandia. Populations are highlighted with dashed lines whenever their genetic diversity strongly deviates from that of the other populations of the same geographic region. The inversion breakpoints are marked with vertical solid lines. Since recombination is often suppressed beyond the inverted region into the colinear region, 1.5Mb buffer region was added at both ends of the inversion and indicated with vertical dashed lines.

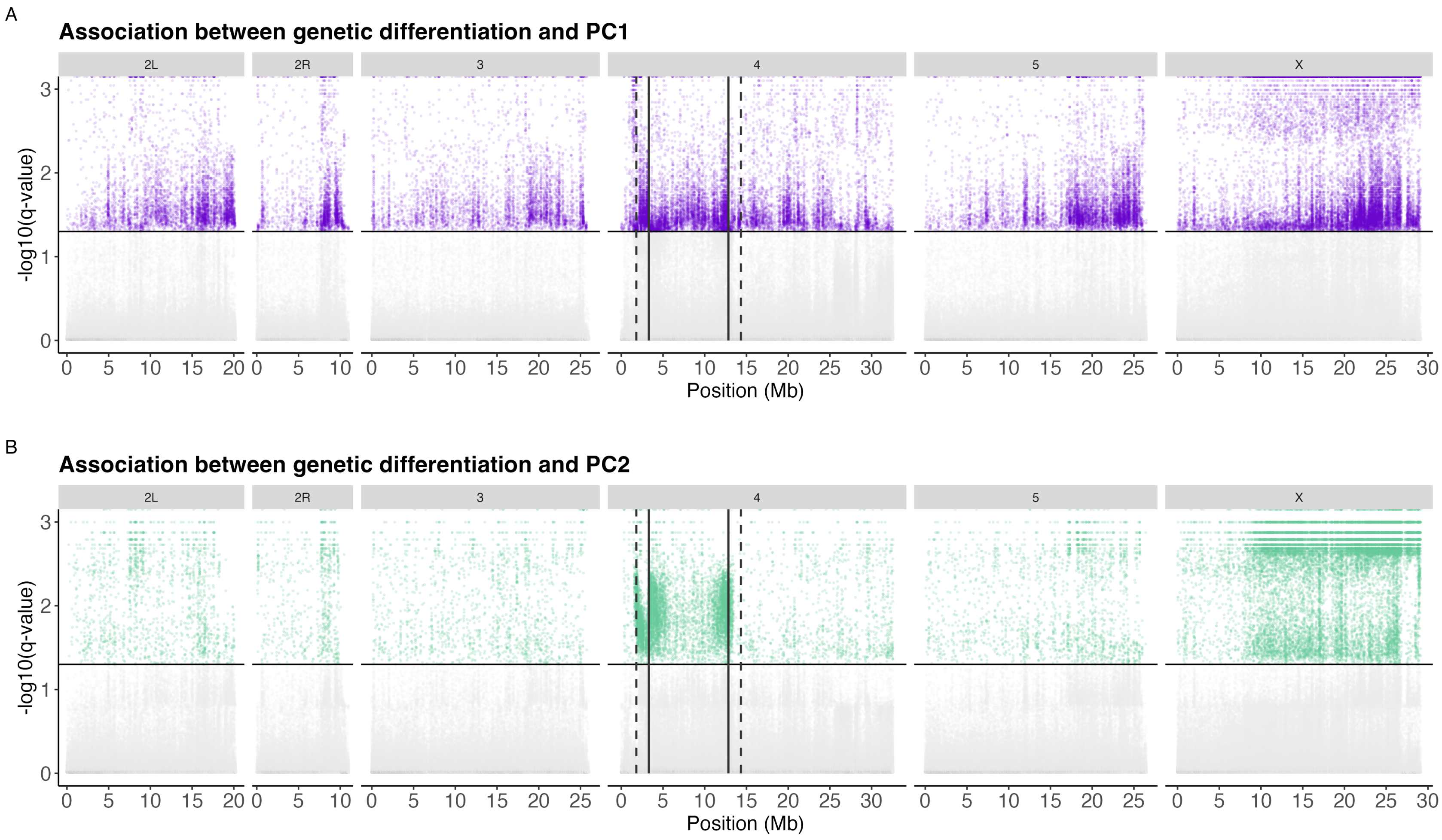
A
Association between genetic differentiation and PC1
2L
2R
3
4
5
X
-log10(q-value)
Position (Mb)
B
Association between genetic differentiation and PC2
2L
2R
3
4
5
X
-log10(q-value)
Position (Mb)

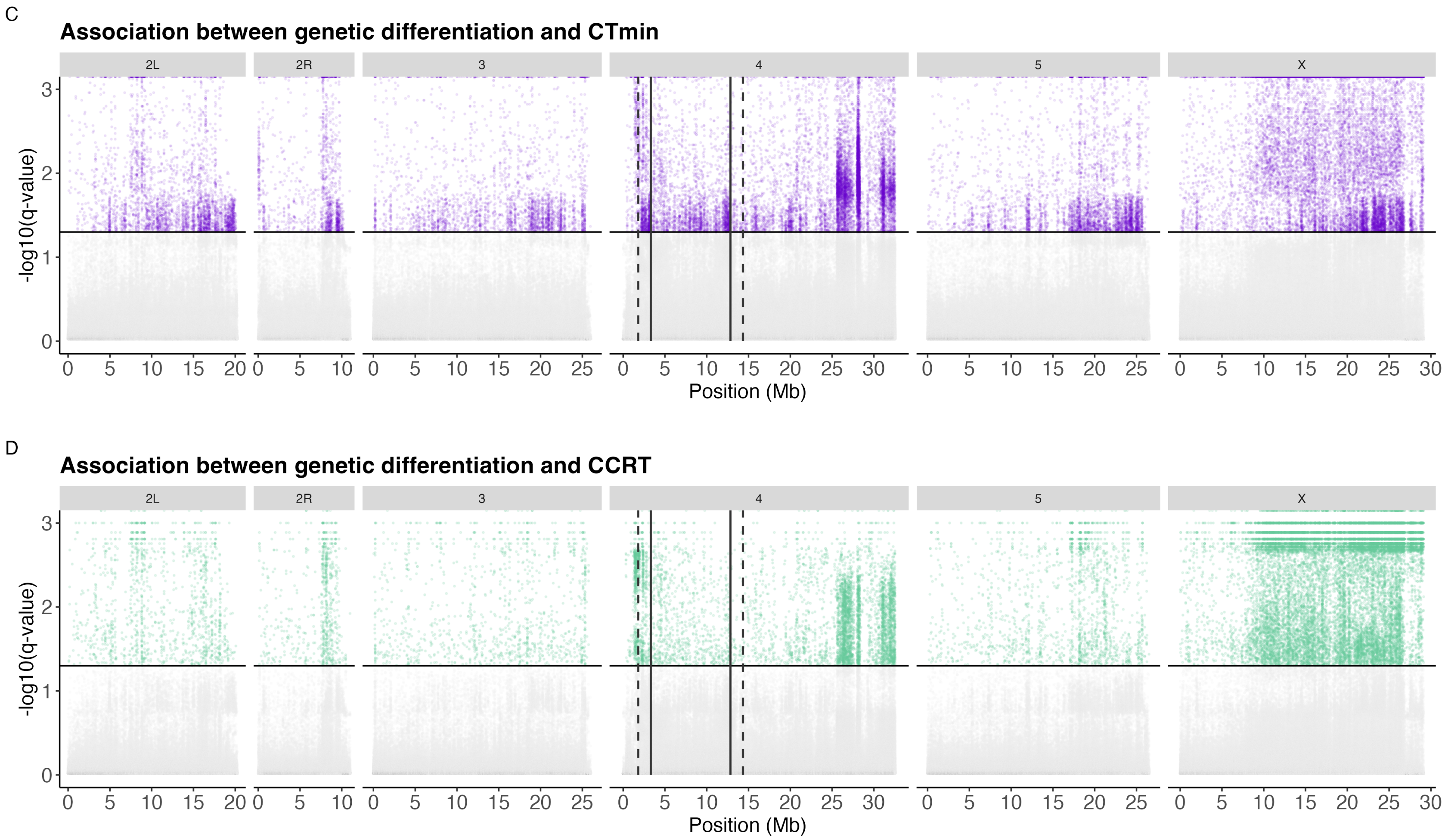

Figure S7. Genotype-environment and genotype-phenotype association analysis (BayeScEnv). Points above the horizontal line indicate significant SNPs, describing the association between genetic differentiation and (A) PC1, (B) PC2, (C) $CT_{min}$ and (D) CCRT. q-values for the g parameter of two chains (Fig. S8-S11) were used to control the false positive rate at 0.05, i.e. SNPs with q-values>1.3, to obtain the final candidate SNPs. The solid vertical lines represent the inversion breakpoints, while the dashed vertical lines mark a conservative estimate of 1.5Mb beyond the breakpoints, where recombination is expected to remain reduced.

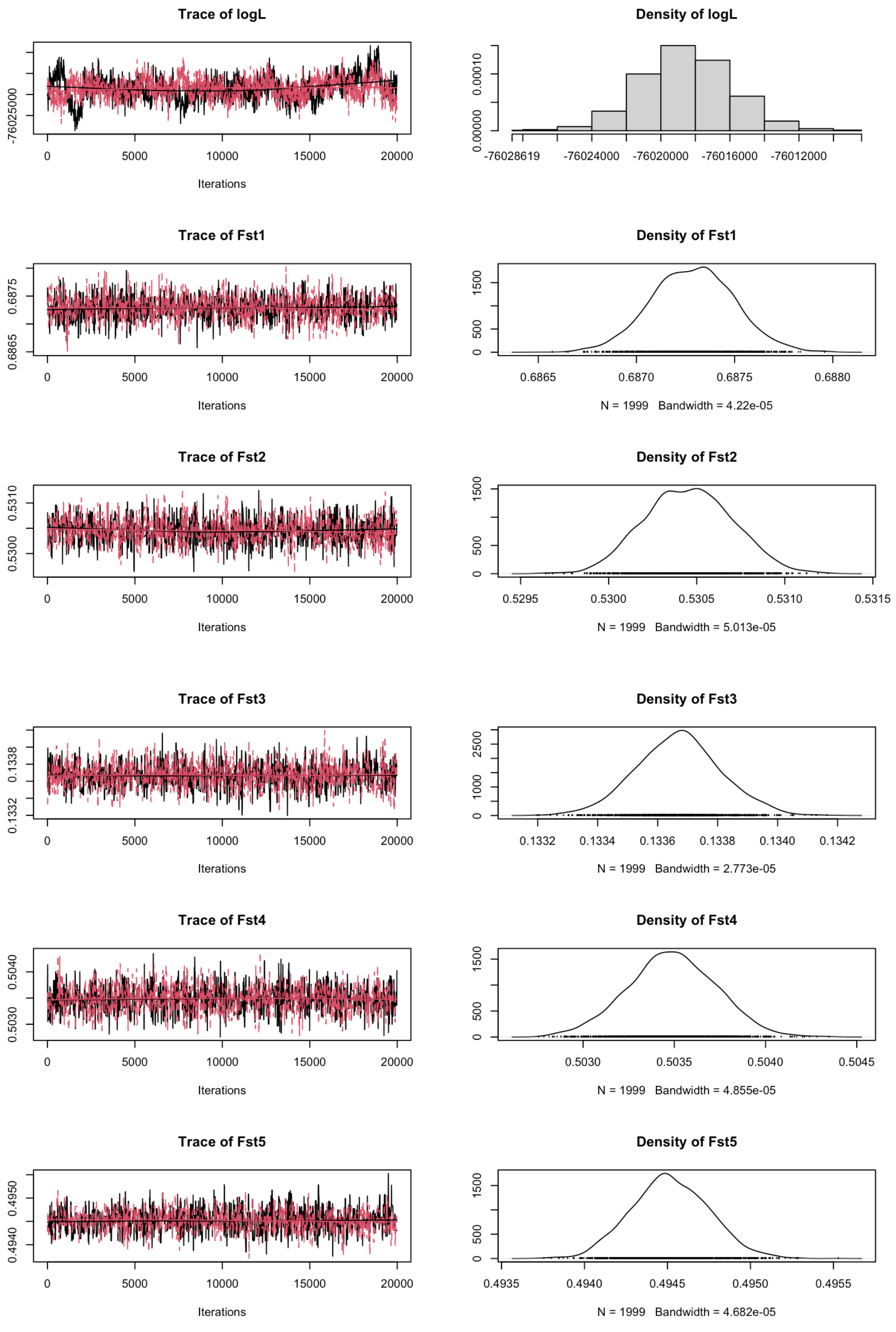
Trace of logL
-76025000
0
5000
10000
15000
20000
Iterations
Density of logL
0.00000
0.00010
-76028619
-76024000
-76020000
-76016000
-76012000
Trace of Fst1
0.6865
0.6875
Iterations
Density of Fst1
0
500
1500
0.6865
0.6870
0.6875
0.6880
N = 1999 Bandwidth = 4.22e-05
Trace of Fst2
0.5300
0.5310
Iterations
Density of Fst2
0.5295
0.5300
0.5305
0.5310
0.5315
N = 1999 Bandwidth = 5.013e-05
Trace of Fst3
0.1332
0.1338
Iterations
Density of Fst3
1000
2500
0.1332
0.1334
0.1336
0.1338
0.1340
0.1342
N = 1999 Bandwidth = 2.773e-05
Trace of Fst4
0.5030
0.5040
Iterations
Density of Fst4
0.5030
0.5035
0.5040
0.5045
N = 1999 Bandwidth = 4.855e-05
Trace of Fst5
0.4940
0.4950
Iterations
Density of Fst5
0.4935
0.4940
0.4945
0.4950
0.4955
N = 1999 Bandwidth = 4.682e-05

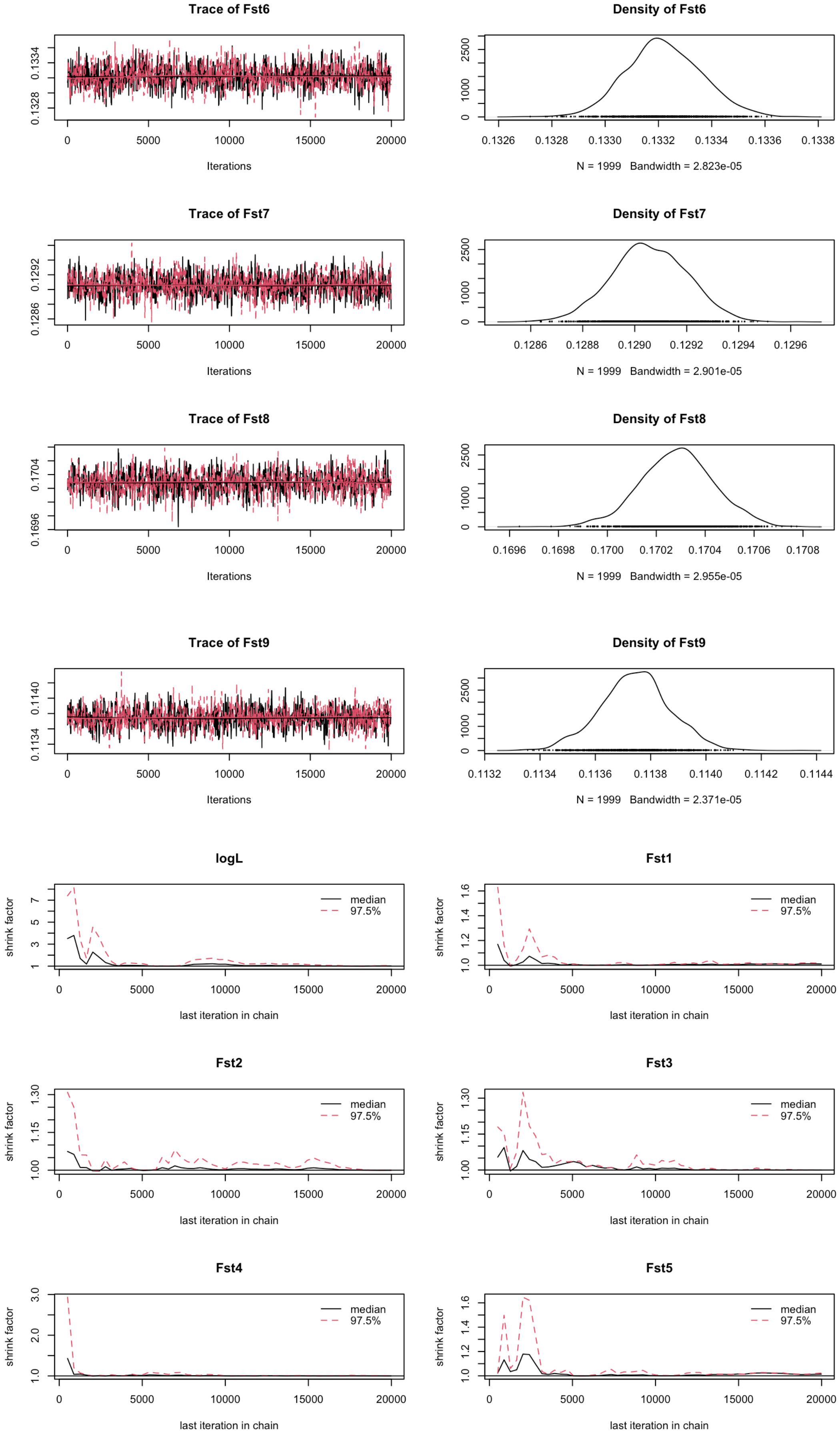
Trace of Fst6
Iterations
Density of Fst6
N = 1999 Bandwidth = 2.823e-05
Trace of Fst7
Iterations
Density of Fst7
N = 1999 Bandwidth = 2.901e-05
Trace of Fst8
Iterations
Density of Fst8
N = 1999 Bandwidth = 2.955e-05
Trace of Fst9
Iterations
Density of Fst9
N = 1999 Bandwidth = 2.371e-05
logL
shrink factor
median
97.5%
last iteration in chain
Fst1
shrink factor
median
97.5%
last iteration in chain
Fst2
shrink factor
median
97.5%
last iteration in chain
Fst3
shrink factor
median
97.5%
last iteration in chain
Fst4
shrink factor
median
97.5%
last iteration in chain
Fst5
shrink factor
median
97.5%
last iteration in chain

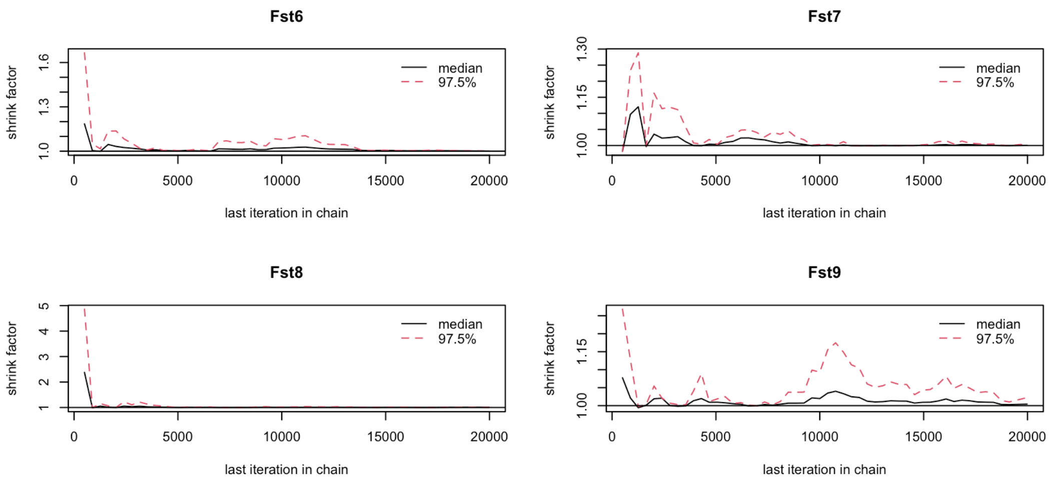

Figure S8. Trace and diagnostic Gelman plots for BayeScEnv MCMC chains with PC1 as the environmental variable. Trace plots show parameter values across iterations of MCMC chains. Gelman plots show the Gelman-Rubin shrink factors that summarise variation in parameter estimates across the two chains that were run.

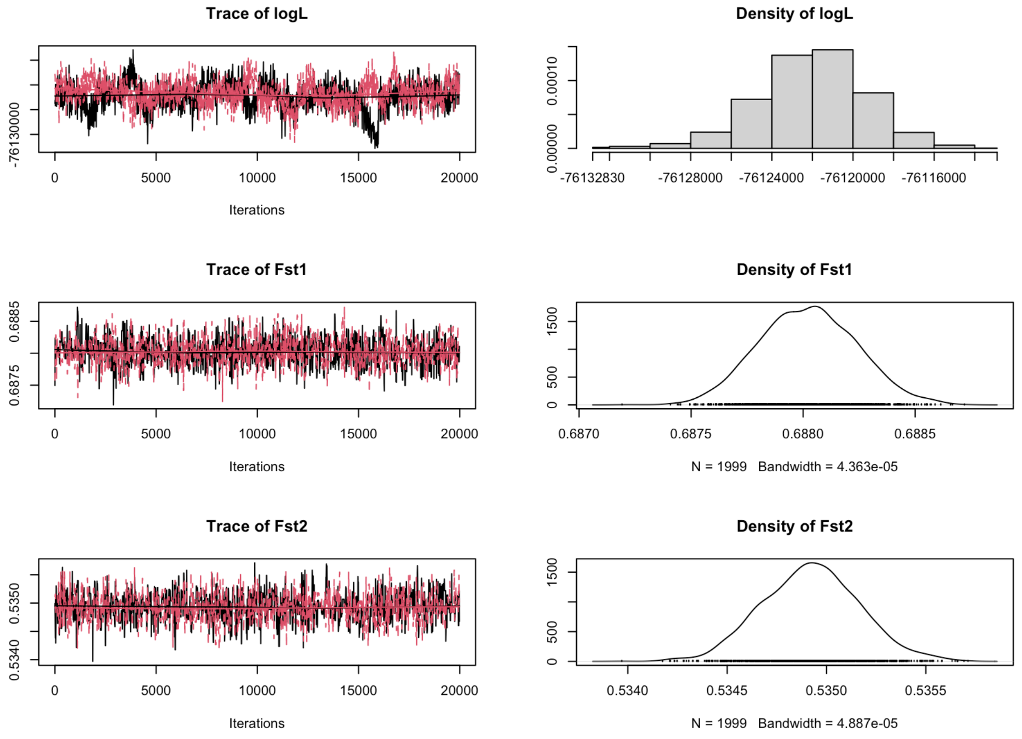

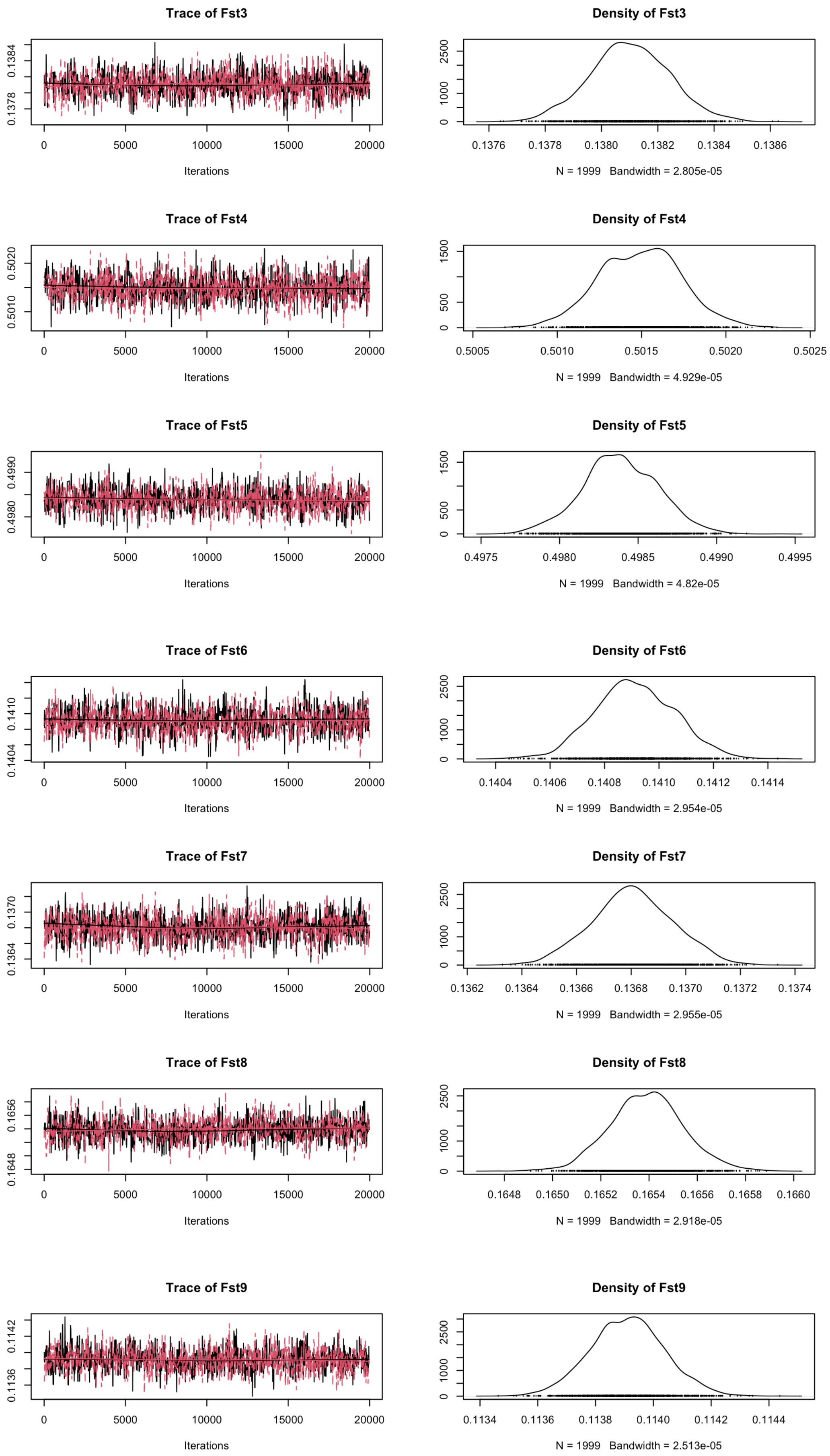
Trace of Fst3
Iterations
Density of Fst3
N = 1999 Bandwidth = 2.805e-05
Trace of Fst4
Iterations
Density of Fst4
N = 1999 Bandwidth = 4.929e-05
Trace of Fst5
Iterations
Density of Fst5
N = 1999 Bandwidth = 4.82e-05
Trace of Fst6
Iterations
Density of Fst6
N = 1999 Bandwidth = 2.954e-05
Trace of Fst7
Iterations
Density of Fst7
N = 1999 Bandwidth = 2.955e-05
Trace of Fst8
Iterations
Density of Fst8
N = 1999 Bandwidth = 2.918e-05
Trace of Fst9
Iterations
Density of Fst9
N = 1999 Bandwidth = 2.513e-05

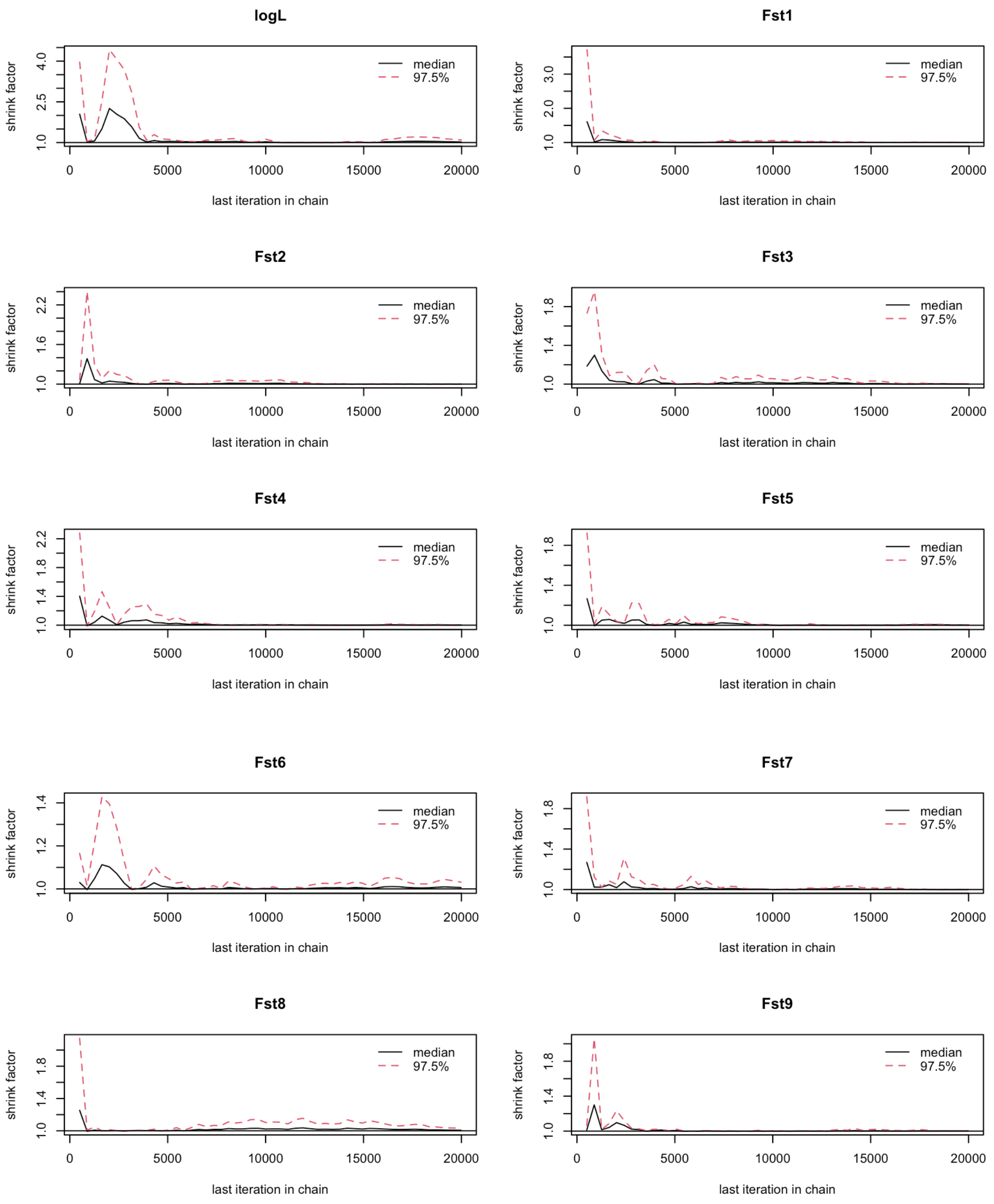

Figure S9. Trace and diagnostic Gelman plots for BayeScEnv MCMC chains with PC2 as the environmental variable. Trace plots show parameter values across iterations of MCMC chains. Gelman plots show the Gelman-Rubin shrink factors that summarise variation in parameter estimates across the two chains that were run.

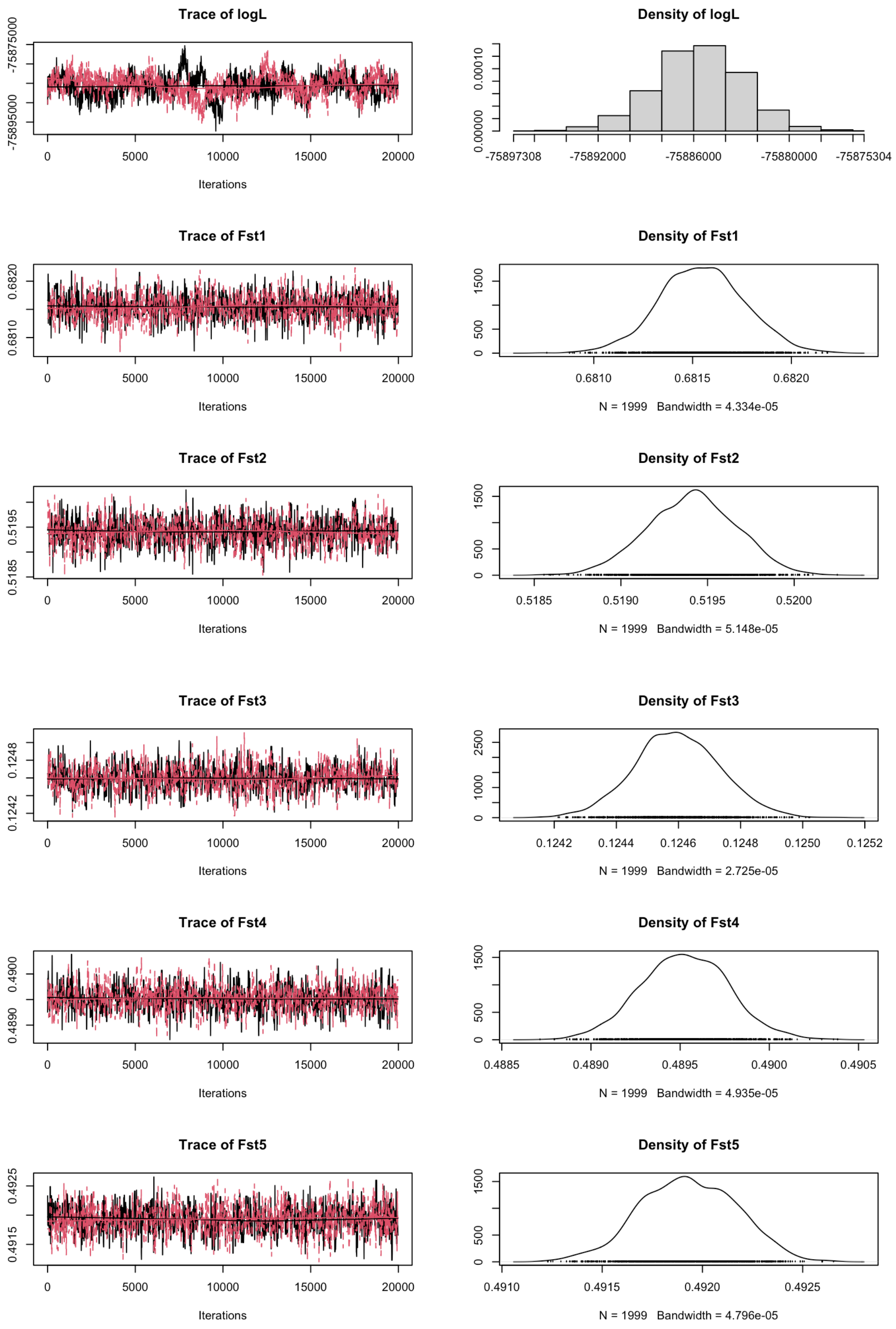
Trace of logL
-75875000
-75895000
0 5000 10000 15000 20000
Iterations
Density of logL
0.00010
0.00000
-75897308 -75892000 -75886000 -75880000 -75875304
Trace of Fst1
0.6820
0.6810
0 5000 10000 15000 20000
Iterations
Density of Fst1
1500
500
0
0.6810 0.6815 0.6820
N = 1999 Bandwidth = 4.334e-05
Trace of Fst2
0.5195
0.5185
0 5000 10000 15000 20000
Iterations
Density of Fst2
1500
500
0
0.5185 0.5190 0.5195 0.5200
N = 1999 Bandwidth = 5.148e-05
Trace of Fst3
0.1248
0.1242
0 5000 10000 15000 20000
Iterations
Density of Fst3
2500
1000
0
0.1242 0.1244 0.1246 0.1248 0.1250 0.1252
N = 1999 Bandwidth = 2.725e-05
Trace of Fst4
0.4900
0.4890
0 5000 10000 15000 20000
Iterations
Density of Fst4
1500
500
0
0.4885 0.4890 0.4895 0.4900 0.4905
N = 1999 Bandwidth = 4.935e-05
Trace of Fst5
0.4925
0.4915
0 5000 10000 15000 20000
Iterations
Density of Fst5
1500
500
0
0.4910 0.4915 0.4920 0.4925
N = 1999 Bandwidth = 4.796e-05

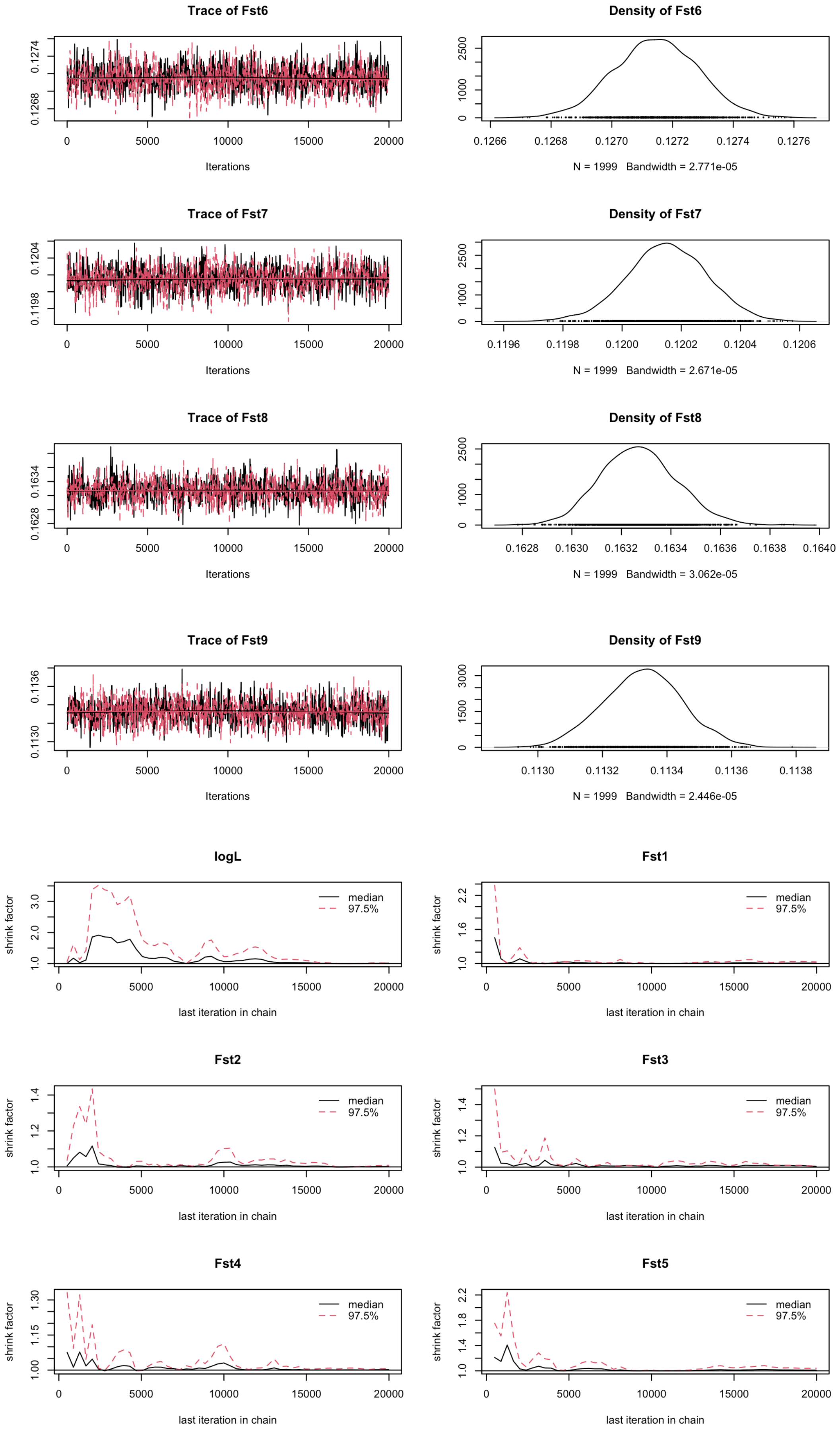
Trace of Fst6
0.1268
0.1274
0
5000
10000
15000
20000
Iterations
Density of Fst6
0
1000
2500
0.1266
0.1268
0.1270
0.1272
0.1274
0.1276
N = 1999 Bandwidth = 2.771e-05
Trace of Fst7
0.1198
0.1204
0
5000
10000
15000
20000
Iterations
Density of Fst7
0
1000
2500
0.1196
0.1198
0.1200
0.1202
0.1204
0.1206
N = 1999 Bandwidth = 2.671e-05
Trace of Fst8
0.1628
0.1634
0
5000
10000
15000
20000
Iterations
Density of Fst8
0
1000
2500
0.1628
0.1630
0.1632
0.1634
0.1636
0.1638
0.1640
N = 1999 Bandwidth = 3.062e-05
Trace of Fst9
0.1130
0.1136
0
5000
10000
15000
20000
Iterations
Density of Fst9
0
1500
3000
0.1130
0.1132
0.1134
0.1136
0.1138
N = 1999 Bandwidth = 2.446e-05
logL
shrink factor
1.0
2.0
3.0
median
97.5%
0
5000
10000
15000
20000
last iteration in chain
Fst1
shrink factor
1.0
1.6
2.2
median
97.5%
0
5000
10000
15000
20000
last iteration in chain
Fst2
shrink factor
1.0
1.2
1.4
median
97.5%
0
5000
10000
15000
20000
last iteration in chain
Fst3
shrink factor
1.0
1.2
1.4
median
97.5%
0
5000
10000
15000
20000
last iteration in chain
Fst4
shrink factor
1.00
1.15
1.30
median
97.5%
0
5000
10000
15000
20000
last iteration in chain
Fst5
shrink factor
1.0
1.4
1.8
2.2
median
97.5%
0
5000
10000
15000
20000
last iteration in chain

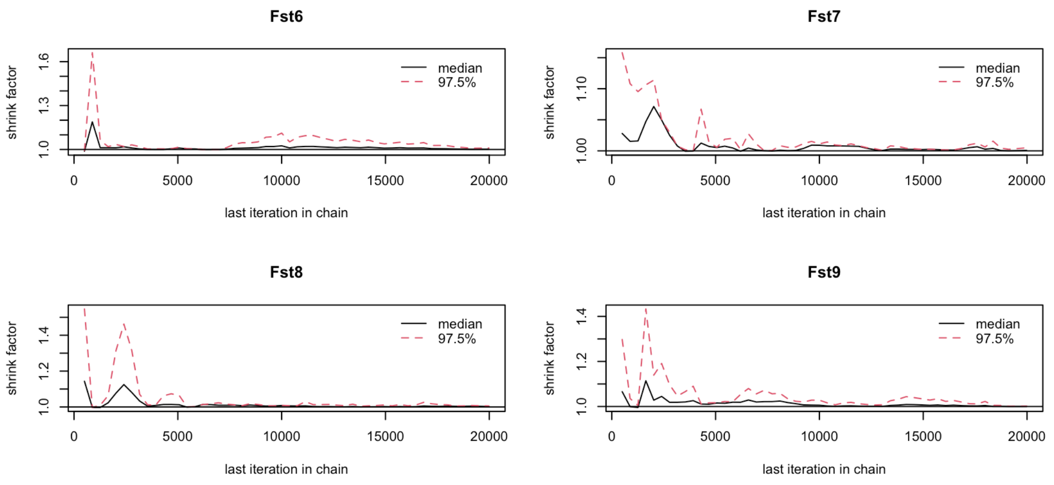

Figure S10. Trace and diagnostic Gelman plots for BayeScEnv MCMC chains with $CT_{min}$ as the environmental variable. Trace plots show parameter values across iterations of MCMC chains. Gelman plots show the Gelman-Rubin shrink factors that summarise variation in parameter estimates across the two chains that were run.

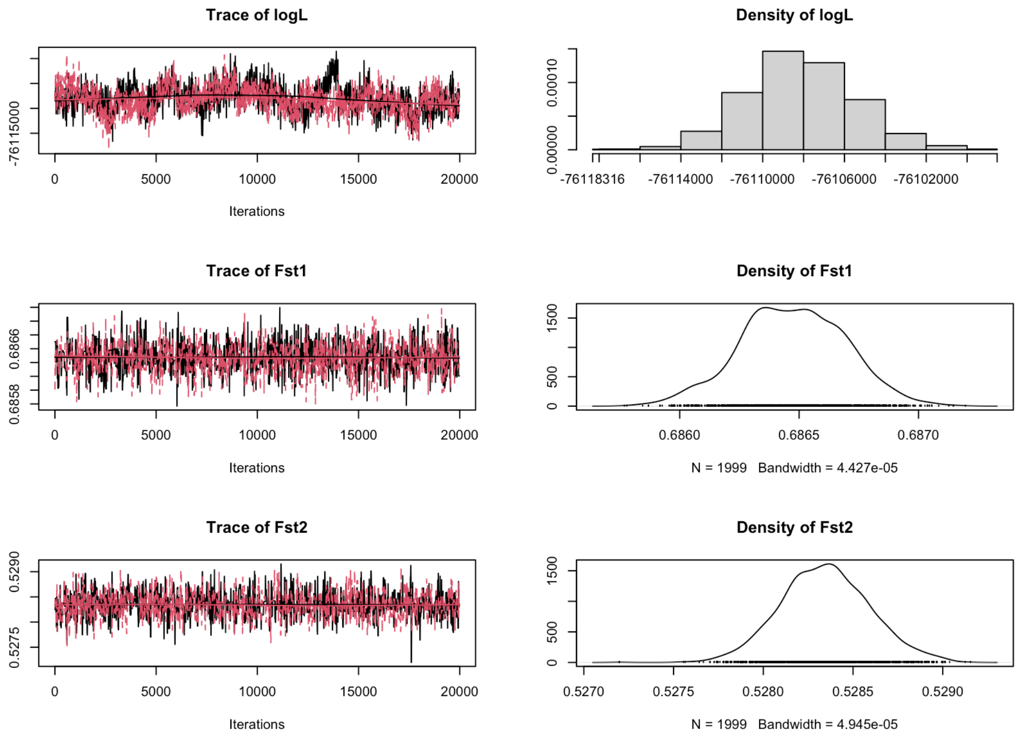

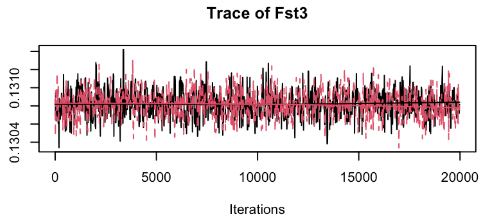
Trace of Fst3
0.1304
0.1310
0 5000 10000 15000 20000
Iterations

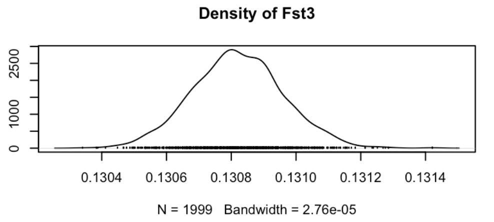
Density of Fst3
0
1000
2500
0.1304 0.1306 0.1308 0.1310 0.1312 0.1314
N = 1999 Bandwidth = 2.76e-05

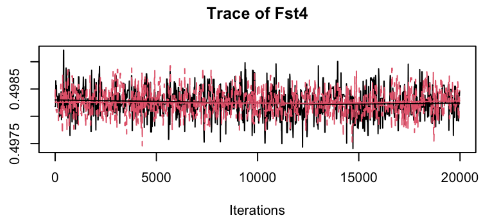
Trace of Fst4
0.4975
0.4985
0 5000 10000 15000 20000
Iterations

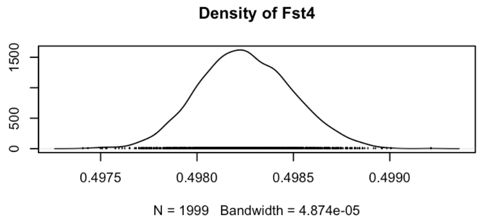
Density of Fst4
0
500
1500
0.4975 0.4980 0.4985 0.4990
N = 1999 Bandwidth = 4.874e-05

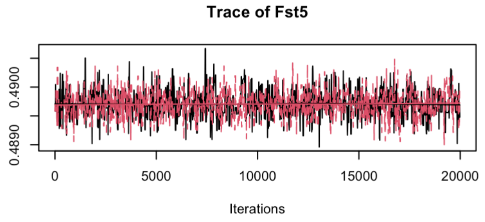
Trace of Fst5
0.4890
0.4900
0 5000 10000 15000 20000
Iterations

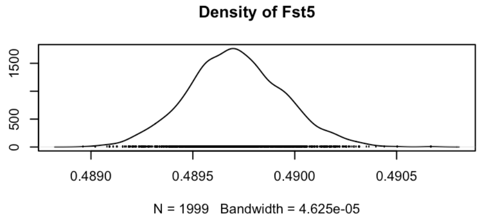
Density of Fst5
0
500
1500
0.4890 0.4895 0.4900 0.4905
N = 1999 Bandwidth = 4.625e-05

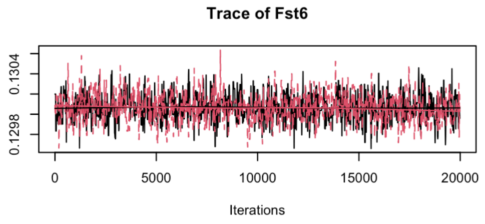
Trace of Fst6
0.1298
0.1304
0 5000 10000 15000 20000
Iterations

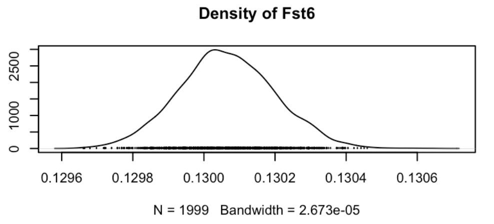
Density of Fst6
0
1000
2500
0.1296 0.1298 0.1300 0.1302 0.1304 0.1306
N = 1999 Bandwidth = 2.673e-05

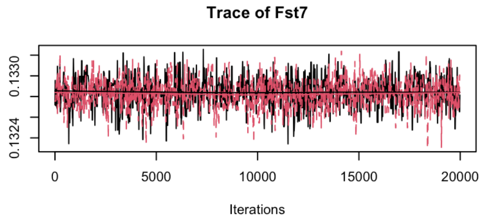
Trace of Fst7
0.1324
0.1330
0 5000 10000 15000 20000
Iterations

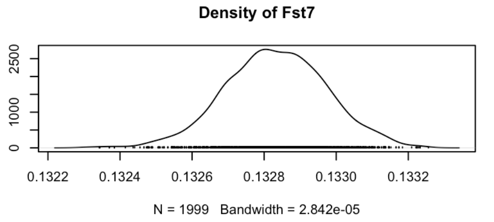
Density of Fst7
0
1000
2500
0.1322 0.1324 0.1326 0.1328 0.1330 0.1332
N = 1999 Bandwidth = 2.842e-05

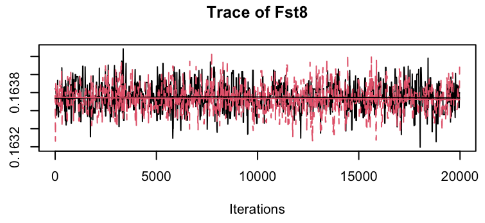
Trace of Fst8
0.1632
0.1638
0 5000 10000 15000 20000
Iterations

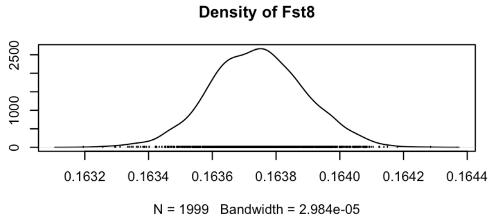
Density of Fst8
0
1000
2500
0.1632 0.1634 0.1636 0.1638 0.1640 0.1642 0.1644
N = 1999 Bandwidth = 2.984e-05

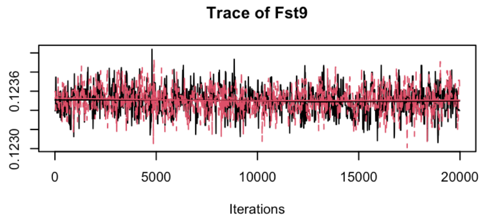
Trace of Fst9
0.1230
0.1236
0 5000 10000 15000 20000
Iterations

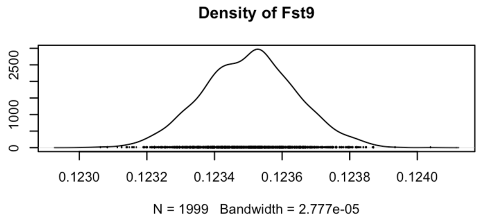
Density of Fst9
0
1000
2500
0.1230 0.1232 0.1234 0.1236 0.1238 0.1240
N = 1999 Bandwidth = 2.777e-05

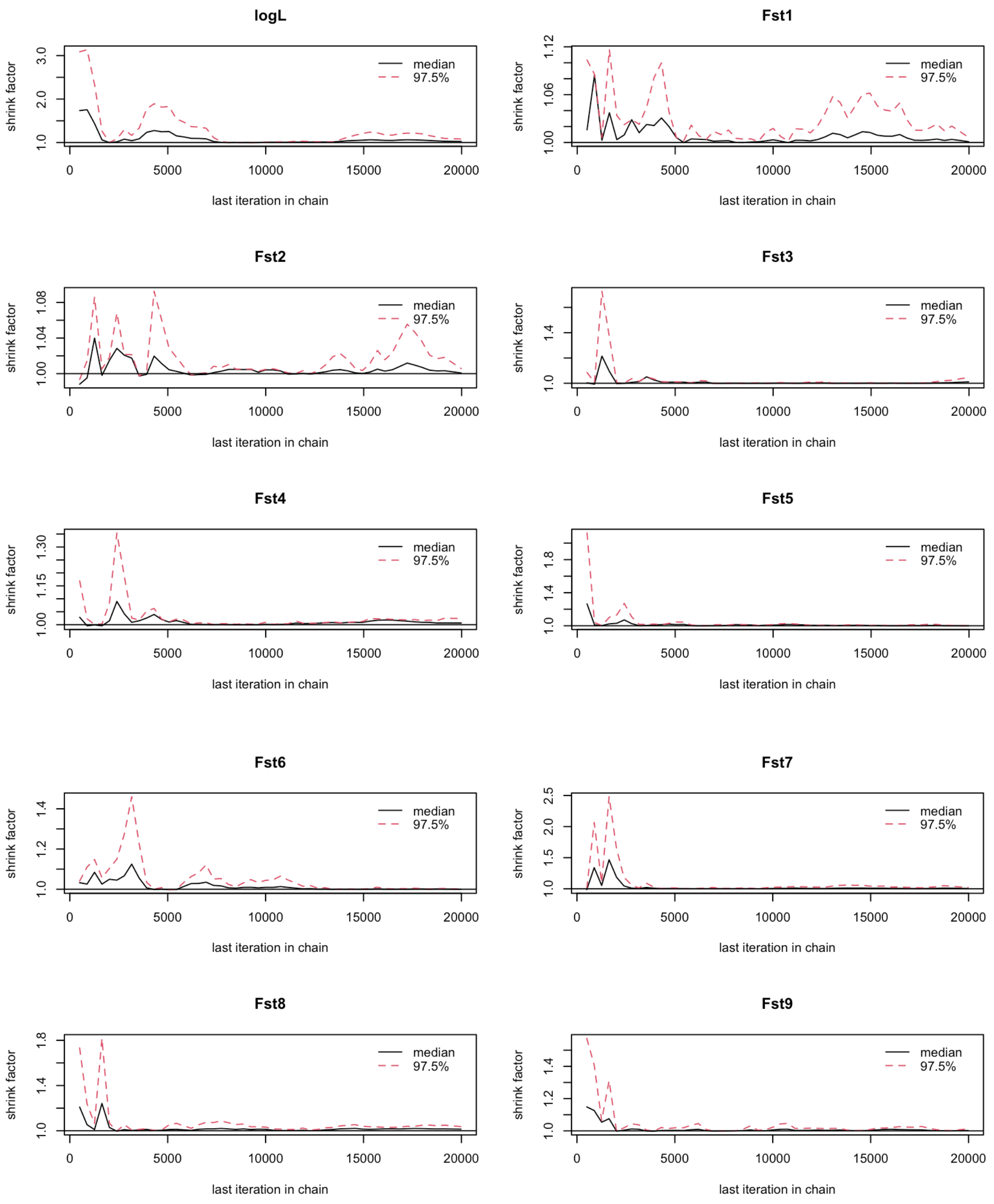

Figure S11. Trace and diagnostic Gelman plots for BayeScEnv MCMC chains with CCRT as the environmental variable. Trace plots show parameter values across iterations of MCMC chains. Gelman plots show the Gelman-Rubin shrink factors that summarise variation in parameter estimates across chains. These plots show data from two chains that were run.

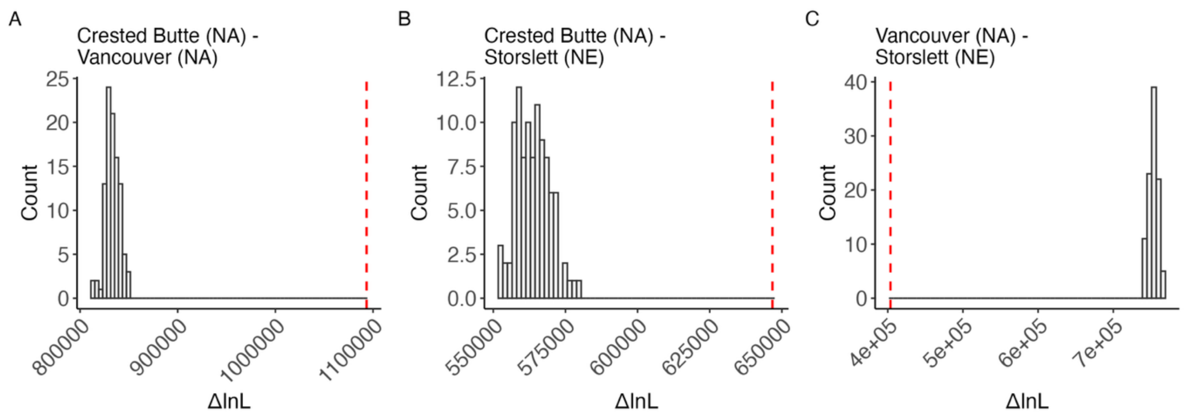

Figure S12. Model comparison was conducted for the following population pairs: (A) Crested Butte (North America, NA) and Vancouver (NA), (B) Crested Butte (NA) and Storslett (Fennoscandia, North Europe, NE), and (C) Vancouver (NA) and Storslett (NE). Datasets simulated under the strict divergence model (DIV) were fitted to both the DIV model and the best-fit isolation with migration (IM) model (see Table S19). The null distribution of ΔlnL was compared to the observed ΔlnL (indicated by vertical dashed lines). If the observed ΔlnL falls on the right side of the null distribution, it indicates that the IM model fits significantly better than the DIV model. The analysis was based on blocks of 256bp and was performed on colinear non-repetitive intergenic regions to minimize the direct effects of selection.

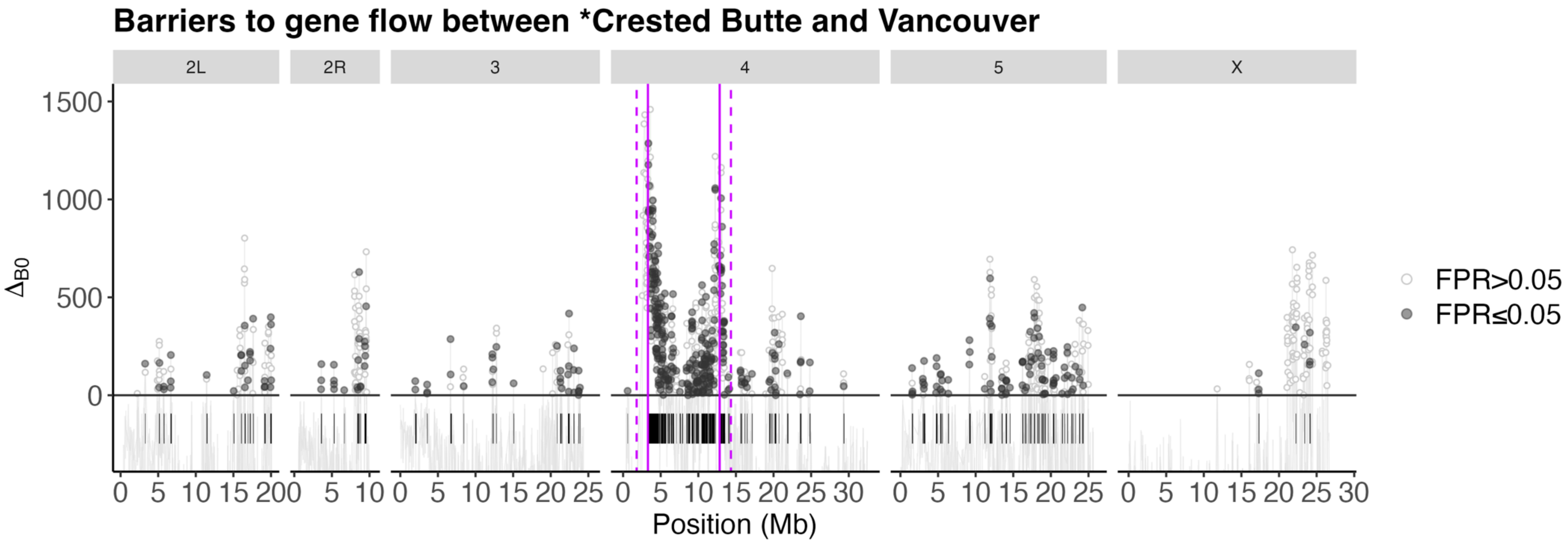

Figure S13. Support for locally reduced $m_e$ between Crested Butte (* inversion fixed) and Vancouver (inversion absent) was measured via positive $\Delta_{B0}$ (data points above the dashed horizontal line) and a false positive rate at 5%. Barrier regions ($\Delta_{B0} > 0$ and FPR≤0.05) are marked with black segments. The solid vertical lines represent the inversion breakpoints, while the dashed vertical lines mark a conservative estimate of 1.5Mb beyond the breakpoints, where recombination is expected to remain reduced.

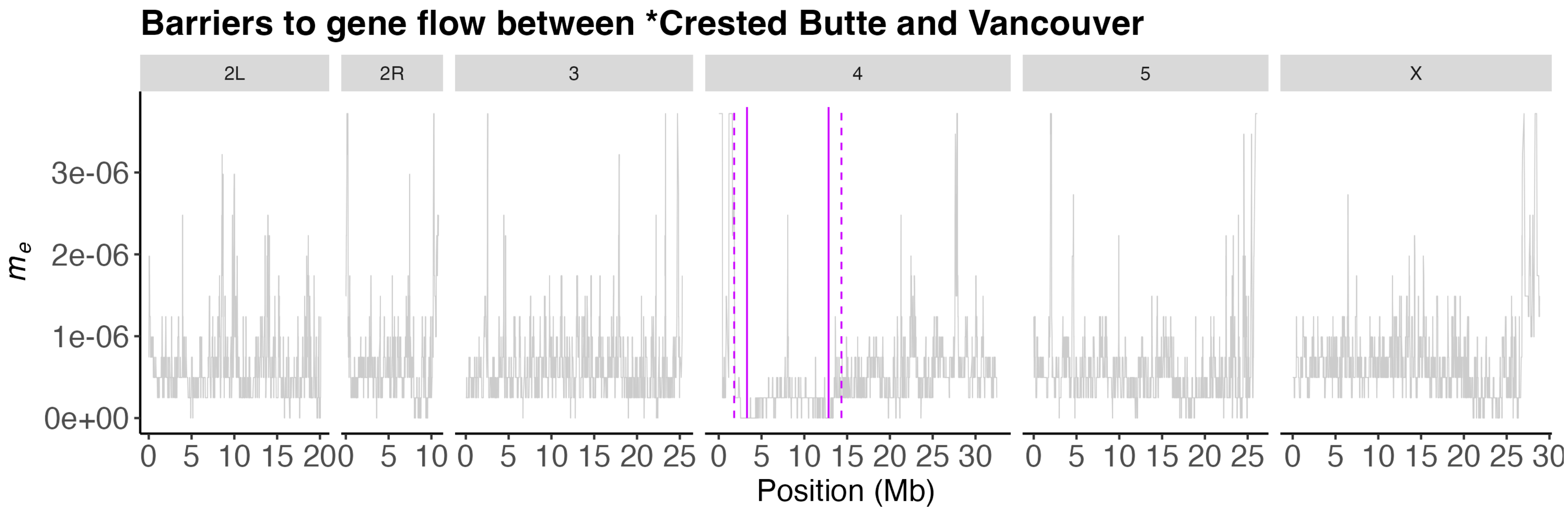

Figure S14. Window-wise variation in the effective migration rate ($m_e$) between Crested Butte (* inversion fixed) and Vancouver (inversion absent). The solid vertical lines represent the inversion breakpoints, while the dashed vertical lines mark a conservative estimate of 1.5Mb beyond the breakpoints, where recombination is expected to remain reduced.

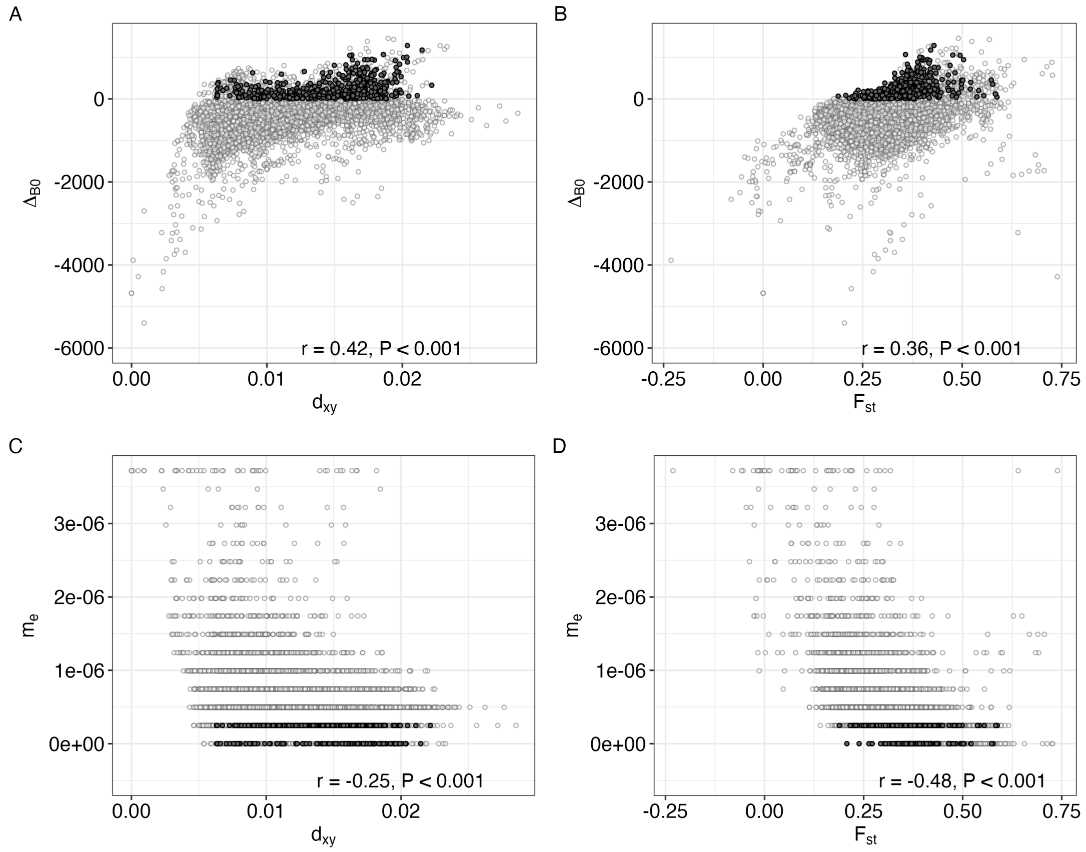

Figure S15. Pearson correlation analysis between (A) $\Delta_{B0}$ and $d_{xy}$, (B) $\Delta_{B0}$ and $F_{st}$, (C) $m_e$ and $d_{xy}$, (D) $m_e$ and $F_{st}$. The strongest barriers to gene flow ($\Delta_{B0}$>0 and FPR ≤0.05, marked with closed circles), only partially overlap with $d_{xy}$ and $F_{st}$ outliers. $m_e$ and $d_{xy}$ or $F_{st}$ are not correlated.

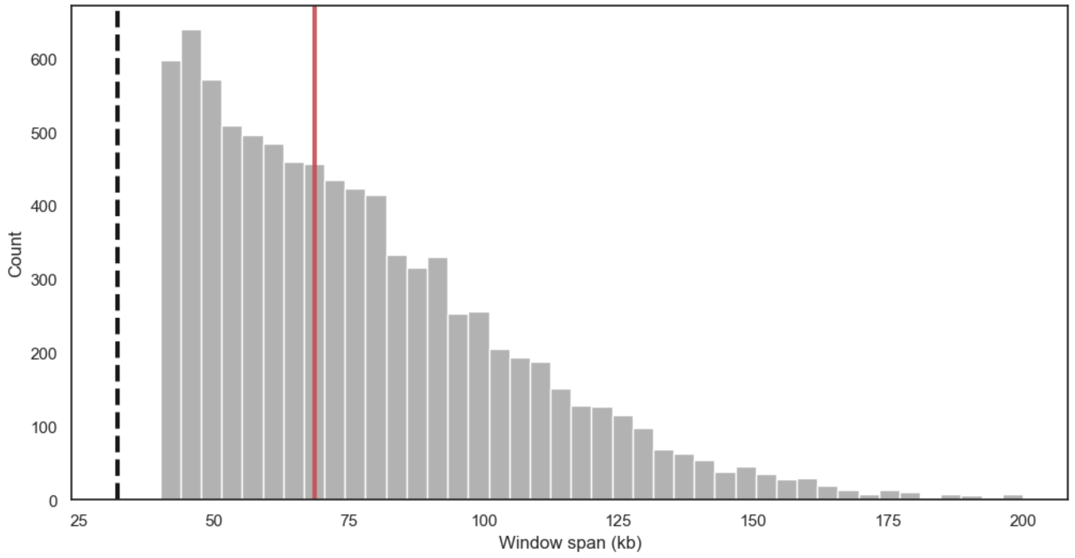

Figure S16. Window span distribution. The genome was analysed in windows of a fixed number of blocks (125 blocks, block size 256bp), which results in a minimum window span of 32 kb (dashed black line). Given that the analysis included only intergenic sequences (repetitive regions excluded), the real window span was usually greater than 32kb (median 68.6kb). Median window span is marked with a solid red line.

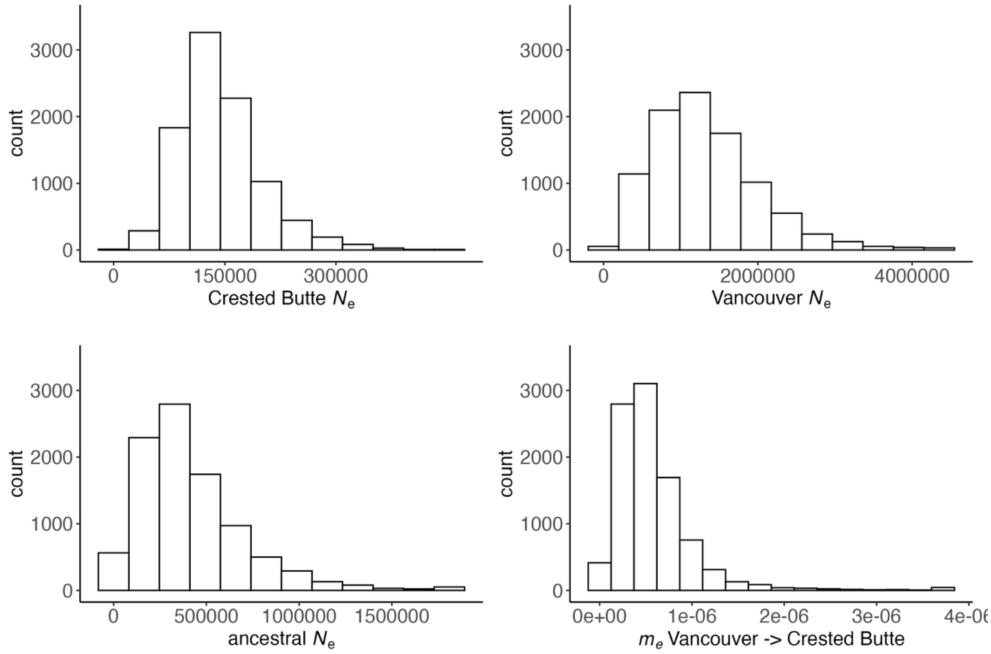

Figure S17. Variation in effective population sizes of Crested Butte (inversion fixed), Vancouver (inversion absent) and their shared ancestral population, and in the migration rate ($m_e$) from Vancouver to Crested Butte (forward in time) across sliding windows. The number of bins in each distribution corresponds to points in the 12 × 12 × 12 × 16 parameter grid used for inference.
